# Taxane induces attenuation of the CXCR2/BCL-2 axis and sensitize prostate cancer to platinum-based treatments

**DOI:** 10.1101/2020.01.20.912824

**Authors:** Vicenç Ruiz de Porras, Xieng C. Wang, Luis Palomero, Mercedes Marin-Aguilera, Alberto Indacochea, Natalia Jimenez, Begoña Mellado, Sara Bystrup, Carme Solé-Blanch, Josep M. Piulats, José F. Suarez, Juan Carlos Pardo, Eva Martinez-Balibrea, Alvaro Aytes, Albert Font

## Abstract

**Background:** Taxanes are the most active chemotherapy agents in metastatic castration resistant prostate cancer (mCRPC) patients, yet resistance almost invariably occurs representing an important clinical challenge. Taxane-platinum combinations have shown clinical benefit in a subset of patients but the mechanistic basis and biomarkers remain elusive.

**Objective:** To identify mechanisms and response biomarkers for the antitumor efficacy of taxane-platinum combinations in mCRPC.

**Design, setting, and participants:** Transcriptomic data from a publicly available mCRPC dataset of taxane-exposed and naïve patients was analysed to identify response biomarkers and emerging vulnerabilities. Functional and preclinical validation was performed in taxane resistant mCRPC cell lines and genetically engineered mouse models (GEMM).

**Intervention:** mCRPC cells were treated with docetaxel, cisplatin, carboplatin and the CXCR2 inhibitor, SB265610. Gain and loss of function in culture of CXCR2 was achieved by overexpression or siRNA-silencing. Preclinical assays in GEMM mice tested the anti-tumor efficacy of taxane-platinum combinations.

**Outcome measurements and statistical analysis:** Proliferation, apoptosis and colony assays measured drug activity *in vitro*. Preclinical endpoints in mice included growth, survival and histopathology. Changes in CXCR2, BCL-2 and chemokines were analysed by RT-qPCR and Western Blot. Human expression data was analyzed using GSEA, hierarchical clustering and correlation studies. GraphPad Prism software, R-studio, were used for statistical and data analyses.

**Results and limitations:** Transcriptomic data from taxane-exposed human mCRPC tumors correlates with a marked negative enrichment of apoptosis and inflammatory response pathways accompanied by a marked downregulation of CXCR2 and BCL-2. Mechanistically, we show that docetaxel treatment inhibits CXCR2 and that BCL-2 downregulation occurs as a downstream effect. Further, we demonstrated that taxane resistance is directly associated to CXCR2 expression and that targeting of CXCR2 sensitizes prostate cancer (PC) cells to cisplatin. Finally, taxane-platinum combinations *in vivo* are highly synergistic and previous exposure to taxanes sensitizes mCRPC tumors to second line cisplatin treatment.

**Conclusions:** Together our data identifies an acquired vulnerability in taxane treated mCRPC patients with potential predictive activity for platinum-based treatments.

**Patient summary:** A subset of patients with aggressive and therapy resistant PC benefits from taxane-platinum combination chemotherapy however, we lack biomarkers and mechanistic basis about how that synergistic effect occurs. Here, using patient data and preclinical models, we found that taxanes reduce cancer cell scape mechanisms to chemotherapy-induced cell death, hence turning these cells more vulnerable to additional platinum treatment.

## Introduction

Prostate cancer (PC) remains the second leading cause of cancer-related mortality among men [1, 2]. Despite significant initial responses to androgen deprivation therapy, most metastatic patients progress to an incurable castration-resistant prostate cancer (mCRPC) stage that is frequently treated with docetaxel or antiandrogenic therapies [2–4]. Sadly, virtually most of them progress to aggressive variants, which often progress indifferently from androgen receptor (AR) signaling [5]. Yet, very few approved non-androgen targeting therapies are available with the exception of platinum-based treatments [6].

Taxanes bind tubulin inhibiting mitosis and AR nuclear transport [7]. To date, several factors have been associated with taxane resistance, including expression of β-tubulin isoforms and activation of drug efflux pumps. Importantly, PTEN loss, and activation of survival signaling pathways such as PI3K/AKT/mTOR, MAPK and NF-kB have also been associated to taxane resistance [8, 9]. Due to their different mechanism of action, platinums and taxanes are often combined in cancer therapy. While platinum agents are not routine for the treatment of mCRPC, there is an increasing use of these agents especially in those patients with a small-cell or neuroendocrine tumor variants [6]. In fact, some antitumor activity has been described for carboplatin, cisplatin, and satraplatin in mCRPC patients [10]. Unfortunately, molecular biomarkers to identificate aggressive variants of PC patients that could benefit from these drug combination remains elusive. This in part reflects an incomplete understanding of the underlying molecular mechanisms of taxane resistance, which limits the identification of vulnerabilities and potential therapeutic targets.

Here, we sought to elucidate the mechanistic basis and potential biomarkers for the synergistic antitumor efficacy of taxane-platinum combinations in mCRPC (Figure 1A). Mining of human PC datasets to compare the transcriptomes of taxane exposed and naïve patients showed a strong enrichment in the apoptosis pathway accompanied by a marked downregulation of CXCR2 and BCL-2 in taxane-exposed patients. Mechanistically, we showed that docetaxel treatment inhibits CXCR2 and that BCL-2 downregulation occurs as a downstream effect. Further, we demonstrated that taxane resistance is directly associated to CXCR2 expression and that targeting of CXCR2 sensitizes PC cells to cisplatin. Finally, we exploited genetically engineered mouse models (GEMM) of PC to test taxane-platinum combinations and sequential treatment in preclinical assays. Our *in vivo* data validates the hypothesis that taxane-platinum combinations are highly synergistic and that previous exposure to taxanes sensitizes mCRPC tumors to second line cisplatin treatment. Together our data identifies an acquired vulnerability in taxane-treated mCRPC patients with potential predictive value for platinum-based drug combinations.

**Figure 1.**
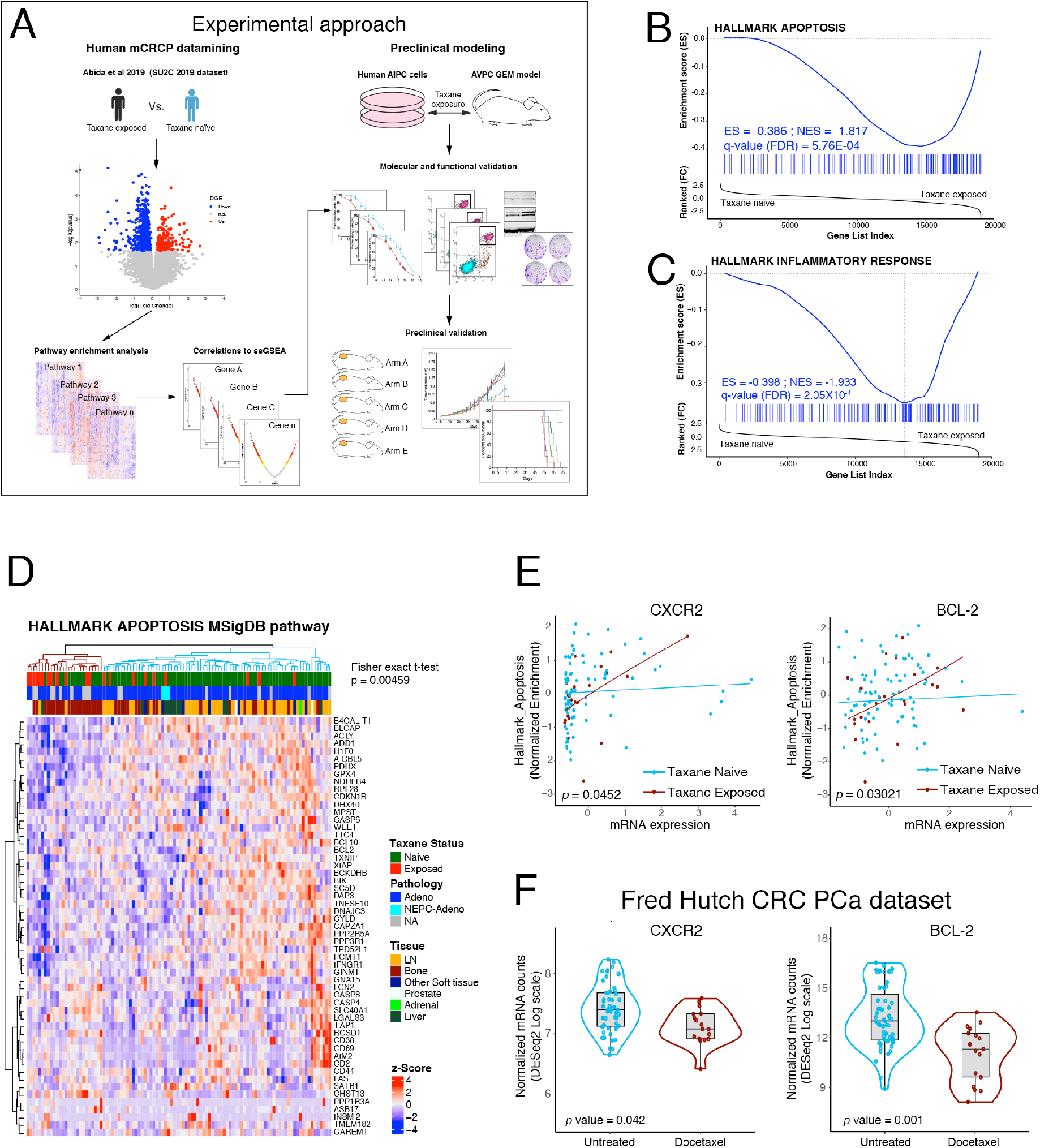
CXCR2 and BCL-2 are downregulated in human prostate cancer exposed to taxane. **(A)** Experimental approach of the study. Differential gene expression between docetaxel naïve and exposed PC patients in the SU2C dataset was used for pathway analysis and ssGSEA correlations. At the same time *in vivo* and *in vitro* modelling of taxane exposure was used for molecular and functional validation of mechanisms from human PC. Finally, preclinical studies were carried out to validate the antitumor efficacy of taxane-platinum combinations in aggressive AIPC. **(B-C)** GSEA showing a significant negative enrichment for the Apoptosis **(B)** and Inflammatory response **(C)** pathways from the Hallmarks Pathways of the MSigDB in the differential gene expression signature from the taxane naïve vs. taxane exposed tumors in the SU2C dataset. **(D)** Unsupervised clustering of patients according to the gene expression of the leading-edge genes from the GSEA in (B). Shown is the Fisher exact t-test p value for the association of taxane exposed status and downregulation of the signature as shown in the heatmap. **(E)** Linear regressions for the correlation between *CXCR2* (left) and *BCL-2* (right) expression and the Normalized Enrichment Score of the Apoptosis pathway. Taxane exposed and taxane naïve patients are shown separately. The p-value indicates the significance of the differential association between gene expression and pathway enrichment. **(F)** mRNA expression differences between taxane exposed and taxane naïve patients in the FHCRC dataset for *CXCR2* (left) and *BCL-2* (right).

## Materials and Methods

### Computational analysis of human prostate cancer data

Human transcriptome data was downloaded directly from Cbioportal [11, 17], used for differential gene expression between taxane exposed (N=22) and taxane naïve (N=82) using a two-sample two-tail Student’s t-test (Supplementary table 1) and Pathway enrichment using Gene Set Enrichment Analysis (GSEA) (Supplementary table 2). Univariated analyses were performed using Spearman rank correlation tests. For bivariate analysis linear regressions were performed (Supplementary table 3).

### Functional assays *in vitro*

Docetaxel resistant DU145-DR and PC3-DR human PC cells had been previously generated [12]. Docetaxel, cabazitaxel, cisplatin (MedChemExpress) and CXCR2 antagonist SB265610 (Sigma Aldrich) were prepared in DMSO or acetic acid:water (1:1). Western Blot, were performed as previously described [13] with primary and secondary antibodies shown in Supplementary Table 4. Cell viability was measured using an MTT assay (Roche Diagnostics), and the synergistic effect of cisplatin and SB265610 assessed on the Compusyn Software (Combosyn Inc.). Apoptosis was determined by using FITC Annexin V Apoptosis Detection Kit I (BD Pharmingen) following the manufacturer’s instructions with appropriate positive and negative controls. *CXCR2* silencing and overexpression in DU145 and/or PC3 cells was carried out as in [13]. RT-qPCR was performed as described previously [13]. Primer pairs used are listed in Supplementary Table 4.

### Preclinical assays *in vivo*

All studies involving mouse models were approved by the Institutional Review Board at IDIBELL. The NPK mouse had been previously published [14]. For preclinical tumor growth and survival assays, allografted NPK tumor bearing mice where enrolled in vehicle (saline), single or combination drug treatment with docetaxel, n=10 (oral gavage,2 mg/kg, once per week), cabazitaxel, n=10 (oral gavage, 2mg/kg, once per week) or cisplatin, n=10 (oral gavage, 2 mg/kg, once per week), monitored using calipers twice/week for the indicated time.

### Immunohistochemical analysis

Immunostaining of mouse prostate tumor tissues was done as described previously [15] on formalin-fixed paraffin-embedded sections incubated with primary and secondary antibodies shown in Supplementary Table 4

### Statistical analysis

Statistical differences between IC50 were determined by non-linear regression analysis and F-test. For viability, colony formation, flow cytometry and proliferation assays, p-values were calculated using a two-tailed Student’s t-test and values ≤ 0.05 were considered significant. Two-way analysis of variance (ANOVA) was used to calculate the significance of the difference between the vehicle and each treatment group. In survival analysis, p-values were calculated using a log-rank test. Where indicated, * means p < 0.01; ** p < 0.001 and *** p < 0.0001.

## Results

### The CXCR2/BCL-2 axis is attenuated in human PC exposed to taxane

Here, we first sought to investigate the molecular changes in PC tumors exposed to taxane therapy in a well-annotated human PC dataset [11]. GSEA identified a number of significantly enriched Hallmark Cancer Pathways (Supplementary figure 1A, Supplementary table 2) including a significantly negative enrichment in the “Androgen response” pathway (NES = −2.002; FDR q-value = 8.16×10^-5^). Interestingly, among others, the “Apoptosis” (NES = −1.817; FDR q-value = 5.76×10^-4^) and “Inflammatory response” (NES = −1.933964; FDR q-value = 2.05×10^-4^) pathways were also significantly negatively enriched in taxane-exposed mCRPC tumors (Figure 1B, C). In particular, unsupervised clustering of the GSEA leading-edge genes from the “Apoptosis” pathway significantly segregated patients exposed to taxanes in the SU2C dataset (p = 0.00459) (Figure 1D). Notably, significantly downregulated genes in the leading edge of these two pathways included the anti-apoptotic master regulator *BCL-2* (p = 0.0071) as well as several members of the CXCL superfamily, including *CXCL8* (p = 0.043) and *CXCL6* (p = 0.0075) (Supplementary table 1), known to bind the CXCR2 receptor and trigger a plethora of intracellular signal responses including anti-apoptosis programs [16]. In fact, *CXCR2* and *BCL-2* expression levels are significantly correlated with the negative enrichment of the “Apoptosis” pathway in taxane-exposed patients but not in taxane naïve ones (p value = 0.0452 and 0.03021, respectively) (Figure 1E; Supplementary table 3) and associated to decreased AR score and increased NEPC score (Supplementary figure 1B). Moreover, *CXCR2* and *BCL-2* mRNA levels were found significantly downregulated (p value = 0.042 and 0.001, respectively) in an independent human dataset [17] (Figure 1F). Together, these data indicate that taxane treatment attenuates the anti-apoptotic signaling mediated by CXCR2 and BCL-2 in mCRPC patients.

### Docetaxel induces CXCR2/BCL-2 downregulation *in vitro* and *in vivo*

Having observed a correlation between taxane exposure and CXCR2/BCL-2 downregulation in human PC, we next sought to investigate if this downregulation was causally linked to the treatment. First, we observed that docetaxel exposure in DU145 and PC3 cells triggers an immediate downregulation of CXCR2 and BCL-2 (Figure 2A). We then tested this causal relationship in the NPK GEMM PC model that is intrinsically castration resistant [14]. In agreement with the *in vitro* observation, tumors from the docetaxel-treated NPK mice showed a marked reduction in CXCR2 and BCL-2 compared to non-treated controls (Figure 2B).

**Figure 2.**
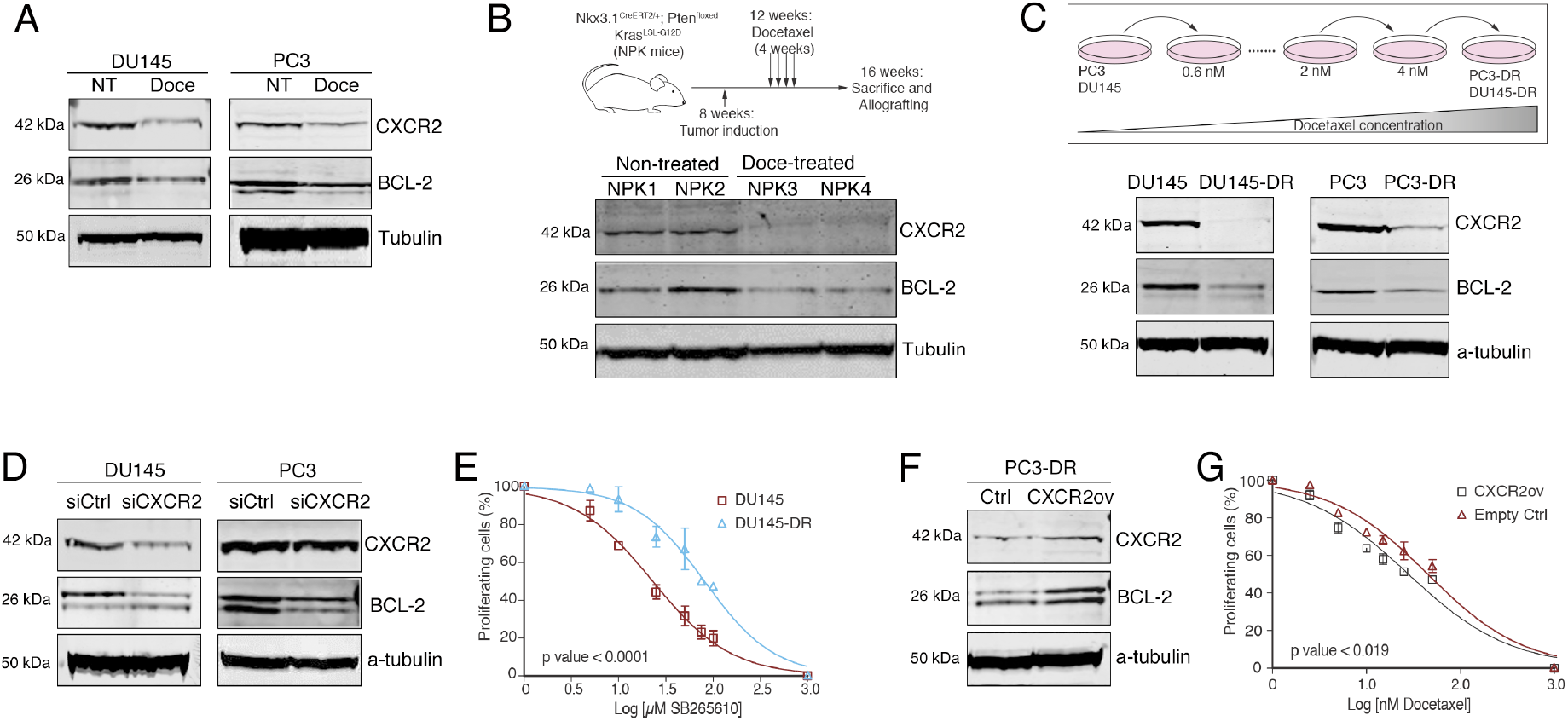
Effect of docetaxel treatment on CXCR2/BCL2 axis regulation. **(A)** Representative western blot images showing protein expression changes of CXCR2 and BCL-2 in DU145 and PC-3 cells after treatment with docetaxel (Doce) at 6.5 and 15 nM, respectively, for 72 hours. α-tubulin was used as endogenous control. NT: Non-treated cells. **(B)** Experimental strategy: one month after tamoxifen-mediated induction of the cre-dependent recombination, tumor-bearing mice (n=4) were exposed to 4 cycles of docetaxel treatment (oral gavage, 2 mg/kg, Monday to Friday) or vehicle (top). Western blot showing CXCR2 and BCL-2 protein expression changes in tumour tissues from NPK GEMM exposed to docetaxel or vehicle (Non-treated). α-tubulin was used as endogenous control (bottom) **(C)** Experimental strategy: DU145 and PC3 cells were converted to docetaxel-resistant cells by exposing them to increasing doses of docetaxel in an intermittent regimen during 1 year and 6 months, respectively (top). Western blot analysis of CXCR2 and BCL-2 basal protein expression in PC3/PC3-DR and DU145/DU145-DR cell lines. α-tubulin was used as endogenous control (bottom). *p-value < 0.05 and **p-value < 0.01; relative to protein expression in the corresponding parental cell line **(D)** Representative western blot images showing changes in CXCR2 and BCL-2 protein expression in PC3 and DU145 cells under negative control (siCtrl) and *CXCR2* (siCXCR2) gene silencing. α-tubulin was used as endogenous control. **(E)** Dose response curves for DU145 and DU145-DR cells after SB265610 treatment at 0-100 *μ*M for 72h (mean ± SEM). **(F)** Western blot analysis of CXCR2 and BCL-2 protein expression changes under empty control (Crtl) and *CXCR2* overexpression (CXCR2ov) in PC3-DR cells. α-tubulin was used as endogenous control. **(G)** Dose-response curves for PC3-DR cell line, after *CXCR2* overexpression (CXCR2ov), treated with 0-50 nM docetaxel for 72 hours (mean ± SEM). Results shown were obtained from at least 3 independent experiments. P-values were calculated using a two-tailed Student’s t-test.

To test whether downregulation of CXCR2 is maintained in docetaxel resistance we evaluated CXCR2 and BCL-2 levels in two docetaxel resistant cell lines, hereafter named DU145-DR and PC3-DR (Supplementary Figure 2A). Consistently with a role in resistance, protein levels of CXCR2 and BCL-2 in untreated DU145-DR and PC3-DR cells were profoundly reduced (Figure 2C). In addition, CXCR2 ligands *CXCL8* and *CXCL6* were also significantly downregulated in the DU145-DR and PC3-DR cells (Supplementary figure 2B), as was *CXCR1*, which has been shown to be co-regulated with CXCR2 [18] (Supplementary figure 2C). Notably, reduced *CXCR2* expression was due to transcriptional downregulation (Supplementary figure 2C) and not promoter hypermethylation as treatment with the 5-AZA did not increase protein expression (Supplementary figure 2D).

Depletion of CXCR2 or CXCL8/CXCL6 has been shown by others and us to promote BCL-2 downregulation in different tumor types [13, 19, 20]. Confirming that the observed downregulation of BCL-2 in the human PC cohort and in our *in vitro* and *in vivo* models was indeed a downstream effect of CXCR2 depletion, silencing of CXCR2 in parental DU145 and PC3 cells (Supplementary figure 2E) induced a marked reduction in BCL-2 protein expression compared with control cells (Figure 2D). Additionally, since DU145-DR cells are depleted of CXCR2 we predicted that these cells would be resistant to the CXCR2 inhibitor SB265610. Data showed that DU145-DR cells were 4-fold more resistant to CXCR2 antagonist treatment as compared to its parental cells (Figure 2E). Finally, to demonstrate that docetaxel resistance is at least in part mediated by CXCR2, we induced expression of CXCR2 in PC3-DR cells. As expected, CXCR2 overexpression promoted a BCL-2 increase (Figure 2F) and significantly sensitized PC3-DR cells to docetaxel (Figure 2G). Collectively, these results suggest that docetaxel treatment induces a downregulation of CXCR2 and its downstream effector BCL-2, *in vitro* and *in vivo,* and that this downregulation is, at least in part, associated to the acquisition of docetaxel resistance.

### CXCR2 downregulation sensitizes PC cells to platinum

Others and we have shown that inhibition of CXCR2 signaling and its downstream effector BCL-2 increases the sensitivity of tumor cells to platinum drugs [13, 20, 21]. Based on our results indicating that docetaxel triggers a marked reduction in the CXCR2/BCL-2 axis, and the clinical benefit observed for the combination of platinums and taxanes in mCRPC [22], we hypothesized that the docetaxel-resistant PC cells sensitivity to platinum treatment is CXCR2-dependent (Figure 3). As predicted, docetaxel-resistant DU145-DR and PC3-DR cells were significantly more sensitive to cisplatin and carboplatin that their respective parental cells (DU145-DR, p = 0.004 and <0.0001; PC3-DR p < 0.0001 and < 0.0001 for cisplatin and carboplatin, respectively) (Figure 3A, B).

**Figure 3.**
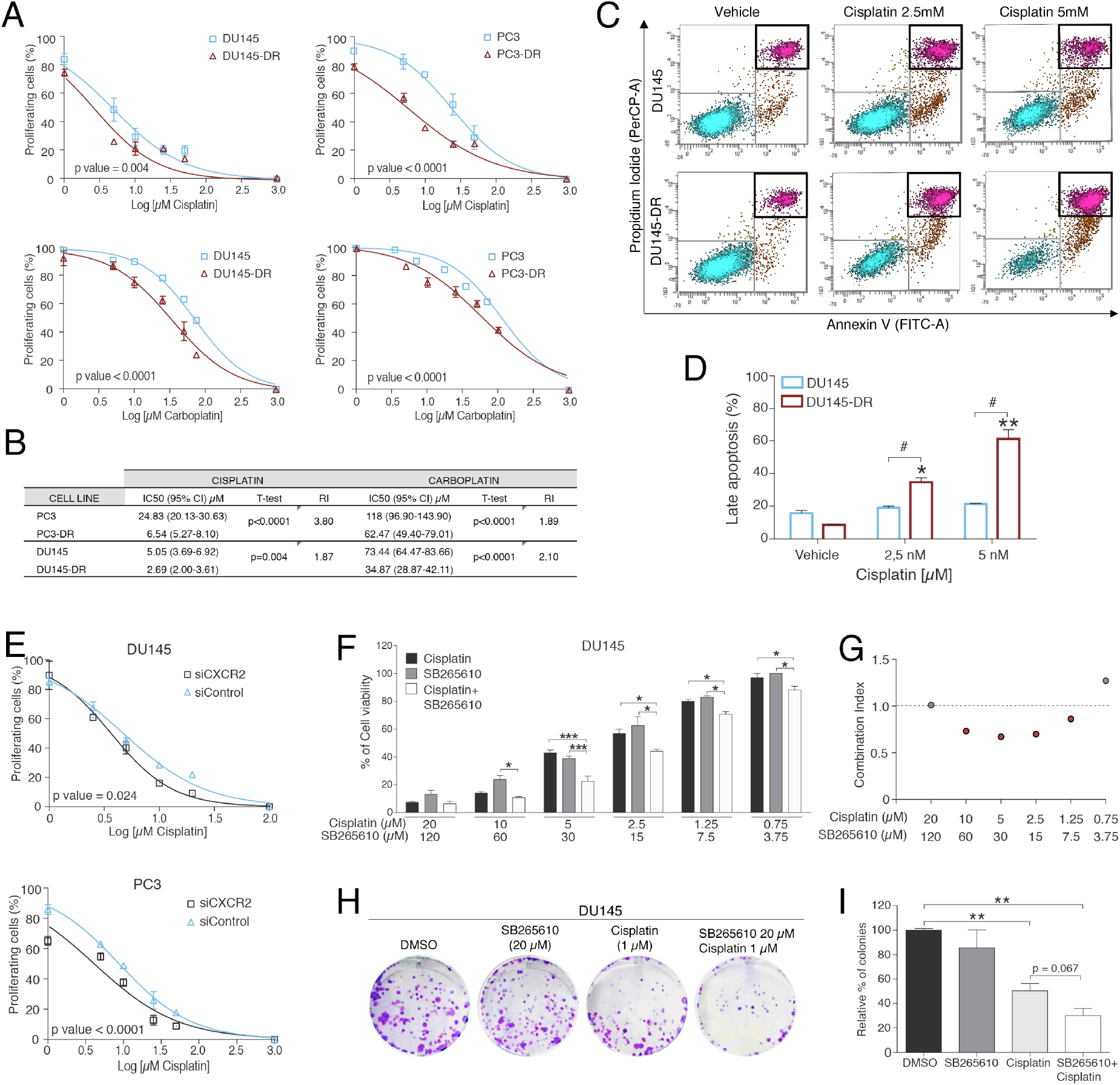
Effect of CXCR2 downregulation on platinum sensitivity in PC cells. **(A)** Dose response curves for PC3/PC3-DR and DU145/DU145-DR cells after cisplatin treatment at 0-50 *μ*M or carboplatin treatment at 0-75 *μ*M for 72h (mean ± SEM). **(B)** Table showing cisplatin and carboplatin IC50 values indicated as mean (95% CI) in PC3/PC3-DR and DU145/DU145-DR cell lines. R.I: Resistance Index, calculated as the ratio between IC50 of resistant sublines and its corresponding sensitive cell lines. **(C)** Representative images of apoptosis activation after 72h-cisplatin treatment in DU145 and DU145-DR cells. **(D)** Bar graph representing the percentage (mean ± SEM) of late apoptotic cells after 72h of treatment with cisplatin in DU145 and DU145-DR cell lines at their corresponding IC50 doses, 2,5 and 5 *μ*M, respectively. *p-value < 0.05; **p-value < 0.01, relative to vehicle. *#*p-value < 0,05, relative to late apoptosis rate after cisplatin treatment in DU145 cells. **(E)** Dose-response curves for PC3 and DU145 cell lines, after *CXCR2* gene silencing (siCXCR2), treated with 0-50 *μ*M cisplatin for 72h (mean ± SEM). siControl: cells transfected with a silencer negative transcription control **(F)** Bar graphs representing mean ± SEM percentage of cell viability after a 72-hour treatment with cisplatin, SB265610 or their concomitant combination at the indicated doses in DU145 cells. *p-value < 0.05; ***p-value < 0.001 relative to the indicated treatment condition. **(G)** Dot plot representing Combination Index values calculated for each combined treatment dose. **(H)** Representative colony assay images after treatment with cisplatin, SB265610 or its combination at the indicated doses in DU145 cells **(I)** Bar graph representing the percentage (mean ± SEM) of colonies in DU145 cells after 24h of the indicated treatments. **p-value < 0.01 relative to vehicle (DMSO). All results were obtained from at least 3 independent experiments. P-values were calculated using a two-tailed Student’s t-test.

Next, to determine the impact on apoptosis after platinum treatment in docetaxel-resistant cells, DU145 and DU145-DR cells were treated with cisplatin at their respective IC50 doses and apoptosis rates were measured by flow cytometry with the Annexin V/PI assay. In agreement with the observed downregulation of anti-apoptotic signaling in the taxane treated patients of the SU2C human PC cohort (Figure 1), cisplatin induced a significant dose dependent increase in late apoptosis rates only in the docetaxel resistant DU145-DR cells but not in the parental cells (p < 0.01 and < 0.001 for 2.5 and 5 µM of cisplatin, respectively) (Figure 3C,D).

To ascertain the role of CXCR2 on cisplatin sensitivity, we next silenced CXCR2 with siRNA and assessed the cytotoxicity of cisplatin treatment in control and siCXCR2 cells. In agreement with the shown downregulation of BCL-2 upon CXCR2 silencing (Figure 2D) and its impact on apoptosis, siCXCR2 DU145 and PC3 cells displayed a significantly enhanced sensitivity for cisplatin compared to non-targeting siRNA control cells (p = 0.024 and < 0.0001, respectively) (Figure 3E). Importantly, treatment of docetaxel sensitive parental DU145 cells with cisplatin plus the CXCR2 inhibitor SB265610 showed significant synergistic effects on cell viability compared to either one single treatment on a wide range of concentrations (Figure F,G). Finally, this synergistic interaction between cisplatin and SB265610 was confirmed in clonogenic assays where the combined treatment markedly reduced colony formation compared to single agents (Figure 3H, I)

Taken together, these data indicate that docetaxel-resistant PC cells are more sensitive to cisplatin treatment, and suggest that this vulnerability is in part mediated, by the downregulation of the CXCR2/BCL-2 signalling.

### Taxane treatment sensitizes a mouse CRPC model to cisplatin treatment

To our knowledge, at least two independent randomized phase 1 and phase 1-2 clinical trials [10, 22] have assessed the efficacy of taxane-planinum combinations in men with mCRPC with apparently contradictory results. To provide preclinical evidence and to test the hypothesis that platinum added to taxane may improve survival and response in aggressive mCRPC, we conducted similar trials in NPK PC GEMM (Figure 4A). As in the parental mice, allografted NPK tumors develop aggressive metastatic PC with 100% penetrance that are AR positive but insensitive to castration or anti-AR treatment [14]. As expected, no significant reduction in tumor growth was observed in mice treated with either docetaxel, cabazitaxel or cisplatin alone albeit a significant improve in survival on mice treated with docetaxel (Median survival of 65 ± 4.9 vs. 55.5 ± 4 days, p < 0.001) and cabazitaxel (61 ± 3.14 days vs. 55.5 ± 4 days, p = 0.0031) but not cisplatin (56 ± 1.7 days vs. 55.5 ± 4 days, p = 0.33) in “Preclinical assay 1” (Figure 4B). Importantly, a significant reduction in tumor growth was observed when either docetaxel or cabazitaxel were combined with cisplatin (p < 0.01 and 0.001, respectively) and a remarkable improvement in survival for both combinations [(20% reaching endpoint in docetaxel+cisplatin; median survival of 75 ± 2.2 days vs. 55.5 ± 4 days, p = 1.2×10^-10^); (0% reaching endpoint in cabazitaxel+cisplatin; median survival 75 ± 0 days vs. 55.5 ± 4 days p = 6.8×10^-12^)] (Figure 4B-C), We next asked whether previous exposure to taxane sensitizes otherwise resistant tumors to cisplatin treatment. To test this in “Preclinical assay 2”, we engrafted mice with control, docetaxel-treated and cabazitaxel-treated NPK tumors from “preclinical assay 1” and treated them with cisplatin for the indicated time (Figure 4D). Notably, sequential cisplatin post docetaxel or cabazitaxel significantly reduced tumor growth (p < 0.01, both) and improved survival [(33% reaching endpoint in cisplatin post docetaxel; median survival of 40.5 ± 5.5 days vs. 32.5 ± 2.3 days, p = 0.00491); (50% reaching endpoint in cisplatin post cabazitaxel; median survival 40.5 ± 4.8 days vs. 32.5 ± 2.3 days p = 0.0019)] (Figure 4E) Finally, immmunostaining of tumors from “Preclinical assay 1” showed that proliferation was significantly reduced in tumors treated with taxane plus cisplatin compared to either of the single taxane treatments (p < 0.0001), thereby explaining the improved anti-tumor response observed (Figure 4F-H). Interestingly, nuclear AR expression was also further reduced in combination treatments (p < 0.0001 and 0.001 as shown). Together, this *in vivo* preclinical data validates the hypothesis that taxane-cisplatin combinations have anti-tumor efficacy in aggressive mCRPC and that taxane resistant tumors are sensitized to cisplatin treatment.

**Figure 4.**
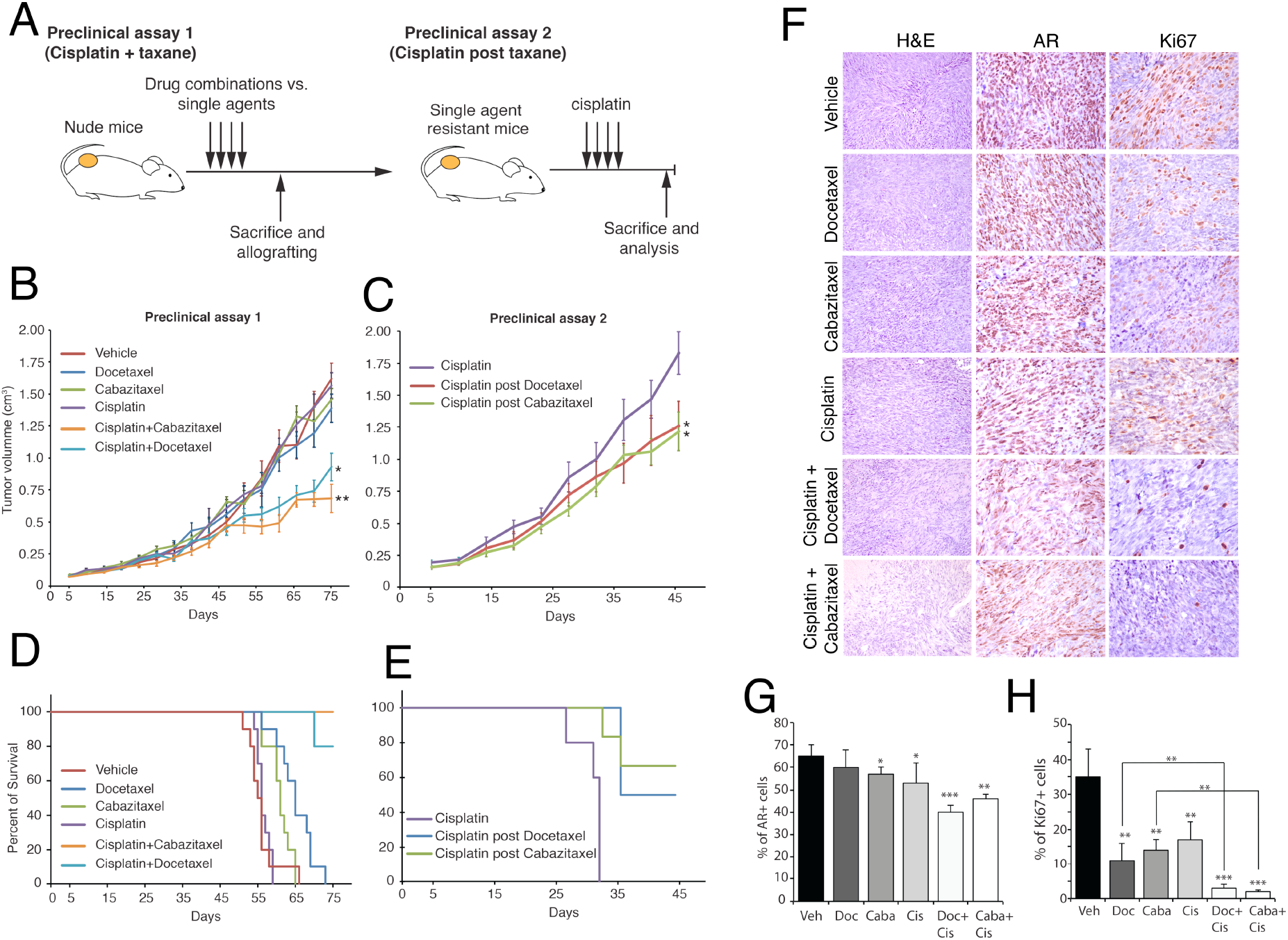
Preclinical validation. **(A)** Scheme of the preclinical assays carried out. Preclinical Assay 1 assessed the anti-tumor efficacy of single agent docetaxel, cabazitaxel or cisplatin versus docetaxel plus cisplatin or cabazitaxel plus cisplatin. Single agent treated tumors from Preclinical assay 1 where shifted to cisplatin to test whether previous taxane exposure would sensitize for cisplatin treatment. **(B)** Tumor growth curves for NPK allografts treated with the indicated drugs or vehicle for 75 days. **(C)** Tumor growth curves for single docetaxel, cabazitaxel or vehicle treated tumors from (B) under cisplatin treatment for 45 days. ANOVA is used to assessed the significance of the differences in tumor growth. *, p < 0.01; ** p < 0.001. **(D-E)** Survival curves for mice enrolled on Preclinical assay 1 **(D)** or Preclinical assay 2 **(E). (F)** Immunohistochemistry for AR and Ki-67 in tumors from Preclinical assay 1. **(G-H)** Quantification of nuclear AR staining **(G)** and Ki-67 staining **(H)**. Shown is the percent of positive nuclear stain over total number of cells. *, p < 0.01; ** p < 0.001; p < 0.0001

## Discussion

Docetaxel, before or after anti-androgens, remains a standard of care for mCRPC patients. Yet, none of these treatments are curative and a subset will eventually progress to aggressive variants of PC (AVPC), which are androgen indifferent (AIPC). Taxane-platinum combinations have shown clinical benefit for a group of patients in clinical trials for this disease stage [6], highlighting the need for predictive response biomarkers. Exploiting publicly available datasets we have found that apoptosis and inflammatory response pathways are dysregulated and that CXCR2/ BCL-2 axis is markedly reduced in taxane exposed PC tumors. Assays *in vitro* and *in vivo* suggest that while loss of CXCR2 and BCl-2 cooperate to induce taxane resistance, they represent an emerging vulnerability against genotoxic platinum-based treatments. Finally, in agreement with other studies [23], our preclinical trials *in vivo* confirm that taxane exposure sensitizes PC cells to platinum.

In addition to stabilizing microtubules, taxanes induce apoptosis by inhibiting antiapoptotic proteins, such as BCL-2 [24]. Accordingly, following treatment with paclitaxel, BCL-2 is phosphorylated in PC3 and LnCaP cells, inhibiting its anti-apoptotic action and enhancing cell death [25, 26]. Consistent with our results, BCL-2 downregulation after docetaxel treatment has been shown in breast cancer cells [27]. Downregulation of a druǵs target and signaling rewiring to elicit pro-survival pathways is a well-known mechanism of cancer treatment resistance [28]. Therefore, our data suggest that mCRPC patients lose BCL-2 expression, at least in part, through CXCR2 downregulation as a mechanism to evade docetaxel-induced apoptosis. In accordance with previous data [13, 20, 21], we demonstrated that the inhibition of the CXCR2 chemokine receptor markedly increased the sensitivity of AIPC cells to cisplatin treatment, suggesting that the inverse relationship between docetaxel- and cisplatin-resistance could be mediated, at least in part, by CXCR2 downregulation after taxane exposure. Conversely, other studies also demonstrated that when cells become resistant to platinum they often become sensitive to taxanes [29].

A phase I/II trial recently demonstrated that men with AVPC have a better Progression free survival after cabazitaxel + carboplatin treatment as compared to cabazitaxel alone [22]. In agreement, our *in vivo* preclinical results validate these data suggesting that therapeutic response to cisplatin-based chemotherapy in docetaxel-resistant mCRPC patients might be predicted by the level of CXCR2/BCL-2 expression before treatment with this platinum. In other words, CXR2/BCL-2– negative mCRPC cases might have a better therapeutic response to cisplatin-based chemotherapy than CXCR2/BCL-2–positive cases.

## Conclusions

CXCR2/BCL-2 anti-apoptotic axis is markedly reduced in taxane exposed PC tumors representing an emerging vulnerability towards genotoxic platinum-based treatments. Therefore, taxane resistant tumors are sensitized to cisplatin treatment and taxane-cisplatin combinations have anti-tumor efficacy in aggressive mCRPC. Our findings also suggest that therapeutic response to cisplatin-based treatment in docetaxel-resistant mCRPC patients might be predicted by the levels of CXCR2/BCL-2 expression.

## Author contributions

Albert Font and Alvaro Aytes had full access to all the data in the study and take responsibility for the integrity of the data and the accuracy of the data analysis.

### Conception and design

Ruiz de Porras, Martinez-Balibrea, Mellado, Font, Aytes

### Acquisition of data

Ruiz de Porras, Wang, Palomero, Marin-Aguilera, Indacochea, Bystrup, Solé-Blanch, Jimenez

### Analysis and interpretation of data

Ruiz de Porras, Palomero, Indacochea, Mellado, Aytes, Font.

### Drafting of the manuscript

Ruiz de Porras, Aytes, Font

### Critical revision of the manuscript for important intellectual content

All authors

### Statistical analysis

Ruiz de Porras, Palomero, Indacochea, Aytes, *Obtaining funding:* Font, Aytes.

### Administrative, technical, or material support

Ruiz de Porras, Wang, Palomero, Marin-Aguilera, Bystrup, Solé-Blanch, Jimenez

### Supervision

Font, Aytes

### Other

None.

## Financial disclosures

Albert Font and Alvaro Aytes certify that no conflicts of interest, including specific financial interests and relationships and affiliations relevant to the subject matter or materials discussed in the manuscript (eg, employment/affiliation, grants or funding, consultancies, honoraria, stock ownership or options, expert testimony, royalties, or patents filed, received, or pending) exist.

## Funding/Support and role of the sponsor

This work was supported by funding from has the “Badalona Foundation against cancer” grant (A.F.); Instituto de Salut Carlos III (PI16/01070 and CP15/00090) to A.A.; the European Association of Urology Research Foundation (EAURF/407003/XH) to A.A,; Fundacion BBVA to A.A.; Department of Defense Award (W81XWH-18-1-0193) to A.A.; the CERCA Program / Generalitat de Catalunya, and FEDER funds/ European Regional Development Fund (ERDF)-a way to Build Europe.

## Acknowledgements statement

V.R.d.P and A.F acknowledge the “Badalona Foundation against cancer”. A.I. acknowledges the Spanish Ministry (MEIC) to EMBL partnership, Severo Ochoa and the CERCA program.

