## Supplementary materials for "Taxane induces attenuation of the CXCR2/BCL-2 axis and sensitize prostate cancer to platinum-based treatments"

*Ruiz de Porras et al.*

### Extended materials and methods

### Supplementary Figures:

Supplementary Figure 1. Pathway analysis on Docetaxel exposed human mCRPC patients. (Related to Figure 1)

Supplementary Figure 2. CXCR2, CXCR1 and CXCL-chemokine expression in docetaxel-sensitive and resistant cell lines (Related to Figure 2)

### Supplementary Tables

Supplementary Table 1. Differentially expressed genes in taxane-exposed vs. naïve PC patients in the SU2C dataset

Supplementary Table 2. Pathway enrichment analysis

Supplementary Table 3. CXCR2 and BCL-2 correlation to Hallmark pathway enrichment

Supplementary Table 4. List of antibodies and primers used in this study (see Extended materials and methods)

**Extended materials and methods:**

**Computational analysis of human prostate cancer data:** Human mCRPC transcriptome data was downloaded directly from Cbioportal [11]. Differential gene expression between taxane exposed (N=22) and taxane naïve (N=82) patients was estimated using a two-sample two-tail Student's t-test (Supplementary table 1) and Pathway enrichment using Gene Set Enrichment Analysis (GSEA) on the Hallmark Pathways dataset from the MSigDB (Supplementary table 2). An unsupervised cluster analysis was done based on the leading edge genes of the Hallmarks apoptosis pathway. Samples annotation data was retrieved directly from SU2C clinical data. Univariated analyses were performed using Spearman rank correlation tests. For bivariate analysis comparing the interaction effects between gene expression and taxane status over different scores, linear regressions were performed (Supplementary table 3). For each analysis, predicted data was normalized using bestNormalize method at homonymous R package. Gene expression microarray from the Fred Hucht CRC PC dataset was compared between untreated controls (n =57) and samples treated only with docetaxel (n = 18). T-test was performed to compare the mean differences between both groups.

**Functional assays *in vitro*:** Docetaxel resistant DU145-DR and PC3-DR human PC cells had been previously generated [12]. Docetaxel, cabazitaxel, cisplatin (MedChemExpress) and CXCR2 antagonist SB265610 (Sigma Aldrich) was prepared in DMSO and stored at 4 °C. A 5-Aza-2'-deoxycytidine (5-AZA, Sigma Aldrich) stock was prepared in acetic acid:water (1:1) at 50mg/ml.

Western Blot, were performed as previously described [13] with primary and secondary antibodies shown in Supplementary Table 4 and scanned in an Odyssey Imaging System (LICOR Biosciences).

Cell viability was measured using an MTT assay (Roche Diagnostics), seeding 6.000 to 8.000 cells/well in 96-well plates and treating with different drug concentrations for 72 h. The synergistic effect of cisplatin and SB265610 was assessed by calculating the Combination Index (CI; CI < 1 indicates synergy) on the Compusyn Software (Combosyn Inc.).

Apoptosis was determined by using FITC Annexin V Apoptosis Detection Kit I (BD Pharmingen) following the manufacturer's instructions in a FACS Canto II flow cytometer (Becton Dickinson Immunocytometry System) with appropriate positive and negative controls.

Colony-formation assays were performed as previously described [13], seeding 500 cells/well treated for 24 h, cultured in complete media for 10 days, washed, fixed with a Methanol/acetic acid (3:1) solution and stained with crystal violet (0,5%) for 10 min.

*CXCR2* were silenced in DU145 and PC3 cells with siRNAs pools as described in [13] (Smartpool On-target plus: *CXCR2* siRNA, #L-005647; Dhamacon, GE) and non-targeting controls (#AM4611; Ambion). CXCR2 was overexpressed by transfection the ORF cloned into the pcDNA3.1$^{+}$/C-(K)DYK vector (GenScript) (*Clone ID: OHu23649*) using Lipofectamine 2000 (Thermo Fisher Scientific).

RT-qPCR was performed as described previously [13]. Primer pairs used are listed in Supplementary Table 4. Relative gene expression quantification was calculated according to the comparative Ct method using β-Actin as endogenous control.

**Preclinical assays *in vivo*:** All studies involving mouse models were approved by the Institutional Review Board at IDIBELL. The Nkx3.1$^{CreERT2/+}$; Pten$^{floxed/floxed}$; Kras$^{lsl\text{-}G12D/+}$ (NPK) mouse have been previously published [14]. For preclinical tumor growth and survival assays, allografted NPK tumor bearing mice where enrolled in vehicle (saline), single or combination drug treatment with docetaxel, n=10 (oral gavage,2 mg/kg, once per week), cabazitaxel, n=10 (oral gavage, 2mg/kg, once per week) or cisplatin, n=10 (oral gavage, 2 mg/kg, once per week). Briefly, treatments were initiated 5 days after engraftment and tumor growth monitored using calipers twice/week until tumor size reached an endpoint of 1.75 $cm^3$ in the control group, at which point tumors where excised and processed for histological analysis. For survival analysis, a tumor size of 1,25 $cm^3$ was predefined as the endpoint.

**Immunohistochemical analysis:** Immunostaining of mouse prostate tumor tissues was done as described previously [15]. Immunohistochemistry was performed on formalin-fixed paraffin-embedded sections. Slides were de-paraffinized and heat-mediated antigen retrieval was performed using the citrate-based antigen unmasking solution (Vector, H-3300) and incubated with primary and secondary antibodies shown in Supplementary Table 4

**Statistical analysis:** In all functional assays *in vitro,* data are presented as mean ± SEM of at least 2 independent experiments of 3 replicates and the statistical analysis was performed with Graphpad Prism V.4 software. Statistical differences between IC50 were determined by graphic representation of dose-response curves and subsequent non-linear regression analysis and F-test. For viability, colony formation, flow cytometry and proliferation assays, p-values were calculated using a two-tailed Student's t-test and values ≤ 0.05 were considered significant. Two-way analysis of variance (ANOVA) was used to calculate the significance of the difference between the vehicle and each treatment group. In survival analysis, p-values were calculated using a log-rank test. Where indicated, * means $p<0.01$; ** $p<0.001$ and *** $p<0.0001$.

<table>
<tr><th colspan="6">Supplementary Table 4: List and description of antibodies and primers used in this study</th></tr>
<tr><td colspan="6">Primary antibodies</td></tr>
<tr><td rowspan="2">Antigen</td><td rowspan="2">Company</td><td rowspan="2">Catalog #</td><td rowspan="2">Type</td><td colspan="2">Use and dilution</td></tr>
<tr><td>IHC</td><td>Western</td></tr>
<tr><td>CXCR2</td><td>Abcam</td><td>14935</td><td>Rabbit mAb</td><td>1:100</td><td>1:1000</td></tr>
<tr><td>BCL-2</td><td>Abcam</td><td>32124</td><td>Rabbit mAb</td><td>1:100</td><td>1:1000</td></tr>
<tr><td>Tubulin</td><td>Sigma</td><td>T6074</td><td>Mouse mAb</td><td></td><td>1:10000</td></tr>
<tr><td>AR</td><td>Abcam</td><td>133273</td><td>Rabbit mAb</td><td>1:200</td><td></td></tr>
<tr><td>Ki-67</td><td>NovusBio</td><td>NB600-1252</td><td>Rabbit pAb</td><td>1:50</td><td></td></tr>
<tr><td colspan="6">Secondary antibodies</td></tr>
<tr><td>IRDye anti-rabbit</td><td>LICOR Biosciences</td><td>926-68071</td><td></td><td></td><td>1:10000</td></tr>
<tr><td>IRDye anti-mouse</td><td>LICOR Biosciences</td><td>926-32210</td><td></td><td></td><td>1:10000</td></tr>
<tr><td>Biotinylated anti-rabbit IgG</td><td>Vector</td><td>BA-1000</td><td></td><td>1:300</td><td></td></tr>
<tr><td colspan="6">Oligonucleotides</td></tr>
<tr><td>Target</td><td colspan="2">Forward oligo</td><td colspan="3">Reverse Oligo</td></tr>
<tr><td>CXCR2</td><td colspan="2">TATGAGGACATGGGCAACAA</td><td colspan="3">AGGGTGAATCCGTAGCAGAA</td></tr>
<tr><td>CXCR1</td><td colspan="2">TGCATCAGTGTGGACCGTTA</td><td colspan="3">TGTCATTTCCCAGGACCTCA</td></tr>
<tr><td>CXCL8</td><td colspan="2">TCTTGGCAGCCTTCCTGATTTC</td><td colspan="3">GTGTGGTCCACTCTCTCAATCACTCT</td></tr>
<tr><td>CXCL6</td><td colspan="2">ACTTGTTTACGCGTTACGCTGAG</td><td colspan="3">TTCTTCAGGGAGGCTACCACTT</td></tr>
<tr><td>Actin</td><td colspan="2">TGAGCGCGGCTACAGCTT</td><td colspan="3">TCCTTAATGTCACGCACGATTT</td></tr>
</table>

**Supplementary Figure 1**

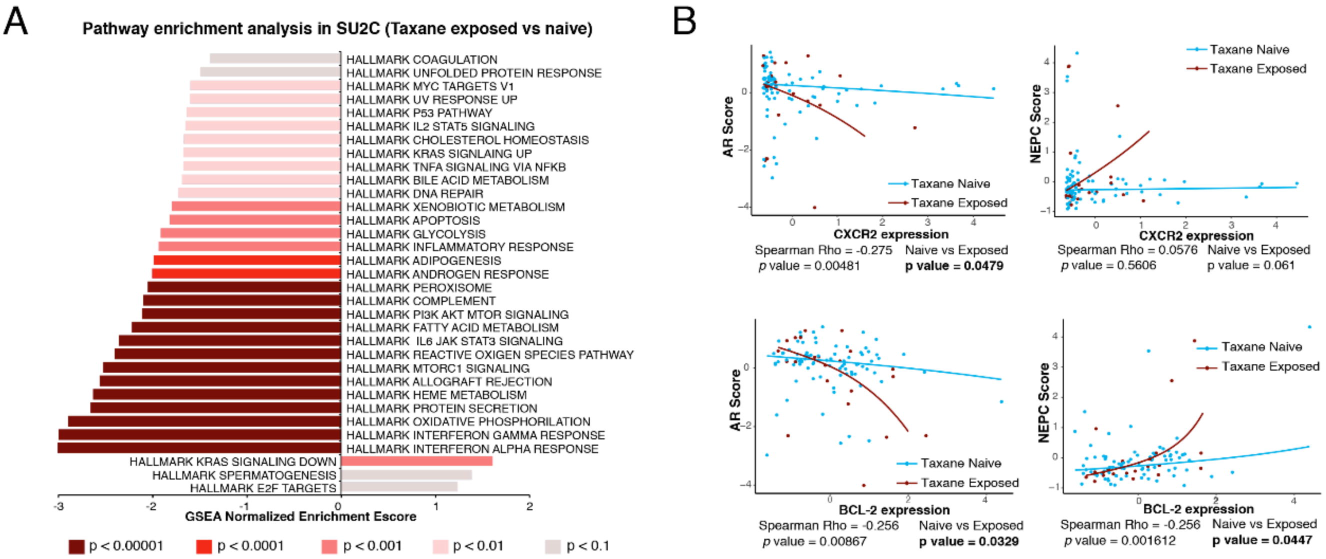


**Legend: Pathway analysis on taxane exposed human mCRPC patients. (A)** Shown if the negative and positive normalized enrichment score for the GSEA on the Hallmarks pathways of the MSigDB against the differential expression signature between taxane exposed and taxane naïve patients in the SU2C dataset. **(B)** Regression showing the correlation between gene expression levels for CXCR2 (top) and BCL-2 (Bottom) and the AR Score (left) and the NEPC score (right). Spearman Rho and p value are shown for all patients. The “naïve vs Exposed” p value indicates the significance of the differential association between gene expression and AR or NEPC score.

**Supplementary Figure 2**

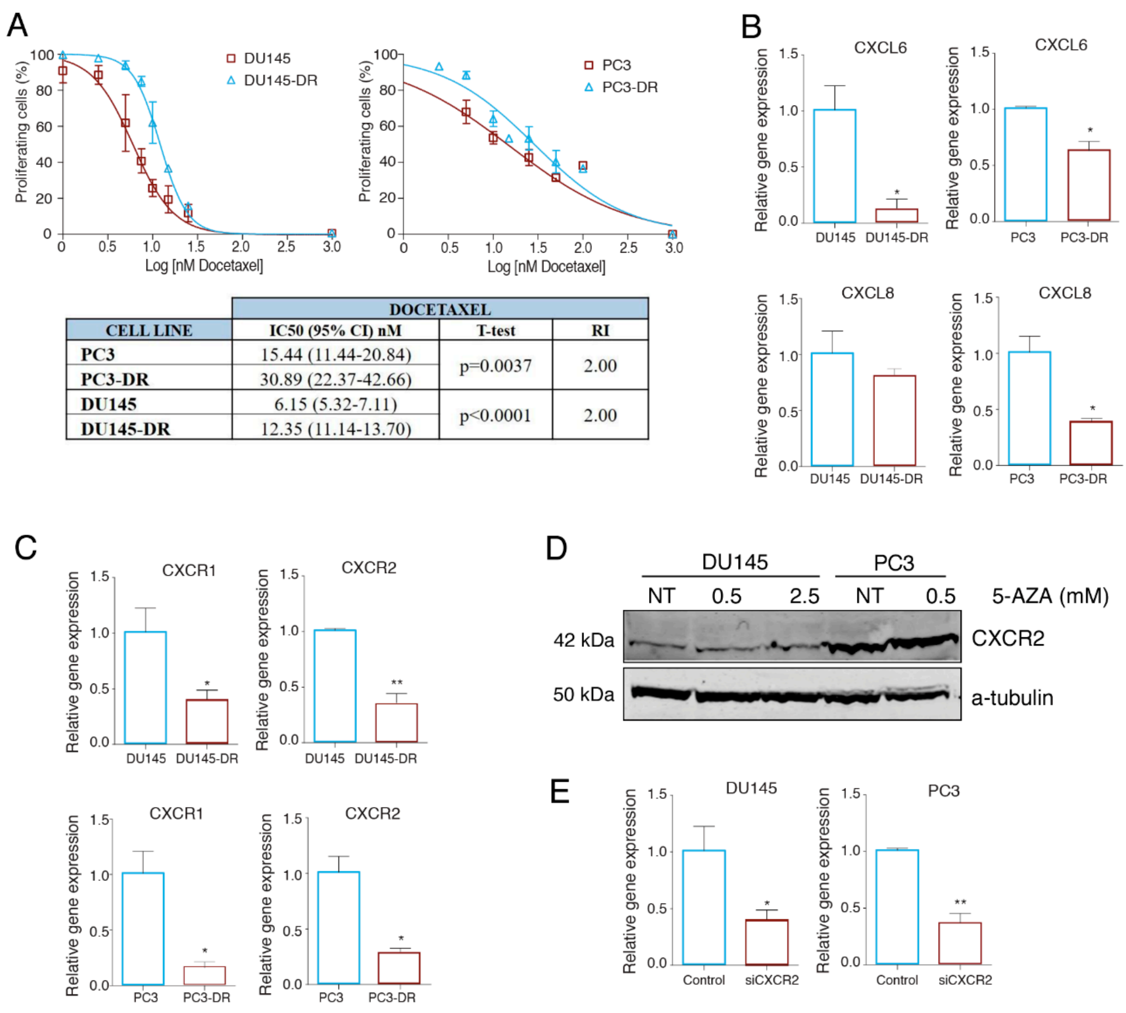


| CELL LINE | DOCETAXEL | | |
|---|---|---|---|
| | IC50 (95% CI) nM | T-test | RI |
| PC3 | 15.44 (11.44-20.84) | p=0.0037 | 2.00 |
| PC3-DR | 30.89 (22.37-42.66) | | |
| DU145 | 6.15 (5.32-7.11) | p<0.0001 | 2.00 |
| DU145-DR | 12.35 (11.14-13.70) | | |

**Legend: CXCR2, CXCR1 and CXCL-chemokine expression in docetaxel-sensitive and resistant cell lines (A)** Effect of docetaxel treatment on docetaxel resistant cells proliferation. Dose response curves for PC3/PC3-DR and DU145/DU145-DR cells after docetaxel treatment at 0-25 nM for 72h (mean ± SEM) (top). Table showing docetaxel IC50 values, indicated as mean (95% CI), for each cell line. R.I: Resistance Index, calculated as the ratio between IC50 of resistant sublines and its corresponding sensitive cell lines (bottom) **(B)** Bar graph illustrating relative gene expression levels (mean ± SEM) of *CXCL8* and *CXCL6* in PC3/PC3-DR and DU145/DU145-DR cells. Gene expression levels of β-actin were used as endogenous control. *p-value < 0.05; relative to gene expression in the corresponding docetaxel sensitive cell line. **(C)** Graphic representation of *CXCR1 and CXCR2* relative gene expression (mean ± SEM), respectively, in PC3/PC3-DR and DU145/DU145-DR cell lines. Gene expression levels of β-actin were used as endogenous control. *p-value < 0.05 and **p-value < 0.01; relative to gene expression in the corresponding parental cell line. (**D)** Representative western blot images showing protein expression changes of CXCR2 in DU145-DR and PC3-DR cells after treatment with 0,5 and 2,5 μM of 5-AZA for 72h. α-tubulin was used as endogenous control. **(E)** Bar graphs illustrating relative gene expression levels (mean ± SEM) of *CXCR2* after its siRNA-mediated gene silencing (siCXCR2) in PC3 and DU145 cells. *p-value < 0.05; relative to control cells (siNTC). All results were obtained from at least 3 independent experiments. P-values were calculated using a two-tailed Student's t-test.

| Supplementary table 1. Differentially expressed genes in taxene-exposed vs. naïve PC SU2C patients | | | | | |
|---|---|---|---|---|---|
| geneName | meanNaive | meanExposed | foldChangeLog2 | tStatistic | pvalue |
| SLCO1B3 | 0.9542 | 0.1982 | 34.5 | -4017.000 | 0.000113 |
| DTX2 | 2695.000 | 2.31 | 0.8446 | -4107.000 | 0.0001167 |
| AGA | 3423.000 | 2839.000 | 0.8479 | -4296.000 | 0.0001314 |
| TMEM126B | 4533.000 | 4066.000 | 0.9281 | -4189.000 | 0.0001464 |
| THAP2 | 1151.000 | 0.883 | -0.8873 | -4117.000 | 0.000222 |
| OST4 | 7584.000 | 6909.000 | 0.954 | -4051.000 | 0.0002833 |
| PSMA2 | 6184.000 | 5839.000 | 0.9684 | -3766.000 | 0.0003104 |
| DNAJB9 | 4.19 | 3631.000 | 0.9001 | -3869.000 | 0.0003351 |
| SLC36A1 | 3126.000 | 3.95 | 1205.000 | 3892.000 | 0.0003399 |
| GLRX2 | 3738.000 | 3182.000 | 0.8778 | -3696.000 | 0.0004844 |
| SUMF2 | 5631.000 | 5192.000 | 0.9531 | -3815.000 | 0.0004943 |
| UQCRQ | 7206.000 | 6737.000 | 0.966 | -3669.000 | 0.0005383 |
| CCDC126 | 2207.000 | 1909.000 | 0.8169 | -3661.000 | 0.0005751 |
| RFESD | 0.8845 | 0.603 | 4122.000 | -3581.000 | 0.001 |
| ADM5 | 0.8029 | 1193.000 | -0.8045 | 3627.000 | 0.0007541 |
| SRD5A3 | 3022.000 | 2483.000 | 0.8224 | -3653.000 | 0.000766 |
| DNAJC3 | 4405.000 | 3982.000 | 0.932 | -3546.000 | 0.000819 |
| SCN2A | 0.5421 | 0.2463 | 2288.000 | -3515.000 | 0.0009056 |
| HTATIP2 | 3574.000 | 2915.000 | 0.8401 | -3589.000 | 0.0009085 |
| GRB2 | 5154.000 | 4775.000 | 0.9534 | -3618.000 | 0.0009111 |
| BCL10 | 2956.000 | 2575.000 | 0.8729 | -3616.000 | 0.0009184 |
| TROVE2 | 3314.000 | 2936.000 | 0.899 | -3629.000 | 0.0009246 |
| BTC | 1272.000 | 0.785 | -1005.000 | -3536.000 | 0.0009308 |
| MRPL32 | 5802.000 | 5304.000 | 0.949 | -3596.000 | 0.0009685 |
| KIAA0922 | 3.07 | 2511.000 | 0.8208 | -3596.000 | 0.0009748 |
| PYROXD1 | 1965.000 | 1616.000 | 0.7103 | -3536.000 | 0.0009808 |
| HELB | 0.7025 | 0.5536 | 1675.000 | -3488.000 | 0.001017 |
| CTSO | 4157.000 | 3585.000 | 0.8961 | -3537.000 | 0.001192 |
| UEVLD | 2763.000 | 2464.000 | 0.8872 | -3469.000 | 0.001194 |
| MRPL54 | 5428.000 | 4821.000 | 0.9299 | -3533.000 | 0.001224 |
| TTC1 | 5259.000 | 4894.000 | 0.9566 | -3503.000 | 0.001228 |
| ATRAID | 6232.000 | 5838.000 | 0.9643 | -3.46 | 0.001284 |
| MPLKIP | 2316.000 | 1893.000 | 0.7598 | -3507.000 | 0.001284 |
| GSAP | 2658.000 | 2249.000 | 0.8292 | -3448.000 | 0.001325 |
| SPATS2L | 3679.000 | 3008.000 | 0.8454 | -3487.000 | 0.00135 |
| CMAS | 4727.000 | 4172.000 | 0.9196 | -3486.000 | 0.001384 |
| CDC73 | 3837.000 | 3468.000 | 0.9249 | -3469.000 | 0.001406 |
| PKD1 | 2861.000 | 3343.000 | 1148.000 | 3456.000 | 0.001434 |
| ZBED6 | 0.1094 | 0.06623 | 1227.000 | -3288.000 | 0.001448 |
| ERLEC1 | 5371.000 | 4817.000 | 0.9352 | -3.42 | 0.001503 |
| TNFAIP3 | 2399.000 | 1968.000 | 0.7741 | -3314.000 | 0.0017 |
| RTCA | 3858.000 | 3504.000 | 0.9288 | -3396.000 | 0.001745 |
| EDEM2 | 4233.000 | 3.85 | 0.9343 | -3363.000 | 0.001748 |
| MAP3K6 | 2635.000 | 3048.000 | 1.15 | 3.38 | 0.001749 |

| | | | | | |
|---|---|---|---|---|---|
| KLHL7 | 2715.000 | 2359.000 | 0.8595 | -3359.000 | 0.001752 |
| AKAP8 | 3423.000 | 3804.000 | 1086.000 | 3421.000 | 0.001778 |
| DDAH1 | 4.39 | 3834.000 | 0.9085 | -3369.000 | 0.001819 |
| CYLD | 2574.000 | 2198.000 | 0.833 | -3351.000 | 0.001831 |
| ETFRF1 | 3574.000 | 2884.000 | 0.8314 | -3369.000 | 0.001909 |
| LYN | 2942.000 | 2.35 | 0.7919 | -3336.000 | 0.001942 |
| SC5D | 4988.000 | 4432.000 | 0.9265 | -3292.000 | 0.001957 |
| SLCO1A2 | 0.3765 | 0.1158 | 2207.000 | -3179.000 | 0.001986 |
| EIF2AK1 | 5886.000 | 5497.000 | 0.9614 | -3342.000 | 0.002061 |
| C7orf26 | 3917.000 | 3589.000 | 0.936 | -3.35 | 0.00207 |
| TCEANC | 1523.000 | 1258.000 | 0.5447 | -3291.000 | 0.002112 |
| NXT2 | 2398.000 | 2003.000 | 0.7942 | -3.17 | 0.002123 |
| COX4I1 | 7019.000 | 6624.000 | 0.9703 | -3231.000 | 0.002236 |
| TMEM106B | 2822.000 | 2453.000 | 0.865 | -3233.000 | 0.002254 |
| AGR3 | 2447.000 | 1345.000 | 0.3308 | -3194.000 | 0.002289 |
| MALSU1 | 2969.000 | 2666.000 | 0.9011 | -3.23 | 0.00234 |
| ZDHHC11B | 0.6738 | 1093.000 | -0.225 | 3326.000 | 0.002371 |
| SMDT1 | 4941.000 | 4.44 | 0.9331 | -3267.000 | 0.002375 |
| LTA4H | 5194.000 | 4849.000 | 0.9583 | -3248.000 | 0.002381 |
| TSG101 | 4743.000 | 4432.000 | 0.9564 | -3212.000 | 0.002467 |
| FRK | 1493.000 | 1075.000 | 0.1794 | -3176.000 | 0.002478 |
| TMEM59 | 5001.000 | 4599.000 | 0.948 | -3221.000 | 0.002491 |
| SLFNL1 | 0.4444 | 0.7583 | 0.3412 | 3365.000 | 0.002493 |
| PRKACB | 3545.000 | 3016.000 | 0.8722 | -3223.000 | 0.002561 |
| CCNG1 | 5605.000 | 4988.000 | 0.9323 | -3286.000 | 0.002588 |
| EIF3M | 4668.000 | 4313.000 | 0.9486 | -3205.000 | 0.002597 |
| TPPP | 1407.000 | 1844.000 | 1793.000 | 3232.000 | 0.002637 |
| MLYCD | 1056.000 | 0.8918 | -2093.000 | -3109.000 | 0.002815 |
| FANCF | 2957.000 | 2515.000 | 0.8507 | -3.22 | 0.002862 |
| COMMD1 | 4271.000 | 3784.000 | 0.9166 | -3202.000 | 0.002934 |
| PRDX4 | 6407.000 | 5843.000 | 0.9504 | -3177.000 | 0.002943 |
| MAPKAPK2 | 4636.000 | 4174.000 | 0.9317 | -3166.000 | 0.00305 |
| COPS9 | 5162.000 | 4474.000 | 0.9129 | -3206.000 | 0.003052 |
| ELP4 | 0.8262 | 0.7021 | 1853.000 | -3151.000 | 0.003053 |
| CAB39 | 4401.000 | 4012.000 | 0.9376 | -3144.000 | 0.003062 |
| SEC61G | 6792.000 | 6172.000 | 0.9501 | -3176.000 | 0.003105 |
| MT1E | 4273.000 | 5457.000 | 1168.000 | 3094.000 | 0.003115 |
| C7orf25 | 3033.000 | 2777.000 | 0.9205 | -3077.000 | 0.00315 |
| FKBP11 | 4641.000 | 4065.000 | 0.9137 | -3161.000 | 0.003249 |
| ZNF512B | 3199.000 | 3.61 | 1104.000 | 3175.000 | 0.003304 |
| C11orf74 | 3341.000 | 3007.000 | 0.9126 | -3114.000 | 0.003306 |
| CCDC90B | 3239.000 | 2913.000 | 0.9097 | -3156.000 | 0.003332 |
| YKT6 | 5151.000 | 4785.000 | 0.955 | -3151.000 | 0.00338 |
| ZNF277 | 2783.000 | 2337.000 | 0.8296 | -3169.000 | 0.003446 |
| RGS13 | 0.744 | 0.2389 | 4842.000 | -2998.000 | 0.003488 |
| WIPI2 | 4362.000 | 4042.000 | 0.9482 | -3132.000 | 0.003508 |

| | | | | | |
|---|---|---|---|---|---|
| NECAP2 | 4.74 | 4.43 | 0.9564 | -3169.000 | 0.003529 |
| AIMP2 | 4206.000 | 3748.000 | 0.9198 | -3136.000 | 0.003531 |
| TTLL3 | 2542.000 | 3.11 | 1216.000 | 3112.000 | 0.003539 |
| TRAF6 | 1997.000 | 1793.000 | 0.8443 | -3069.000 | 0.003706 |
| GPN1 | 3772.000 | 3442.000 | 0.9312 | -3067.000 | 0.00378 |
| CTBS | 2.1 | 1708.000 | 0.7218 | -3069.000 | 0.003825 |
| FUT9 | 0.1209 | 0.01476 | 1995.000 | -2.97 | 0.003842 |
| RAP1A | 3842.000 | 3535.000 | 0.9381 | -3078.000 | 0.003855 |
| ZNF593 | 3348.000 | 2884.000 | 0.8766 | -3093.000 | 0.003865 |
| TANK | 3716.000 | 3361.000 | 0.9236 | -3084.000 | 0.003883 |
| VPS26B | 4.91 | 4363.000 | 0.9257 | -3053.000 | 0.003895 |
| ZNF77 | 1786.000 | 2057.000 | 1243.000 | 3125.000 | 0.003946 |
| FUZ | 3258.000 | 3648.000 | 1096.000 | 3042.000 | 0.003987 |
| IDH1 | 6644.000 | 5815.000 | 0.9296 | -3125.000 | 0.003989 |
| GPR37 | 1449.000 | 0.8812 | -0.3413 | -3057.000 | 0.004013 |
| ARHGEF25 | 2586.000 | 3049.000 | 1174.000 | 3044.000 | 0.004042 |
| MTMR6 | 2871.000 | 2505.000 | 0.8706 | -3079.000 | 0.004052 |
| TMEM190 | 0.6545 | 0.917 | 0.2043 | 3107.000 | 0.004058 |
| MRPS9 | 4389.000 | 4039.000 | 0.9437 | -3078.000 | 0.004076 |
| TMEM263 | 4255.000 | 3795.000 | 0.921 | -3086.000 | 0.004201 |
| MB | 3731.000 | 2.96 | 0.8242 | -2991.000 | 0.004304 |
| ARSD | 4344.000 | 3656.000 | 0.8826 | -3101.000 | 0.004337 |
| IPCEF1 | 0.5878 | 0.3925 | 1.76 | -2951.000 | 0.004378 |
| DESI2 | 3466.000 | 3.04 | 0.8945 | -3034.000 | 0.004418 |
| TTC9C | 4275.000 | 3966.000 | 0.9483 | -3055.000 | 0.004456 |
| STK17A | 2933.000 | 2462.000 | 0.8373 | -3024.000 | 0.004472 |
| ARPC2 | 6824.000 | 6.42 | 0.9682 | -3074.000 | 0.004484 |
| BIRC3 | 1828.000 | 1346.000 | 0.492 | -2962.000 | 0.00449 |
| ADCY3 | 2595.000 | 2983.000 | 1146.000 | 3016.000 | 0.004495 |
| U2AF1 | 0.5241 | 0.8548 | 0.2428 | 3047.000 | 0.004547 |
| DTX3L | 4263.000 | 3712.000 | 0.9046 | -3051.000 | 0.004572 |
| PSME1 | 7102.000 | 6634.000 | 0.9652 | -3022.000 | 0.004673 |
| NUDT8 | 4062.000 | 2946.000 | 0.7707 | -3.03 | 0.004704 |
| TMBIM4 | 4534.000 | 4.06 | 0.9269 | -3023.000 | 0.004815 |
| SAP30L | 3121.000 | 2805.000 | 0.9063 | -3026.000 | 0.004818 |
| FAM46D | 0.05509 | 0.0185 | 1376.000 | -2878.000 | 0.004884 |
| SMYD2 | 4524.000 | 4102.000 | 0.9352 | -2924.000 | 0.004887 |
| TNFSF10 | 5671.000 | 4701.000 | 0.8919 | -3027.000 | 0.004892 |
| POMP | 5681.000 | 5208.000 | 0.9501 | -3016.000 | 0.004921 |
| LMTK2 | 2904.000 | 2598.000 | 0.8955 | -2974.000 | 0.004993 |
| TRAPPC2L | 3181.000 | 2842.000 | 0.9026 | -2949.000 | 0.004996 |
| TMED9 | 6947.000 | 6.52 | 0.9673 | -2997.000 | 0.005128 |
| TSTD3 | 2.11 | 1.6 | 0.6299 | -3027.000 | 0.005147 |
| TMEM156 | 0.665 | 0.415 | 2156.000 | -2869.000 | 0.005165 |
| GNPTAB | 4106.000 | 3607.000 | 0.9084 | -2941.000 | 0.005191 |
| HBP1 | 3983.000 | 3644.000 | 0.9355 | -2949.000 | 0.005245 |

| | | | | | |
|---|---|---|---|---|---|
| RARRES3 | 4116.000 | 3246.000 | 0.8321 | -2955.000 | 0.00529 |
| C3orf58 | 2693.000 | 2136.000 | 0.7661 | -3.01 | 0.005356 |
| CPOX | 3383.000 | 3101.000 | 0.9286 | -2874.000 | 0.005364 |
| CAPZA1 | 5785.000 | 5458.000 | 0.9668 | -2962.000 | 0.0054 |
| SSR3 | 5816.000 | 5411.000 | 0.959 | -2957.000 | 0.005449 |
| DOCK3 | 0.5455 | 1289.000 | -0.4191 | 3036.000 | 0.00548 |
| MYO19 | 3441.000 | 3926.000 | 1107.000 | 2.98 | 0.005577 |
| TNFSF18 | 0.3259 | 0.1725 | 1567.000 | -2845.000 | 0.005589 |
| CCDC160 | 1459.000 | 0.9084 | -0.2544 | -2917.000 | 0.005592 |
| LRG1 | 2374.000 | 1353.000 | 0.3494 | -2.92 | 0.005615 |
| DNALI1 | 1984.000 | 2976.000 | 1592.000 | 2952.000 | 0.005681 |
| SPAG8 | 0.833 | 1223.000 | -1103.000 | 2966.000 | 0.005683 |
| ADAM33 | 0.5623 | 1551.000 | -0.7618 | 3052.000 | 0.005725 |
| CAT | 5391.000 | 4832.000 | 0.935 | -2954.000 | 0.005775 |
| C20orf96 | 3057.000 | 3451.000 | 1108.000 | 2895.000 | 0.005792 |
| DAD1 | 7583.000 | 6958.000 | 0.9575 | -2988.000 | 0.005797 |
| TALDO1 | 6965.000 | 6466.000 | 0.9617 | -2981.000 | 0.005836 |
| TTC14 | 3392.000 | 3031.000 | 0.9079 | -2947.000 | 0.005913 |
| IFNGR2 | 5619.000 | 5299.000 | 0.9661 | -2916.000 | 0.005914 |
| SRI | 4.69 | 4.29 | 0.9424 | -2.9 | 0.005921 |
| ZNF343 | 2091.000 | 2.37 | 1.17 | 2947.000 | 0.006005 |
| VPS41 | 3646.000 | 3387.000 | 0.9431 | -2916.000 | 0.006109 |
| UFC1 | 6511.000 | 6.16 | 0.9704 | -2.9 | 0.00615 |
| TPPP3 | 2343.000 | 3312.000 | 1406.000 | 2977.000 | 0.006178 |
| AGL | 2534.000 | 2209.000 | 0.8524 | -2895.000 | 0.006256 |
| ACOT13 | 3161.000 | 2.83 | 0.9037 | -2862.000 | 0.006333 |
| CCNYL1 | 2554.000 | 2268.000 | 0.8734 | -2914.000 | 0.00644 |
| C2orf49 | 2615.000 | 2393.000 | 0.9076 | -2836.000 | 0.006564 |
| NOP10 | 6.79 | 6273.000 | 0.9587 | -2906.000 | 0.006612 |
| WFIKKN1 | 0.6245 | 1005.000 | -0.01082 | 2939.000 | 0.006634 |
| FAM104B | 2.9 | 2.56 | 0.8828 | -2838.000 | 0.006687 |
| MTHFD2L | 1968.000 | 1661.000 | 0.7495 | -2.86 | 0.006786 |
| COX7A2L | 5037.000 | 4543.000 | 0.9362 | -2903.000 | 0.006823 |
| C9orf16 | 5664.000 | 5071.000 | 0.9362 | -2938.000 | 0.006847 |
| KDELR2 | 7006.000 | 6564.000 | 0.9666 | -2881.000 | 0.006869 |
| ANKRD13A | 3705.000 | 3392.000 | 0.9326 | -2845.000 | 0.006872 |
| TMEM167A | 4454.000 | 4065.000 | 0.9389 | -2885.000 | 0.006891 |
| ZBTB41 | 2231.000 | 1942.000 | 0.827 | -2849.000 | 0.006907 |
| TBRG1 | 3295.000 | 2964.000 | 0.9113 | -2862.000 | 0.00691 |
| BCAS2 | 4988.000 | 4645.000 | 0.9557 | -2854.000 | 0.006941 |
| STYXL1 | 4157.000 | 3829.000 | 0.9424 | -2824.000 | 0.006943 |
| RHAG | 0.5939 | 0.2087 | 3007.000 | -2752.000 | 0.007021 |
| TMED3 | 3007.000 | 2505.000 | 0.834 | -2892.000 | 0.007036 |
| RNF5 | 5744.000 | 5208.000 | 0.9439 | -2909.000 | 0.007046 |
| APOL2 | 3933.000 | 3469.000 | 0.9083 | -2875.000 | 0.007082 |
| DAGLB | 3.14 | 2905.000 | 0.9318 | -2836.000 | 0.007107 |

| | | | | | |
|---|---|---|---|---|---|
| PSMC3IP | 2123.000 | 2522.000 | 1229.000 | 2871.000 | 0.007111 |
| YAF2 | 1346.000 | 1198.000 | 0.6071 | -2846.000 | 0.007113 |
| OSTF1 | 4856.000 | 4509.000 | 0.9531 | -2834.000 | 0.007127 |
| BCL2 | 3.23 | 2945.000 | 0.9211 | -2845.000 | 0.007172 |
| MRPL58 | 4995.000 | 4471.000 | 0.931 | -2868.000 | 0.007393 |
| ADPRHL1 | 0.8075 | 0.5594 | 2717.000 | -2789.000 | 0.0075 |
| CXCL6 | 0.6808 | 0.6002 | 1328.000 | -0.3127 | 0.007563 |
| SVIP | 3578.000 | 3071.000 | 0.8801 | -2839.000 | 0.007653 |
| NUPL2 | 3922.000 | 3631.000 | 0.9436 | -2825.000 | 0.007683 |
| ARFRP1 | 3532.000 | 3861.000 | 1071.000 | 2823.000 | 0.007684 |
| FDCSP | 1206.000 | 0.2434 | -7553.000 | -4715.000 | 0.008 |
| NIPAL2 | 3624.000 | 2909.000 | 0.8292 | -2875.000 | 0.007787 |
| USP25 | 3055.000 | 2688.000 | 0.8855 | -2.85 | 0.007804 |
| TSPAN17 | 4163.000 | 3891.000 | 0.9526 | -2803.000 | 0.007832 |
| ARHGDIB | 5979.000 | 5424.000 | 0.9456 | -2798.000 | 0.007929 |
| TNFRSF11A | 1019.000 | 0.7068 | -18.6 | -2805.000 | 0.007931 |
| CLEC16A | 2.51 | 2779.000 | 1.11 | 2806.000 | 0.007966 |
| LINC00998 | 4895.000 | 4247.000 | 0.9106 | -2822.000 | 0.007972 |
| COBLL1 | 3479.000 | 2884.000 | 0.8495 | -2821.000 | 0.008017 |
| PPP1CC | 5192.000 | 4.77 | 0.9485 | -2838.000 | 0.008023 |
| TIGD2 | 2003.000 | 1677.000 | 0.7441 | -2839.000 | 0.008076 |
| TMEM43 | 4644.000 | 4955.000 | 1042.000 | 2834.000 | 0.008098 |
| TAX1BP1 | 5526.000 | 5222.000 | 0.967 | -2776.000 | 0.008102 |
| RBM43 | 1768.000 | 1419.000 | 0.6137 | -2817.000 | 0.008104 |
| GTF2H1 | 3173.000 | 2925.000 | 0.9294 | -2812.000 | 0.008112 |
| ZNF621 | 2367.000 | 2674.000 | 1141.000 | 2845.000 | 0.008121 |
| MRPL33 | 6274.000 | 5.71 | 0.9488 | -2827.000 | 0.008121 |
| MRPL44 | 4417.000 | 4032.000 | 0.9386 | -2829.000 | 0.008126 |
| BBS12 | 1098.000 | 0.9032 | -1086.000 | -2744.000 | 0.00819 |
| KCND1 | 0.5829 | 0.8689 | 0.2604 | 2845.000 | 0.008223 |
| VASP | 5.08 | 4737.000 | 0.9569 | -2778.000 | 0.008363 |
| TOM1 | 4158.000 | 3.86 | 0.9479 | -2795.000 | 0.008363 |
| CLN5 | 2.37 | 2029.000 | 0.82 | -2794.000 | 0.008364 |
| UBE2J1 | 5008.000 | 4.53 | 0.9378 | -2811.000 | 0.008422 |
| TRAFD1 | 4.05 | 3737.000 | 0.9425 | -2823.000 | 0.008533 |
| TXNL1 | 3522.000 | 3.16 | 0.9138 | -2778.000 | 0.008576 |
| ELANE | 0.9799 | 0.3317 | 54.44 | -2679.000 | 0.008626 |
| TTC21A | 1293.000 | 1631.000 | 1905.000 | 2807.000 | 0.008628 |
| PHPT1 | 6426.000 | 5993.000 | 0.9625 | -2789.000 | 0.008695 |
| SPRN | 1608.000 | 1974.000 | 1431.000 | 2752.000 | 0.008773 |
| SMYD4 | 2056.000 | 2358.000 | 1191.000 | 2797.000 | 0.008898 |
| YIPF1 | 4.68 | 4257.000 | 0.9387 | -2752.000 | 0.009003 |
| LOXL3 | 1232.000 | 1541.000 | 2071.000 | 2777.000 | 0.009065 |
| KLRC2 | 0.4721 | 0.1645 | 2405.000 | -2658.000 | 0.009119 |
| GCA | 3465.000 | 3072.000 | 0.9029 | -2726.000 | 0.009167 |
| FBXO33 | 2.57 | 2334.000 | 0.8978 | -2746.000 | 0.009244 |

| | | | | | |
|---|---|---|---|---|---|
| FAM218A | 1097.000 | 0.7707 | -2.8 | -2745.000 | 0.009301 |
| ST3GAL6 | 1.6 | 1245.000 | 0.4664 | -2734.000 | 0.009335 |
| LINC01207 | 1283.000 | 0.6765 | -1569.000 | -2702.000 | 0.009356 |
| MMP8 | 0.4479 | 0.1087 | 2764.000 | -2649.000 | 0.009362 |
| KRCC1 | 4406.000 | 3776.000 | 0.896 | -2794.000 | 0.009362 |
| SLC26A10 | 1127.000 | 1756.000 | 4708.000 | 2797.000 | 0.009442 |
| ABCA13 | 0.2073 | 0.04021 | 2042.000 | -2651.000 | 0.009485 |
| TSPAN6 | 3325.000 | 2823.000 | 0.8638 | -2706.000 | 0.009579 |
| COQ10B | 3376.000 | 3035.000 | 0.9127 | -2763.000 | 0.009604 |
| CRY1 | 3236.000 | 2911.000 | 0.91 | -2726.000 | 0.009624 |
| PORCN | 2.62 | 2966.000 | 1129.000 | 2732.000 | 0.009665 |
| TSPYL2 | 4.19 | 4689.000 | 1079.000 | 2739.000 | 0.009711 |
| CDK5RAP1 | 3506.000 | 3737.000 | 1051.000 | 2748.000 | 0.009726 |
| LIN7C | 3.56 | 3235.000 | 0.9244 | -2722.000 | 0.009728 |
| CMPK1 | 5831.000 | 5437.000 | 0.9602 | -2735.000 | 0.009798 |
| WDR90 | 2916.000 | 3419.000 | 1149.000 | 2729.000 | 0.009913 |
| ACYP2 | 2096.000 | 1759.000 | 0.7633 | -2713.000 | 0.009922 |
| TMEM126A | 4639.000 | 4266.000 | 0.9454 | -2725.000 | 0.009925 |
| KCNAB3 | 0.7766 | 1146.000 | -0.5382 | 2754.000 | 0.01 |
| AZU1 | 0.8555 | 0.2822 | 8104.000 | -2621.000 | 0.01011 |
| UPRT | 3.12 | 2751.000 | 0.8893 | -2711.000 | 0.01012 |
| RPF1 | 4308.000 | 3893.000 | 0.9307 | -2733.000 | 0.01028 |
| CHMP2B | 4481.000 | 4092.000 | 0.9394 | -2717.000 | 0.01036 |
| NOL12 | 2964.000 | 2705.000 | 0.9157 | -2703.000 | 0.01049 |
| SLC35A3 | 2414.000 | 1988.000 | 0.7797 | -2704.000 | 0.01056 |
| DNAJC30 | 3378.000 | 2896.000 | 0.8737 | -2713.000 | 0.01072 |
| C11orf70 | 0.9539 | 0.6564 | 8914.000 | -2624.000 | 0.01075 |
| SCX | 1375.000 | 2077.000 | 2294.000 | 2735.000 | 0.01085 |
| ERCC5 | 3064.000 | 2758.000 | 0.906 | -2687.000 | 0.01094 |
| RAB1A | 5915.000 | 5642.000 | 0.9735 | -2686.000 | 0.01105 |
| NAT8L | 0.9146 | 1274.000 | -2712.000 | 2656.000 | 0.01114 |
| TMEM123 | 6313.000 | 5821.000 | 0.956 | -2698.000 | 0.01118 |
| VPS37B | 4157.000 | 3749.000 | 0.9276 | -2706.000 | 0.01124 |
| EIF3K | 6385.000 | 6125.000 | 0.9776 | -2655.000 | 0.01125 |
| KAT2B | 2843.000 | 2314.000 | 0.8028 | -2706.000 | 0.01127 |
| LIMD2 | 2776.000 | 2225.000 | 0.7835 | -2679.000 | 0.01131 |
| ATP5E | 5135.000 | 4797.000 | 0.9584 | -2648.000 | 0.01133 |
| RGS10 | 5708.000 | 5039.000 | 0.9284 | -2692.000 | 0.01134 |
| FKBP2 | 5426.000 | 4974.000 | 0.9486 | -2.69 | 0.01135 |
| TMCO3 | 4058.000 | 3723.000 | 0.9385 | -2651.000 | 0.01141 |
| TRAIP | 1946.000 | 2348.000 | 1282.000 | 2.7 | 0.01142 |
| ASNSD1 | 4085.000 | 3698.000 | 0.9293 | -2685.000 | 0.01157 |
| TMEM258 | 5122.000 | 4646.000 | 0.9402 | -2.69 | 0.01163 |
| LAMTOR1 | 5362.000 | 4987.000 | 0.9568 | -2673.000 | 0.01166 |
| SDHAF4 | 3044.000 | 2.64 | 0.8719 | -2615.000 | 0.01169 |
| TOR1AIP1 | 3445.000 | 3118.000 | 0.9194 | -2689.000 | 0.01186 |

| | | | | | |
|---|---|---|---|---|---|
| DIP2C | 2312.000 | 2649.000 | 1162.000 | 2673.000 | 0.01192 |
| LANCL2 | 2987.000 | 2.6 | 0.8732 | -2678.000 | 0.01195 |
| STARD3NL | 5447.000 | 5046.000 | 0.9548 | -2613.000 | 0.01196 |
| SSR4 | 6.92 | 6482.000 | 0.9662 | -2636.000 | 0.01196 |
| RHOD | 3743.000 | 3142.000 | 0.8674 | -2652.000 | 0.01197 |
| MRPL37 | 5518.000 | 5042.000 | 0.9472 | -2688.000 | 0.01206 |
| MCC | 1359.000 | 2044.000 | 2329.000 | 2.7 | 0.01209 |
| ATP6V1G1 | 7465.000 | 7052.000 | 0.9717 | -2646.000 | 0.01209 |
| MRAP | 0.1277 | 0.2409 | 0.6918 | 2721.000 | 0.01217 |
| C11orf58 | 6123.000 | 5859.000 | 0.9757 | -2637.000 | 0.0122 |
| SNX17 | 5446.000 | 5179.000 | 0.9703 | -2655.000 | 0.01221 |
| MAMDC4 | 2067.000 | 2498.000 | 1261.000 | 2626.000 | 0.01226 |
| TM9SF2 | 6307.000 | 5928.000 | 0.9663 | -2.64 | 0.01226 |
| PRTN3 | 0.8185 | 0.2406 | 7113.000 | -2549.000 | 0.01232 |
| FUCA1 | 5054.000 | 4582.000 | 0.9395 | -2657.000 | 0.01232 |
| RFLNA | 1341.000 | 0.8775 | -0.4455 | -2602.000 | 0.01235 |
| C11orf54 | 3303.000 | 2921.000 | 0.8972 | -2629.000 | 0.01237 |
| MAP3K12 | 1556.000 | 1904.000 | 1457.000 | 2.65 | 0.01241 |
| PRDX5 | 7216.000 | 6753.000 | 0.9665 | -2658.000 | 0.01245 |
| STXBP3 | 4164.000 | 3887.000 | 0.9518 | -2616.000 | 0.01277 |
| GNG5 | 6135.000 | 5541.000 | 0.9438 | -2648.000 | 0.01279 |
| ALG14 | 1243.000 | 1064.000 | 0.2861 | -2608.000 | 0.01286 |
| C3orf38 | 3343.000 | 3056.000 | 0.9255 | -2637.000 | 0.0129 |
| UBE2N | 4495.000 | 4135.000 | 0.9444 | -2604.000 | 0.01299 |
| ARHGAP33 | 2111.000 | 2547.000 | 1252.000 | 2592.000 | 0.01302 |
| TEX15 | 0.1006 | 0.02303 | 1642.000 | -2528.000 | 0.01302 |
| ICAM3 | 3.67 | 3248.000 | 0.906 | -2615.000 | 0.01302 |
| MRPS7 | 4459.000 | 4084.000 | 0.9413 | -2611.000 | 0.01304 |
| OSTM1 | 3.03 | 2663.000 | 0.8833 | -2634.000 | 0.01305 |
| ADSL | 3762.000 | 3505.000 | 0.9466 | -2574.000 | 0.01308 |
| GPR3 | 0.3338 | 0.7172 | 0.303 | 2668.000 | 0.01324 |
| APELA | 0.229 | 0.1005 | 1559.000 | -2528.000 | 0.0133 |
| DDX11 | 2924.000 | 3508.000 | 1.17 | 2625.000 | 0.01332 |
| FABP7 | 0.4023 | 0.04707 | 3356.000 | -2527.000 | 0.01334 |
| SYAP1 | 3941.000 | 3472.000 | 0.9076 | -2642.000 | 0.01347 |
| IRF5 | 2176.000 | 1.87 | 0.805 | -2595.000 | 0.0136 |
| SLC9A3R1 | 5875.000 | 5254.000 | 0.9369 | -2621.000 | 0.0136 |
| AFAP1L2 | 1873.000 | 2641.000 | 1548.000 | 2664.000 | 0.0137 |
| RPS6KL1 | 0.736 | 1239.000 | -0.6983 | 2614.000 | 0.01371 |
| FAM220A | 3529.000 | 3217.000 | 0.9266 | -2612.000 | 0.01371 |
| PPP1CA | 6553.000 | 6189.000 | 0.9696 | -2607.000 | 0.01375 |
| TSEN2 | 2088.000 | 2381.000 | 1178.000 | 2617.000 | 0.01384 |
| NMRK1 | 3102.000 | 2732.000 | 0.8877 | -2572.000 | 0.01385 |
| PRPF40B | 1.41 | 1.8 | 1713.000 | 2635.000 | 0.01388 |
| ATAD3A | 3522.000 | 3835.000 | 1068.000 | 2599.000 | 0.01394 |
| CTR9 | 4021.000 | 3739.000 | 0.9477 | -2596.000 | 0.01398 |

| | | | | | |
|---|---|---|---|---|---|
| TMEM199 | 3186.000 | 2.92 | 0.9249 | -2.58 | 0.01406 |
| UNC119 | 3053.000 | 2728.000 | 0.899 | -2608.000 | 0.01406 |
| KRBA1 | 1844.000 | 2.18 | 1274.000 | 2559.000 | 0.01407 |
| MUM1L1 | 0.842 | 0.4282 | 4.93 | -2536.000 | 0.01407 |
| POPDC3 | 0.5901 | 0.2959 | 2308.000 | -2497.000 | 0.01415 |
| RPL13 | 7.71 | 7345.000 | 0.9762 | -2561.000 | 0.01416 |
| SMIM19 | 3159.000 | 2829.000 | 0.9042 | -2521.000 | 0.01419 |
| NUDCD2 | 2443.000 | 2116.000 | 0.8394 | -2589.000 | 0.01432 |
| TOMM7 | 7.12 | 6692.000 | 0.9684 | -2595.000 | 0.01437 |
| ODAM | 0.8374 | 0.3366 | 6138.000 | -2491.000 | 0.01442 |
| AMMECR1 | 2249.000 | 1923.000 | 0.8066 | -2544.000 | 0.0145 |
| ERAP2 | 2508.000 | 1977.000 | 0.7415 | -2548.000 | 0.0145 |
| UBAC2 | 4651.000 | 4396.000 | 0.9633 | -2559.000 | 0.01451 |
| ROS1 | 0.1518 | 0.04637 | 1629.000 | -2486.000 | 0.01456 |
| CMTM6 | 5114.000 | 4553.000 | 0.9287 | -2586.000 | 0.0146 |
| IFI35 | 4341.000 | 3864.000 | 0.9207 | -2567.000 | 0.01471 |
| TBC1D8B | 1395.000 | 1087.000 | 0.2503 | -2554.000 | 0.01482 |
| PRSS48 | 0.0957 | 0.1462 | 0.8194 | 2585.000 | 0.01484 |
| MPO | 0.8385 | 0.2771 | 7288.000 | -2479.000 | 0.01484 |
| HAUS5 | 2489.000 | 2844.000 | 1146.000 | 2575.000 | 0.01487 |
| YAE1D1 | 3196.000 | 2932.000 | 0.9257 | -2527.000 | 0.01488 |
| C11orf71 | 2778.000 | 2239.000 | 0.789 | -2531.000 | 0.01492 |
| SRPK3 | 0.8542 | 1.21 | -1207.000 | 2592.000 | 0.01494 |
| USP17L2 | 0.01812 | 0.006637 | 1.25 | -2483.000 | 0.01494 |
| PSMB5 | 6206.000 | 5738.000 | 0.957 | -2598.000 | 0.01495 |
| BTD | 3506.000 | 3105.000 | 0.9032 | -2.55 | 0.01496 |
| BET1 | 3879.000 | 3545.000 | 0.9336 | -2555.000 | 0.01497 |
| C15orf62 | 0.8927 | 1.21 | -1678.000 | 2579.000 | 0.015 |
| GPR160 | 5204.000 | 4405.000 | 0.899 | -2576.000 | 0.01504 |
| UQCRB | 5.48 | 5.02 | 0.9484 | -2561.000 | 0.01509 |
| ZNF239 | 2203.000 | 2614.000 | 1217.000 | 2.58 | 0.01512 |
| SELENOH | 5629.000 | 5096.000 | 0.9425 | -2572.000 | 0.01528 |
| DNAJC10 | 2972.000 | 2572.000 | 0.8674 | -2546.000 | 0.01531 |
| COMMD6 | 4484.000 | 4192.000 | 0.9551 | -2529.000 | 0.01534 |
| APRT | 6505.000 | 5953.000 | 0.9526 | -2572.000 | 0.01536 |
| KCNJ14 | 0.7249 | 1015.000 | -0.04773 | 2592.000 | 0.01541 |
| SZT2 | 2343.000 | 2662.000 | 1.15 | 2563.000 | 0.01541 |
| TRDN | 0.3003 | 0.08724 | 2028.000 | -2464.000 | 0.01543 |
| COMMD8 | 3201.000 | 2782.000 | 0.8796 | -2532.000 | 0.01544 |
| SH3D21 | 2226.000 | 2678.000 | 1231.000 | 2554.000 | 0.01547 |
| BNIP1 | 3169.000 | 2.9 | 0.9232 | -2514.000 | 0.0155 |
| TCP11L2 | 1329.000 | 1099.000 | 0.3322 | -2536.000 | 0.0155 |
| IFNAR1 | 3978.000 | 3664.000 | 0.9403 | -2554.000 | 0.01559 |
| BAIAP2L1 | 4676.000 | 4333.000 | 0.9507 | -2523.000 | 0.01566 |
| PARP12 | 3527.000 | 3134.000 | 0.9062 | -2537.000 | 0.01566 |
| LY75 | 0.8765 | 0.6196 | 3631.000 | -2485.000 | 0.01574 |

| | | | | | |
|---|---|---|---|---|---|
| CDC14A | 2407.000 | 1914.000 | 0.7388 | -2502.000 | 0.01576 |
| RSBN1 | 2.48 | 2237.000 | 0.8862 | -2547.000 | 0.01578 |
| HIST1H4F | 0.314 | 0.1397 | 1699.000 | -2452.000 | 0.016 |
| SH3GLB1 | 3706.000 | 3353.000 | 0.9235 | -2551.000 | 0.01601 |
| RHOC | 6232.000 | 5806.000 | 0.9613 | -2546.000 | 0.01605 |
| UCHL5 | 3.22 | 2946.000 | 0.9239 | -2.51 | 0.0161 |
| NPHP4 | 2008.000 | 2277.000 | 1.18 | 2526.000 | 0.01613 |
| C2orf72 | 2177.000 | 1.56 | 0.5717 | -2484.000 | 0.01621 |
| RWDD1 | 3347.000 | 2922.000 | 0.8875 | -2546.000 | 0.01623 |
| C12orf45 | 0.921 | 0.741 | 3641.000 | -2523.000 | 0.01626 |
| SETD5 | 4364.000 | 4671.000 | 1046.000 | 2532.000 | 0.0163 |
| DDX60 | 2628.000 | 2108.000 | 0.7721 | -2.52 | 0.01632 |
| ATRIP | 2443.000 | 2649.000 | 1091.000 | 2514.000 | 0.01639 |
| COX7B | 5618.000 | 5226.000 | 0.9581 | -2495.000 | 0.01646 |
| IL1RAPL1 | 0.4924 | 0.2598 | 1903.000 | -2.46 | 0.01651 |
| RNASE3 | 0.6788 | 0.2142 | 3978.000 | -2439.000 | 0.01655 |
| ARCN1 | 5558.000 | 5227.000 | 0.9642 | -2517.000 | 0.01661 |
| PSMA1 | 5671.000 | 5335.000 | 0.9647 | -2509.000 | 0.0167 |
| KIAA0907 | 3773.000 | 4141.000 | 1.07 | 2.54 | 0.01672 |
| ATG9B | 0.7872 | 1227.000 | -0.854 | 2547.000 | 0.01679 |
| AP2S1 | 5932.000 | 5.49 | 0.9566 | -2548.000 | 0.0168 |
| APOL3 | 2759.000 | 2.28 | 0.8121 | -2497.000 | 0.01681 |
| MERTK | 2564.000 | 2164.000 | 0.8197 | -2522.000 | 0.01685 |
| FBXO9 | 3977.000 | 3623.000 | 0.9323 | -2518.000 | 0.01687 |
| SLC30A1 | 3041.000 | 2732.000 | 0.9036 | -2507.000 | 0.01692 |
| CDC42EP3 | 3247.000 | 2814.000 | 0.8786 | -2542.000 | 0.01694 |
| PLAC8 | 0.7841 | 0.4504 | 3279.000 | -2.45 | 0.01695 |
| TBCC | 4113.000 | 3.72 | 0.9291 | -2541.000 | 0.01696 |
| NANS | 6242.000 | 5718.000 | 0.9521 | -2529.000 | 0.01697 |
| SCGB2B2 | 0.5156 | 0.7594 | 0.4156 | 2513.000 | 0.01701 |
| MOB1B | 2818.000 | 2542.000 | 0.9004 | -2488.000 | 0.01705 |
| LAMTOR4 | 6.02 | 5652.000 | 0.9649 | -2515.000 | 0.01706 |
| CAMTA2 | 3641.000 | 3922.000 | 1058.000 | 2501.000 | 0.0171 |
| CAMP | 1263.000 | 0.6157 | -2078.000 | -2431.000 | 0.01712 |
| DHX35 | 2.29 | 2.56 | 1135.000 | 2533.000 | 0.01715 |
| LRIF1 | 2974.000 | 2658.000 | 0.8972 | -2.52 | 0.01716 |
| SORL1 | 3559.000 | 3061.000 | 0.8813 | -2507.000 | 0.01719 |
| ARHGEF39 | 1382.000 | 1789.000 | 1797.000 | 2508.000 | 0.01722 |
| FAIM | 3455.000 | 3111.000 | 0.9156 | -2478.000 | 0.01725 |
| NHP2 | 6142.000 | 5756.000 | 0.9643 | -2506.000 | 0.01725 |
| EIF2A | 5645.000 | 5286.000 | 0.9621 | -2498.000 | 0.01739 |
| KRI1 | 3844.000 | 4178.000 | 1062.000 | 2515.000 | 0.01752 |
| CEP72 | 1631.000 | 1925.000 | 1339.000 | 2493.000 | 0.01754 |
| NDUFB5 | 4229.000 | 3794.000 | 0.9248 | -2.49 | 0.01759 |
| ADRA2A | 2058.000 | 2.99 | 1517.000 | 2506.000 | 0.0176 |
| PSMA5 | 4181.000 | 3893.000 | 0.9502 | -2462.000 | 0.01762 |

| | | | | | |
|---|---|---|---|---|---|
| ZNF620 | 1328.000 | 1.65 | 1765.000 | 2515.000 | 0.01769 |
| TOMM6 | 7371.000 | 7147.000 | 0.9846 | -2.43 | 0.01781 |
| AHRR | 0.381 | 0.599 | 0.5311 | 2508.000 | 0.01783 |
| MAN1A2 | 3.09 | 2839.000 | 0.9248 | -2489.000 | 0.01784 |
| IL7 | 0.8893 | 0.6304 | 3933.000 | -2446.000 | 0.0179 |
| CFAP74 | 0.169 | 0.5431 | 0.3434 | 2542.000 | 0.01791 |
| RNF7 | 4604.000 | 4.25 | 0.9475 | -2462.000 | 0.01792 |
| ABHD4 | 3644.000 | 3205.000 | 0.9007 | -2.51 | 0.01798 |
| CCDC114 | 0.4954 | 0.8665 | 0.204 | 2498.000 | 0.018 |
| SLITRK4 | 0.6751 | 0.3553 | 2634.000 | -2.43 | 0.01801 |
| TRPV1 | 1494.000 | 1747.000 | 1391.000 | 2484.000 | 0.01802 |
| CCDC83 | 0.2594 | 0.1475 | 1418.000 | -2414.000 | 0.01803 |
| ALKBH7 | 4809.000 | 4336.000 | 0.9341 | -2479.000 | 0.01804 |
| NMNAT3 | 1634.000 | 2083.000 | 1495.000 | 2468.000 | 0.01814 |
| TEN1 | 4359.000 | 4018.000 | 0.9446 | -2488.000 | 0.01815 |
| TMEM139 | 0.4407 | 0.8202 | 0.2419 | 2525.000 | 0.01817 |
| DEFA4 | 0.8159 | 0.2729 | 6382.000 | -2401.000 | 0.01818 |
| UBE2D4 | 3.01 | 2732.000 | 0.9121 | -2494.000 | 0.01819 |
| HINT3 | 3.9 | 3352.000 | 0.8888 | -2513.000 | 0.0182 |
| ARHGEF11 | 3362.000 | 3602.000 | 1057.000 | 2467.000 | 0.01821 |
| NDFIP1 | 5601.000 | 5274.000 | 0.9651 | -2.47 | 0.01825 |
| CENPT | 3.8 | 4222.000 | 1079.000 | 2.48 | 0.01831 |
| AKAP7 | 1813.000 | 1415.000 | 0.5832 | -2491.000 | 0.01835 |
| FBXO18 | 4.27 | 4544.000 | 1043.000 | 2485.000 | 0.01838 |
| SAP18 | 5649.000 | 5.24 | 0.9567 | -2483.000 | 0.01839 |
| AURKC | 0.6703 | 0.9555 | 0.1137 | 2514.000 | 0.01843 |
| PPP2R3C | 3297.000 | 2994.000 | 0.919 | -2497.000 | 0.01844 |
| ATAD3B | 2902.000 | 3.33 | 1129.000 | 2463.000 | 0.01847 |
| ANXA3 | 3784.000 | 2797.000 | 0.7728 | -2.48 | 0.01847 |
| HIST1H3G | 0.1693 | 0.05146 | 1671.000 | -2397.000 | 0.01857 |
| TMEM50A | 5586.000 | 5171.000 | 0.9551 | -2492.000 | 0.01857 |
| CCDC189 | 0.6934 | 1014.000 | -0.03819 | 2505.000 | 0.01863 |
| FANCC | 2047.000 | 2403.000 | 1223.000 | 2496.000 | 0.01868 |
| PPP1CB | 6254.000 | 5954.000 | 0.9731 | -2464.000 | 0.01876 |
| CORO2A | 3608.000 | 3102.000 | 0.8822 | -2475.000 | 0.01877 |
| MYH3 | 0.679 | 0.9961 | 0.01009 | 2492.000 | 0.01881 |
| ACKR3 | 2877.000 | 3639.000 | 1223.000 | 2499.000 | 0.01884 |
| DESI1 | 3838.000 | 3461.000 | 0.9231 | -2472.000 | 0.01888 |
| SLC9A2 | 2.29 | 1.67 | 0.6188 | -2492.000 | 0.019 |
| LYRM1 | 3911.000 | 3548.000 | 0.9286 | -2449.000 | 0.0191 |
| SIAE | 2969.000 | 2524.000 | 0.8507 | -2478.000 | 0.01912 |
| SPAG16 | 2396.000 | 1978.000 | 0.7805 | -2459.000 | 0.01917 |
| MRPL49 | 5261.000 | 4909.000 | 0.9583 | -2446.000 | 0.01933 |
| CDV3 | 4993.000 | 4527.000 | 0.939 | -2.48 | 0.01933 |
| ARL14EP | 3.81 | 3581.000 | 0.9538 | -2421.000 | 0.01939 |
| TMEM256 | 6043.000 | 5584.000 | 0.9561 | -2428.000 | 0.01944 |

| | | | | | |
|---|---|---|---|---|---|
| USO1 | 4875.000 | 4528.000 | 0.9533 | -2.46 | 0.01946 |
| UBQLN1 | 5182.000 | 4833.000 | 0.9577 | -2.46 | 0.01946 |
| SAMD10 | 2905.000 | 3235.000 | 1101.000 | 2438.000 | 0.0195 |
| BTBD1 | 4419.000 | 4153.000 | 0.9582 | -2447.000 | 0.01955 |
| CTD-2659N19 | 0.2677 | 0.4146 | 0.6681 | 2478.000 | 0.01972 |
| CES4A | 1103.000 | 1745.000 | 5697.000 | 2.45 | 0.01979 |
| C9orf85 | 2018.000 | 1787.000 | 0.8268 | -2424.000 | 0.01979 |
| HIGD2A | 6176.000 | 5738.000 | 0.9596 | -2431.000 | 0.01984 |
| HSH2D | 2696.000 | 2224.000 | 0.806 | -2414.000 | 0.0199 |
| BRF1 | 3197.000 | 3425.000 | 1059.000 | 2447.000 | 0.02002 |
| IL18R1 | 0.7967 | 0.5895 | 2326.000 | -2398.000 | 0.02002 |
| IDNK | 2483.000 | 2202.000 | 0.8681 | -2434.000 | 0.02008 |
| YPEL2 | 3048.000 | 2713.000 | 0.8956 | -2437.000 | 0.02013 |
| ALDH1A2 | 0.2978 | 0.7189 | 0.2724 | 2498.000 | 0.02014 |
| CAPRIN1 | 5686.000 | 5385.000 | 0.9687 | -2435.000 | 0.02014 |
| ATG4A | 3093.000 | 2865.000 | 0.9321 | -2401.000 | 0.02015 |
| FUT4 | 1.37 | 1104.000 | 0.3141 | -2384.000 | 0.02018 |
| BCL2L2 | 4341.000 | 3964.000 | 0.9382 | -2451.000 | 0.02019 |
| CRACR2A | 1.15 | 0.768 | -1892.000 | -2408.000 | 0.02024 |
| ALKBH6 | 3111.000 | 3435.000 | 1087.000 | 2451.000 | 0.02028 |
| SH3BGRL2 | 2.93 | 2.36 | 0.799 | -2427.000 | 0.02029 |
| CYB5A | 4449.000 | 3806.000 | 0.8954 | -2454.000 | 0.02037 |
| UCKL1 | 4207.000 | 4547.000 | 1054.000 | 2443.000 | 0.02039 |
| EGR2 | 1214.000 | 1811.000 | 3061.000 | 2452.000 | 0.02046 |
| IKZF1 | 1143.000 | 0.8321 | -1378.000 | -2372.000 | 0.02049 |
| IL10RB | 4488.000 | 4.13 | 0.9446 | -2448.000 | 0.02049 |
| TAS2R3 | 0.1398 | 0.236 | 0.7338 | 2475.000 | 0.0205 |
| UBE2V2 | 3711.000 | 3203.000 | 0.8879 | -2455.000 | 0.02053 |
| PPP1R7 | 4348.000 | 4038.000 | 0.9498 | -2438.000 | 0.02055 |
| AMIGO2 | 2041.000 | 1554.000 | 0.6176 | -2407.000 | 0.02056 |
| DR1 | 2921.000 | 2656.000 | 0.9112 | -2435.000 | 0.02057 |
| MS4A3 | 0.4267 | 0.117 | 2519.000 | -2351.000 | 0.02066 |
| MBNL1 | 4192.000 | 3826.000 | 0.9362 | -2441.000 | 0.02068 |
| MGAT2 | 4.28 | 3.89 | 0.9343 | -2461.000 | 0.02071 |
| FAM120AOS | 4493.000 | 4242.000 | 0.9617 | -2406.000 | 0.02072 |
| HPS3 | 3.02 | 2674.000 | 0.8901 | -2441.000 | 0.02082 |
| SRPRB | 5194.000 | 4739.000 | 0.9443 | -2428.000 | 0.02089 |
| ANKRD6 | 1496.000 | 1231.000 | 0.515 | -2421.000 | 0.0209 |
| ARPC1B | 5631.000 | 5203.000 | 0.9543 | -2421.000 | 0.02103 |
| SPON2 | 6.4 | 7788.000 | 1106.000 | 2417.000 | 0.02107 |
| IQSEC3 | 0.2884 | 0.5705 | 0.4514 | 2461.000 | 0.02114 |
| MDH2 | 6487.000 | 6071.000 | 0.9645 | -2437.000 | 0.02122 |
| INPP4A | 2071.000 | 2324.000 | 1158.000 | 2.41 | 0.02128 |
| USP16 | 4193.000 | 3885.000 | 0.9468 | -2.43 | 0.02133 |
| GTF3A | 6103.000 | 5.74 | 0.9661 | -2412.000 | 0.02138 |
| THBS3 | 3435.000 | 3893.000 | 1102.000 | 2418.000 | 0.02146 |

| | | | | | |
|---|---|---|---|---|---|
| ZNF483 | 0.3229 | 0.4352 | 0.736 | 2419.000 | 0.02149 |
| HMBS | 3891.000 | 3.53 | 0.9284 | -2367.000 | 0.02149 |
| ZNF860 | 1359.000 | 1044.000 | 0.1417 | -2388.000 | 0.0217 |
| OGG1 | 2784.000 | 3112.000 | 1109.000 | 2437.000 | 0.02177 |
| MBTPS1 | 4735.000 | 4518.000 | 0.9698 | -2388.000 | 0.0218 |
| ATP5F1 | 5624.000 | 5.31 | 0.9667 | -2397.000 | 0.02189 |
| TM2D2 | 3638.000 | 3297.000 | 0.9237 | -2386.000 | 0.02203 |
| PEG3 | 0.6964 | 1.03 | -0.08082 | 2436.000 | 0.02216 |
| INPP5F | 1831.000 | 1988.000 | 1136.000 | 2359.000 | 0.02217 |
| NDUFC2 | 5515.000 | 5055.000 | 0.949 | -2406.000 | 0.02222 |
| CARNMT1 | 2566.000 | 2333.000 | 0.8988 | -2387.000 | 0.0223 |
| FAM134C | 4489.000 | 4219.000 | 0.9588 | -2396.000 | 0.02235 |
| LUZP2 | 1892.000 | 1131.000 | 0.1931 | -2385.000 | 0.02236 |
| PDE1A | 1311.000 | 0.9441 | -0.2125 | -2.37 | 0.02238 |
| CFAP45 | 0.8384 | 1331.000 | -1621.000 | 2406.000 | 0.02241 |
| PTPN22 | 0.9292 | 0.6262 | 6375.000 | -2356.000 | 0.02243 |
| DEFA3 | 0.7188 | 0.2536 | 4155.000 | -2324.000 | 0.02248 |
| STK17B | 2.99 | 2517.000 | 0.8426 | -2396.000 | 0.02266 |
| OAS2 | 2978.000 | 2298.000 | 0.7625 | -2.37 | 0.02268 |
| TRIAP1 | 4496.000 | 3859.000 | 0.8983 | -2414.000 | 0.02281 |
| IDO1 | 1218.000 | 0.7528 | -1443.000 | -2334.000 | 0.02283 |
| PARP6 | 2307.000 | 3106.000 | 1356.000 | 2414.000 | 0.02287 |
| PRH1 | 0.8465 | 1239.000 | -1284.000 | 2422.000 | 0.02299 |
| PIN4 | 3062.000 | 2742.000 | 0.9014 | -2375.000 | 0.02299 |
| CRIPAK | 2121.000 | 2.84 | 1388.000 | 2422.000 | 0.02301 |
| IVNS1ABP | 4461.000 | 4.8 | 1049.000 | 2371.000 | 0.02302 |
| MMADHC | 5296.000 | 4932.000 | 0.9573 | -2387.000 | 0.02311 |
| WSB2 | 5409.000 | 4959.000 | 0.9485 | -2405.000 | 0.02311 |
| TMED5 | 3499.000 | 3137.000 | 0.9128 | -2379.000 | 0.02312 |
| TMEM5 | 3615.000 | 3227.000 | 0.9114 | -2.4 | 0.02315 |
| HIGD1B | 1736.000 | 2114.000 | 1357.000 | 2389.000 | 0.02317 |
| HLA-C | 8623.000 | 8101.000 | 0.971 | -2.39 | 0.02317 |
| RBX1 | 5493.000 | 5216.000 | 0.9696 | -2343.000 | 0.02323 |
| TMED2 | 6664.000 | 6256.000 | 0.9667 | -2386.000 | 0.02323 |
| ARFIP1 | 3757.000 | 3.45 | 0.9355 | -2376.000 | 0.02325 |
| SLC4A1AP | 3559.000 | 3301.000 | 0.9407 | -2412.000 | 0.02325 |
| BLOC1S1 | 4949.000 | 4596.000 | 0.9537 | -2376.000 | 0.02331 |
| STK38L | 3163.000 | 2791.000 | 0.8913 | -2.38 | 0.02347 |
| C16orf52 | 1964.000 | 2195.000 | 1165.000 | 2.37 | 0.02351 |
| RTEL1 | 2561.000 | 2845.000 | 1112.000 | 2355.000 | 0.02351 |
| TRAK1 | 2568.000 | 2.81 | 1096.000 | 2369.000 | 0.02352 |
| COX17 | 5408.000 | 4942.000 | 0.9467 | -2384.000 | 0.02359 |
| SDHB | 5794.000 | 5408.000 | 0.9608 | -2383.000 | 0.0236 |
| FGF17 | 0.2957 | 0.5289 | 0.5228 | 2404.000 | 0.02363 |
| HAUS7 | 0.2113 | 0.3296 | 0.7142 | 2.38 | 0.02363 |
| C8orf88 | 0.7999 | 0.527 | 2868.000 | -2329.000 | 0.02366 |

| | | | | | |
|---|---|---|---|---|---|
| PAPD5 | 2604.000 | 2301.000 | 0.8709 | -2366.000 | 0.02374 |
| PCMTD1 | 4089.000 | 3661.000 | 0.9216 | -2376.000 | 0.02374 |
| CHCHD7 | 3795.000 | 3439.000 | 0.926 | -2339.000 | 0.02376 |
| ZDHHC6 | 3555.000 | 3805.000 | 1054.000 | 2353.000 | 0.02377 |
| CHTF8 | 4432.000 | 4.13 | 0.9525 | -2375.000 | 0.02379 |
| CAPZA2 | 4317.000 | 3954.000 | 0.9399 | -2363.000 | 0.02383 |
| TRIM43 | 0.005744 | 0.0009175 | 1356.000 | -2294.000 | 0.02384 |
| MAPK8IP3 | 3.51 | 3895.000 | 1083.000 | 2341.000 | 0.0239 |
| SDHAF3 | 3227.000 | 2803.000 | 0.8799 | -2368.000 | 0.02391 |
| IL1R1 | 4252.000 | 3.63 | 0.8906 | -2366.000 | 0.02392 |
| SULT1B1 | 0.5053 | 0.2921 | 1803.000 | -2304.000 | 0.02395 |
| IFIH1 | 2889.000 | 2496.000 | 0.862 | -2355.000 | 0.02418 |
| KITLG | 1.93 | 2483.000 | 1383.000 | 2397.000 | 0.02419 |
| CRYZ | 3352.000 | 2952.000 | 0.895 | -2337.000 | 0.02421 |
| BIRC2 | 3815.000 | 3488.000 | 0.9332 | -2357.000 | 0.02421 |
| JCHAIN | 4324.000 | 3144.000 | 0.7824 | -2363.000 | 0.02425 |
| CPT1C | 0.9171 | 1512.000 | -4.78 | 2398.000 | 0.02427 |
| APAF1 | 2001.000 | 1769.000 | 0.822 | -2353.000 | 0.02427 |
| C1orf53 | 2333.000 | 1844.000 | 0.7224 | -2344.000 | 0.02433 |
| SPTY2D1 | 3102.000 | 2873.000 | 0.9323 | -2343.000 | 0.02434 |
| COMMD9 | 3498.000 | 3227.000 | 0.9355 | -2362.000 | 0.02442 |
| DPM3 | 5826.000 | 5244.000 | 0.9403 | -2.37 | 0.02443 |
| NKX2-2 | 0.844 | 0.4685 | 4.47 | -2317.000 | 0.02444 |
| MRPL15 | 5546.000 | 5029.000 | 0.9429 | -2362.000 | 0.02444 |
| ETV4 | 0.6956 | 1876.000 | -1734.000 | 2408.000 | 0.02455 |
| OPTN | 3725.000 | 3.28 | 0.9033 | -2363.000 | 0.02456 |
| DBNL | 5622.000 | 5359.000 | 0.9723 | -2359.000 | 0.0246 |
| TAS2R5 | 0.7378 | 1.1 | -0.3135 | 2386.000 | 0.02461 |
| CNDP2 | 5151.000 | 4741.000 | 0.9494 | -2.36 | 0.02466 |
| SCP2 | 4322.000 | 3937.000 | 0.9363 | -2359.000 | 0.02474 |
| ZNF266 | 3202.000 | 3492.000 | 1075.000 | 2352.000 | 0.02485 |
| ZC3HAV1L | 2187.000 | 2.57 | 1206.000 | 2349.000 | 0.02485 |
| CD59 | 6.04 | 5697.000 | 0.9674 | -2332.000 | 0.02486 |
| STARD9 | 0.5599 | 0.8933 | 0.1945 | 2.39 | 0.02501 |
| ASL | 4145.000 | 3818.000 | 0.9422 | -2337.000 | 0.02501 |
| SEC62 | 5212.000 | 4805.000 | 0.9507 | -2367.000 | 0.02504 |
| RNASE2 | 1011.000 | 0.557 | -55.96 | -2279.000 | 0.02506 |
| THAP6 | 2.03 | 1808.000 | 0.836 | -2348.000 | 0.02506 |
| C9orf24 | 0.3639 | 0.6317 | 0.4544 | 2359.000 | 0.02507 |
| MOV10 | 4013.000 | 3727.000 | 0.9468 | -2328.000 | 0.02511 |
| ICK | 3149.000 | 2782.000 | 0.8918 | -2338.000 | 0.02517 |
| CEP128 | 1147.000 | 1416.000 | 2536.000 | 2365.000 | 0.02519 |
| SLC9A5 | 0.843 | 1219.000 | -1.16 | 2361.000 | 0.02526 |
| ARSB | 1814.000 | 1.52 | 0.7033 | -2351.000 | 0.02529 |
| PRR26 | 0.1879 | 0.3009 | 0.7183 | 2375.000 | 0.02531 |
| TAMM41 | 2349.000 | 2611.000 | 1124.000 | 2365.000 | 0.02541 |

| | | | | | |
|---|---|---|---|---|---|
| NDUFB7 | 7348.000 | 6981.000 | 0.9743 | -2331.000 | 0.02541 |
| BPGM | 3632.000 | 3319.000 | 0.9301 | -2323.000 | 0.02546 |
| MTERF1 | 2084.000 | 1827.000 | 0.8207 | -2342.000 | 0.02548 |
| LINC00493 | 5105.000 | 4473.000 | 0.9189 | -2381.000 | 0.02552 |
| GPR137C | 1005.000 | 1479.000 | 78.74 | 2367.000 | 0.0256 |
| RABIF | 3375.000 | 3084.000 | 0.9259 | -2323.000 | 0.02566 |
| C15orf65 | 2343.000 | 1922.000 | 0.7674 | -2347.000 | 0.02587 |
| TAP1 | 4917.000 | 4384.000 | 0.9279 | -2331.000 | 0.02593 |
| BIK | 4167.000 | 3731.000 | 0.9226 | -2319.000 | 0.02594 |
| RPS27L | 2972.000 | 2625.000 | 0.8859 | -2329.000 | 0.02598 |
| CTRL | 0.9074 | 1138.000 | -1334.000 | 2331.000 | 0.02603 |
| ARL16 | 4313.000 | 3977.000 | 0.9444 | -2333.000 | 0.02603 |
| SAMM50 | 4685.000 | 4459.000 | 0.968 | -2318.000 | 0.02605 |
| FOXD4 | 0.6584 | 1007.000 | -0.01586 | 2341.000 | 0.02612 |
| TDO2 | 1961.000 | 1.2 | 0.2711 | -2275.000 | 0.0262 |
| SF3A3 | 4471.000 | 4698.000 | 1033.000 | 2341.000 | 0.02623 |
| POLB | 4314.000 | 3983.000 | 0.9454 | -2273.000 | 0.02623 |
| CINP | 4878.000 | 4597.000 | 0.9626 | -2319.000 | 0.02627 |
| VANGL1 | 2545.000 | 2209.000 | 0.8482 | -2334.000 | 0.02632 |
| NEK7 | 3898.000 | 3555.000 | 0.9323 | -2327.000 | 0.02634 |
| WDR35 | 1949.000 | 2221.000 | 1196.000 | 2334.000 | 0.02638 |
| ZNF589 | 2.77 | 3183.000 | 1136.000 | 2331.000 | 0.02638 |
| DPP4 | 3181.000 | 2347.000 | 0.7372 | -2314.000 | 0.02638 |
| TMEM88 | 2001.000 | 2357.000 | 1236.000 | 2.32 | 0.0264 |
| MRPS24 | 6015.000 | 5546.000 | 0.9548 | -2341.000 | 0.02643 |
| DRD4 | 0.6169 | 0.8984 | 0.2218 | 2332.000 | 0.02644 |
| CYP20A1 | 3.16 | 2921.000 | 0.9317 | -2335.000 | 0.02644 |
| TMEM248 | 5213.000 | 4788.000 | 0.9485 | -2.33 | 0.02645 |
| TMEM68 | 2.4 | 2046.000 | 0.8177 | -2312.000 | 0.02648 |
| MORN1 | 2052.000 | 2342.000 | 1184.000 | 2322.000 | 0.02649 |
| NUDT2 | 4309.000 | 3871.000 | 0.9265 | -2323.000 | 0.02653 |
| RMI1 | 3.26 | 2846.000 | 0.8849 | -2321.000 | 0.02655 |
| SH3BGRL3 | 5696.000 | 5292.000 | 0.9577 | -2342.000 | 0.02661 |
| NDUFA4 | 6393.000 | 6111.000 | 0.9756 | -2278.000 | 0.02667 |
| HBD | 1297.000 | 0.6403 | -1716.000 | -2253.000 | 0.02675 |
| SDHAF2 | 4173.000 | 3.94 | 0.9598 | -2312.000 | 0.02676 |
| GPRIN3 | 0.6801 | 0.5097 | 1748.000 | -2.26 | 0.02677 |
| ZNF232 | 2438.000 | 2.73 | 1127.000 | 2322.000 | 0.02693 |
| NPR2 | 1659.000 | 2065.000 | 1432.000 | 2334.000 | 0.02713 |
| CBX2 | 2321.000 | 1822.000 | 0.7125 | -2292.000 | 0.02719 |
| GAPDHS | 1881.000 | 2213.000 | 1258.000 | 2314.000 | 0.0272 |
| ITGA7 | 1886.000 | 2421.000 | 1394.000 | 2332.000 | 0.02722 |
| COX8A | 8672.000 | 8.28 | 0.9786 | -2313.000 | 0.0273 |
| EIF4E2 | 3827.000 | 3.65 | 0.9646 | -2294.000 | 0.02731 |
| TUBG2 | 3171.000 | 3516.000 | 1089.000 | 2295.000 | 0.02732 |
| COPS4 | 3846.000 | 3559.000 | 0.9426 | -2304.000 | 0.02736 |

| | | | | | |
|---|---|---|---|---|---|
| NECAP1 | 3802.000 | 3481.000 | 0.9339 | -2312.000 | 0.02749 |
| C11orf68 | 4548.000 | 4099.000 | 0.9314 | -2339.000 | 0.02749 |
| ZNF558 | 1826.000 | 2048.000 | 1.19 | 2305.000 | 0.02752 |
| YIPF5 | 4146.000 | 3.89 | 0.9552 | -2292.000 | 0.02759 |
| TAB3 | 3314.000 | 2.97 | 0.9087 | -2289.000 | 0.02767 |
| RARS | 4904.000 | 4629.000 | 0.9637 | -2306.000 | 0.02771 |
| NDUFV2 | 6071.000 | 5565.000 | 0.9517 | -2319.000 | 0.02771 |
| UQCC1 | 2973.000 | 3221.000 | 1073.000 | 2301.000 | 0.02772 |
| ABRACL | 4302.000 | 3814.000 | 0.9175 | -2.29 | 0.02785 |
| BCAP29 | 3594.000 | 3.17 | 0.9018 | -2.3 | 0.02787 |
| MRPL41 | 5925.000 | 5376.000 | 0.9453 | -2325.000 | 0.02796 |
| GON4L | 2662.000 | 2877.000 | 1079.000 | 2302.000 | 0.02799 |
| PRH2 | 0.1418 | 0.2212 | 0.7725 | 2329.000 | 0.02805 |
| TIMMDC1 | 5004.000 | 4739.000 | 0.9663 | -2269.000 | 0.02814 |
| HIST1H4I | 4297.000 | 3656.000 | 0.8892 | -2278.000 | 0.02814 |
| UHRF1BP1L | 2663.000 | 2389.000 | 0.8888 | -2313.000 | 0.02814 |
| RWDD2B | 2.81 | 2517.000 | 0.8931 | -2296.000 | 0.02816 |
| THTPA | 3488.000 | 3.22 | 0.9361 | -2278.000 | 0.02825 |
| NDUFB6 | 4786.000 | 4.33 | 0.9361 | -2306.000 | 0.02846 |
| CEP131 | 2807.000 | 3079.000 | 1.09 | 2261.000 | 0.02863 |
| ARFGAP1 | 4674.000 | 4.94 | 1036.000 | 2279.000 | 0.02865 |
| CIB1 | 6191.000 | 5799.000 | 0.9641 | -2286.000 | 0.02889 |
| PNPLA4 | 3025.000 | 2644.000 | 0.8784 | -2.27 | 0.02906 |
| TSSK3 | 1685.000 | 2.05 | 1376.000 | 2296.000 | 0.02909 |
| RPS6KC1 | 3102.000 | 2716.000 | 0.8829 | -2249.000 | 0.02909 |
| MIER1 | 2805.000 | 2596.000 | 0.9249 | -2286.000 | 0.02915 |
| POP4 | 2973.000 | 2.72 | 0.9182 | -2272.000 | 0.02921 |
| SDCBP | 5386.000 | 4899.000 | 0.9437 | -2301.000 | 0.02921 |
| MCAM | 4528.000 | 5.04 | 1071.000 | 2268.000 | 0.02923 |
| COX6A1 | 8834.000 | 8307.000 | 0.9718 | -2294.000 | 0.02923 |
| DZIP1L | 0.9654 | 1369.000 | -8929.000 | 2.3 | 0.02927 |
| UBL3 | 3653.000 | 3331.000 | 0.9288 | -2268.000 | 0.02936 |
| IGLL5 | 5192.000 | 3903.000 | 0.8268 | -2274.000 | 0.02941 |
| PEX13 | 2849.000 | 2605.000 | 0.9145 | -2272.000 | 0.02942 |
| MED10 | 4481.000 | 4211.000 | 0.9586 | -2291.000 | 0.0296 |
| DUSP19 | 0.6435 | 0.4889 | 1623.000 | -2265.000 | 0.02964 |
| HIF1AN | 2423.000 | 2127.000 | 0.8531 | -2279.000 | 0.02967 |
| BLVRB | 5961.000 | 5473.000 | 0.9521 | -2261.000 | 0.02973 |
| STAG3 | 1015.000 | 1374.000 | 21.39 | 2306.000 | 0.02975 |
| SSSCA1 | 3892.000 | 3658.000 | 0.9543 | -2266.000 | 0.02977 |
| GTF3C2 | 4023.000 | 3802.000 | 0.9593 | -2288.000 | 0.02979 |
| CDK5RAP2 | 3.53 | 3802.000 | 1059.000 | 2279.000 | 0.02981 |
| L3MBTL1 | 1278.000 | 1586.000 | 1.88 | 2279.000 | 0.02983 |
| GTF2H5 | 2522.000 | 2.12 | 0.8123 | -2.28 | 0.02984 |
| ARL6IP5 | 6201.000 | 5696.000 | 0.9534 | -2294.000 | 0.02987 |
| CUX2 | 2062.000 | 1551.000 | 0.606 | -2241.000 | 0.02991 |

| | | | | | |
|---|---|---|---|---|---|
| COLQ | 1084.000 | 1465.000 | 4716.000 | 2288.000 | 0.02994 |
| TCN1 | 0.6966 | 0.2967 | 3361.000 | -2.21 | 0.02997 |
| COPB1 | 5691.000 | 5403.000 | 0.9701 | -2251.000 | 0.02998 |
| TERF2IP | 5.18 | 4827.000 | 0.9571 | -2274.000 | 0.03005 |
| ZNF750 | 0.522 | 0.8587 | 0.2343 | 2305.000 | 0.03006 |
| TNFRSF6B | 1992.000 | 2355.000 | 1243.000 | 2258.000 | 0.03011 |
| THNSL1 | 2078.000 | 1684.000 | 0.712 | -2264.000 | 0.0302 |
| KANSL3 | 3813.000 | 4.02 | 1.04 | 2281.000 | 0.03025 |
| ARF4 | 7142.000 | 6765.000 | 0.9724 | -2276.000 | 0.03032 |
| ZNF442 | 0.5769 | 0.7092 | 0.6248 | 2275.000 | 0.03037 |
| NDUFA6 | 5425.000 | 5112.000 | 0.9648 | -2254.000 | 0.03045 |
| CDC5L | 3438.000 | 3185.000 | 0.9381 | -2258.000 | 0.03045 |
| CMTM2 | 0.5921 | 0.8149 | 0.3906 | 2.25 | 0.03046 |
| NDUFC1 | 5049.000 | 4684.000 | 0.9537 | -2.27 | 0.03062 |
| PHACTR2 | 2002.000 | 1724.000 | 0.785 | -2.25 | 0.03081 |
| CDAN1 | 2399.000 | 2668.000 | 1122.000 | 2247.000 | 0.03086 |
| TJP2 | 3402.000 | 3046.000 | 0.9098 | -2.24 | 0.03087 |
| AIMP1 | 4244.000 | 3.93 | 0.9469 | -2251.000 | 0.03089 |
| MAT2B | 3836.000 | 3559.000 | 0.9443 | -2239.000 | 0.03098 |
| LRRTM2 | 0.09798 | 0.1312 | 0.8744 | 2262.000 | 0.03099 |
| OAS3 | 3215.000 | 2635.000 | 0.8295 | -2254.000 | 0.031 |
| PKIA | 2772.000 | 2.17 | 0.76 | -2.26 | 0.03112 |
| NOA1 | 3.56 | 3223.000 | 0.9218 | -2256.000 | 0.03114 |
| LSMEM2 | 0.1282 | 0.1969 | 0.791 | 2268.000 | 0.03115 |
| OXA1L | 5066.000 | 4748.000 | 0.96 | -2.27 | 0.03116 |
| RNF11 | 5232.000 | 4878.000 | 0.9577 | -2264.000 | 0.03122 |
| SEMA4C | 3398.000 | 3708.000 | 1071.000 | 2252.000 | 0.03125 |
| NR2C2 | 3015.000 | 3214.000 | 1058.000 | 2253.000 | 0.0313 |
| MRPL3 | 6097.000 | 5832.000 | 0.9755 | -2232.000 | 0.03136 |
| CHMP5 | 5381.000 | 5155.000 | 0.9744 | -2204.000 | 0.0314 |
| TRIM59 | 1814.000 | 1515.000 | 0.6974 | -2.24 | 0.03144 |
| PSMA4 | 4831.000 | 4433.000 | 0.9454 | -2245.000 | 0.03145 |
| ABCD4 | 3453.000 | 3669.000 | 1049.000 | 2248.000 | 0.0315 |
| IQCD | 0.9342 | 1.18 | -2437.000 | 2258.000 | 0.03151 |
| MRPS35 | 5173.000 | 4744.000 | 0.9473 | -2252.000 | 0.03154 |
| CCDC71L | 2385.000 | 2032.000 | 0.8155 | -2239.000 | 0.03157 |
| ZNF783 | 2607.000 | 2822.000 | 1083.000 | 2197.000 | 0.0317 |
| SATB1 | 1975.000 | 1492.000 | 0.5883 | -2231.000 | 0.03179 |
| REEP2 | 0.9265 | 1496.000 | -5281.000 | 2261.000 | 0.03195 |
| B3GALT4 | 2461.000 | 2091.000 | 0.8194 | -2249.000 | 0.03196 |
| CD63 | 7548.000 | 7271.000 | 0.9815 | -2218.000 | 0.03204 |
| CDC16 | 4696.000 | 4421.000 | 0.9611 | -2238.000 | 0.03207 |
| TRAPPC1 | 5.76 | 5365.000 | 0.9594 | -2263.000 | 0.03211 |
| LRRC56 | 2204.000 | 2495.000 | 1157.000 | 2202.000 | 0.03213 |
| HSBP1 | 3664.000 | 3352.000 | 0.9315 | -2228.000 | 0.03215 |
| GABARAPL2 | 5863.000 | 5.44 | 0.9576 | -2239.000 | 0.03217 |

| | | | | | |
|---|---|---|---|---|---|
| NRIP1 | 3387.000 | 2936.000 | 0.8829 | -2256.000 | 0.0322 |
| MAIP1 | 3485.000 | 3106.000 | 0.9078 | -2.26 | 0.03226 |
| GTF2B | 4319.000 | 4076.000 | 0.9605 | -2219.000 | 0.03232 |
| NUP210L | 0.09725 | 0.1642 | 0.7753 | 2272.000 | 0.03235 |
| BID | 3182.000 | 2854.000 | 0.906 | -2217.000 | 0.03239 |
| KIAA1551 | 2946.000 | 2.64 | 0.8986 | -2.24 | 0.03244 |
| NCKIPSD | 3461.000 | 3825.000 | 1081.000 | 2254.000 | 0.03251 |
| CBX8 | 2572.000 | 2327.000 | 0.894 | -2.23 | 0.03258 |
| ZFAND2A | 3642.000 | 3389.000 | 0.9443 | -2189.000 | 0.03261 |
| CMIP | 2662.000 | 2351.000 | 0.8731 | -2223.000 | 0.03267 |
| DDX19A | 3384.000 | 3589.000 | 1048.000 | 2.23 | 0.03272 |
| CCDC183 | 3101.000 | 3592.000 | 1.13 | 2225.000 | 0.03273 |
| PARP8 | 2681.000 | 2313.000 | 0.8503 | -2221.000 | 0.03275 |
| GJC1 | 1.31 | 1678.000 | 1918.000 | 2.24 | 0.03285 |
| LMAN2 | 6533.000 | 6196.000 | 0.9718 | -2234.000 | 0.03286 |
| GRTP1 | 2921.000 | 2642.000 | 0.9062 | -2188.000 | 0.03289 |
| HIGD1A | 5664.000 | 6162.000 | 1049.000 | 2226.000 | 0.03292 |
| GTPBP4 | 3.46 | 3702.000 | 1054.000 | 2216.000 | 0.03293 |
| SOCS7 | 2549.000 | 2768.000 | 1088.000 | 2.21 | 0.03303 |
| EP400NL | 1322.000 | 1618.000 | 1721.000 | 2225.000 | 0.03305 |
| HYDIN | 0.1514 | 0.312 | 0.6169 | 2262.000 | 0.03306 |
| SPA17 | 1324.000 | 1053.000 | 0.1827 | -2211.000 | 0.03323 |
| ATP7A | 2.23 | 1946.000 | 0.83 | -2219.000 | 0.03337 |
| C14orf80 | 2407.000 | 2746.000 | 1.15 | 2218.000 | 0.03339 |
| DNAJB2 | 5072.000 | 4785.000 | 0.9642 | -2226.000 | 0.0335 |
| SMPD1 | 4264.000 | 3877.000 | 0.9344 | -2226.000 | 0.0336 |
| H2AFJ | 7.48 | 6853.000 | 0.9565 | -2206.000 | 0.03363 |
| PRG2 | 1033.000 | 0.51 | -21.01 | -2158.000 | 0.03365 |
| SLCO4C1 | 0.2399 | 0.09876 | 1622.000 | -2158.000 | 0.03368 |
| GGPS1 | 4006.000 | 3625.000 | 0.9281 | -2221.000 | 0.03379 |
| RUBCN | 2575.000 | 2777.000 | 1.08 | 2196.000 | 0.03387 |
| SRP72 | 4811.000 | 4497.000 | 0.9571 | -2224.000 | 0.03399 |
| GPS1 | 5724.000 | 5325.000 | 0.9586 | -2235.000 | 0.034 |
| MSMO1 | 4661.000 | 4154.000 | 0.9251 | -2211.000 | 0.03401 |
| SLC7A6 | 1516.000 | 1853.000 | 1484.000 | 2234.000 | 0.03404 |
| KLK12 | 1.21 | 2241.000 | 4242.000 | 2224.000 | 0.03405 |
| BCL2A1 | 1437.000 | 0.9994 | -0.00152 | -2169.000 | 0.03405 |
| C19orf57 | 0.9172 | 1364.000 | -3594.000 | 2.24 | 0.03438 |
| XPC | 3685.000 | 3956.000 | 1055.000 | 2.22 | 0.03444 |
| GSKIP | 2883.000 | 2606.000 | 0.9046 | -2161.000 | 0.03444 |
| C22orf23 | 1805.000 | 2092.000 | 1.25 | 2219.000 | 0.03453 |
| PPP4R3B | 4441.000 | 4181.000 | 0.9596 | -2221.000 | 0.03454 |
| HEBP2 | 3922.000 | 3537.000 | 0.9244 | -2195.000 | 0.03456 |
| DUOX1 | 0.8308 | 1177.000 | -0.8776 | 2211.000 | 0.03464 |
| ZSCAN20 | 0.5434 | 0.721 | 0.5364 | 2202.000 | 0.03471 |
| TTC39A | 2272.000 | 1821.000 | 0.7301 | -2195.000 | 0.03473 |

| | | | | | |
|---|---|---|---|---|---|
| UBE2K | 4.27 | 3928.000 | 0.9425 | -2234.000 | 0.03473 |
| AVL9 | 3526.000 | 3293.000 | 0.9458 | -2201.000 | 0.03474 |
| NDUFB4 | 6437.000 | 6022.000 | 0.9643 | -2217.000 | 0.03478 |
| PCCB | 3495.000 | 3188.000 | 0.9266 | -2189.000 | 0.03484 |
| IAH1 | 4109.000 | 3873.000 | 0.9582 | -2179.000 | 0.03501 |
| SZRD1 | 5333.000 | 5031.000 | 0.9651 | -2216.000 | 0.03506 |
| YIPF4 | 2738.000 | 2491.000 | 0.9059 | -2211.000 | 0.0351 |
| IRS2 | 3073.000 | 3757.000 | 1179.000 | 2176.000 | 0.03511 |
| GSDMB | 2577.000 | 3052.000 | 1179.000 | 2194.000 | 0.03516 |
| NGRN | 4155.000 | 3877.000 | 0.9514 | -2205.000 | 0.03519 |
| SPRY4 | 2795.000 | 3453.000 | 1206.000 | 2208.000 | 0.03528 |
| TMEM134 | 4019.000 | 3591.000 | 0.9191 | -2201.000 | 0.03528 |
| AMPD1 | 0.4144 | 0.1909 | 1.88 | -2156.000 | 0.03538 |
| TP53BP1 | 2542.000 | 2767.000 | 1091.000 | 2188.000 | 0.03542 |
| ASIP | 0.428 | 0.5773 | 0.6474 | 2193.000 | 0.03544 |
| FAM186B | 0.2943 | 0.4585 | 0.6376 | 2222.000 | 0.03548 |
| PEX2 | 3778.000 | 3.39 | 0.9186 | -2192.000 | 0.03548 |
| PHF19 | 2591.000 | 2919.000 | 1125.000 | 2169.000 | 0.03578 |
| TP53TG5 | 0.4099 | 0.5798 | 0.6112 | 2209.000 | 0.03582 |
| TDRD7 | 2545.000 | 2286.000 | 0.8847 | -2191.000 | 0.03587 |
| RTKN | 3423.000 | 3815.000 | 1088.000 | 2.18 | 0.03589 |
| BORCS7 | 3899.000 | 3552.000 | 0.9316 | -2187.000 | 0.036 |
| MRPL42 | 2536.000 | 2231.000 | 0.8625 | -2184.000 | 0.03606 |
| IRF2 | 4052.000 | 3775.000 | 0.9494 | -2194.000 | 0.0361 |
| LYG2 | 0.3295 | 0.4626 | 0.6943 | 2192.000 | 0.03616 |
| GLUL | 5895.000 | 5307.000 | 0.9408 | -2207.000 | 0.03622 |
| CCDC50 | 3.26 | 3015.000 | 0.9339 | -2164.000 | 0.03627 |
| SEL1L3 | 3662.000 | 3304.000 | 0.9208 | -2142.000 | 0.0363 |
| AGPAT2 | 4846.000 | 4462.000 | 0.9476 | -2178.000 | 0.03636 |
| SP110 | 3059.000 | 2719.000 | 0.8946 | -2167.000 | 0.03637 |
| MAP1LC3B | 4295.000 | 3875.000 | 0.9294 | -2.2 | 0.03642 |
| VBP1 | 4944.000 | 4477.000 | 0.9379 | -2189.000 | 0.03646 |
| EPDR1 | 3668.000 | 3121.000 | 0.8758 | -2183.000 | 0.03648 |
| UBE2L6 | 5252.000 | 4519.000 | 0.9093 | -2185.000 | 0.03669 |
| TNNI2 | 1113.000 | 0.6767 | -3645.000 | -2127.000 | 0.03673 |
| MAPK4 | 0.8355 | 1281.000 | -1376.000 | 2185.000 | 0.03676 |
| TDRD5 | 0.4812 | 0.2691 | 1794.000 | -2156.000 | 0.03677 |
| CCDC85B | 4019.000 | 3524.000 | 0.9056 | -2188.000 | 0.03678 |
| TMEM55A | 2296.000 | 2023.000 | 0.8475 | -2155.000 | 0.03681 |
| IRAK4 | 2537.000 | 2314.000 | 0.9013 | -2151.000 | 0.03691 |
| YWHAB | 6562.000 | 6216.000 | 0.9712 | -2.19 | 0.03693 |
| TXNDC12 | 4.57 | 4.24 | 0.9508 | -2176.000 | 0.03697 |
| TMSB10 | 10.02 | 9554.000 | 0.9792 | -2184.000 | 0.03698 |
| CPD | 3.96 | 3.61 | 0.9328 | -2177.000 | 0.03702 |
| EGFLAM | 1149.000 | 1827.000 | 4338.000 | 2211.000 | 0.03712 |
| MRPS22 | 4337.000 | 4069.000 | 0.9565 | -2155.000 | 0.03714 |

| | | | | | |
|---|---|---|---|---|---|
| OBSCN | 0.8382 | 1159.000 | -0.838 | 2181.000 | 0.03721 |
| THAP1 | 2709.000 | 2497.000 | 0.9184 | -2151.000 | 0.03722 |
| DECR1 | 5.23 | 4737.000 | 0.9401 | -2174.000 | 0.03727 |
| PLEKHG4 | 0.9491 | 1455.000 | -7174.000 | 2196.000 | 0.03735 |
| RBSN | 2734.000 | 2953.000 | 1077.000 | 2174.000 | 0.03738 |
| CCDC157 | 1209.000 | 1398.000 | 1767.000 | 2179.000 | 0.03741 |
| PLP1 | 0.2701 | 0.6933 | 0.2799 | 2213.000 | 0.03747 |
| CNPY2 | 6317.000 | 6048.000 | 0.9764 | -2167.000 | 0.03751 |
| REV1 | 3065.000 | 3248.000 | 1052.000 | 2156.000 | 0.03752 |
| C12orf29 | 3097.000 | 2823.000 | 0.9181 | -2147.000 | 0.03752 |
| NETO2 | 2021.000 | 2436.000 | 1266.000 | 2.15 | 0.03756 |
| FAM96A | 4675.000 | 4251.000 | 0.9383 | -2168.000 | 0.0376 |
| KAT7 | 2775.000 | 3047.000 | 1091.000 | 2185.000 | 0.03764 |
| TAS2R20 | 0.4199 | 0.5719 | 0.6439 | 2179.000 | 0.03769 |
| DUS2 | 2478.000 | 2701.000 | 1095.000 | 2173.000 | 0.03783 |
| ARMCX5 | 2346.000 | 2102.000 | 0.8713 | -2153.000 | 0.03791 |
| PABPC1L | 2421.000 | 2892.000 | 1201.000 | 2164.000 | 0.03792 |
| WASHC5 | 4152.000 | 3885.000 | 0.9532 | -2144.000 | 0.03792 |
| RAB27B | 2701.000 | 2219.000 | 0.802 | -2159.000 | 0.03799 |
| C8orf4 | 4259.000 | 3.58 | 0.8801 | -2165.000 | 0.03802 |
| GOLGA5 | 4644.000 | 4211.000 | 0.9364 | -2175.000 | 0.03803 |
| METTL25 | 1939.000 | 1648.000 | 0.7548 | -2166.000 | 0.03806 |
| HAUS4 | 4.04 | 3738.000 | 0.9444 | -2132.000 | 0.03808 |
| KMO | 0.4431 | 0.294 | 1504.000 | -2114.000 | 0.03813 |
| SYT2 | 0.2429 | 0.3921 | 0.6616 | 2183.000 | 0.03823 |
| RPA3 | 3.58 | 3193.000 | 0.9103 | -2159.000 | 0.03824 |
| FAM122A | 2858.000 | 2615.000 | 0.9154 | -2164.000 | 0.03824 |
| SURF1 | 5339.000 | 5082.000 | 0.9706 | -2152.000 | 0.03833 |
| TMEM170A | 2254.000 | 2085.000 | 0.9041 | -2119.000 | 0.03835 |
| DNPEP | 4703.000 | 4387.000 | 0.9551 | -2.18 | 0.03836 |
| ST3GAL3 | 2677.000 | 2893.000 | 1079.000 | 2135.000 | 0.0385 |
| IL15 | 0.4341 | 0.2886 | 1489.000 | -2107.000 | 0.03869 |
| CCDC110 | 1516.000 | 1245.000 | 0.5275 | -2.14 | 0.0387 |
| RINT1 | 3053.000 | 2839.000 | 0.9348 | -2138.000 | 0.03883 |
| FGGY | 2344.000 | 2092.000 | 0.8665 | -2.12 | 0.03888 |
| LRP6 | 2339.000 | 2635.000 | 1.14 | 2135.000 | 0.03891 |
| SERF2 | 7279.000 | 7.03 | 0.9825 | -2.13 | 0.03892 |
| YIF1A | 5661.000 | 5257.000 | 0.9572 | -2165.000 | 0.03892 |
| POLR2L | 6552.000 | 6081.000 | 0.9603 | -2168.000 | 0.03896 |
| HSPA6 | 1457.000 | 1087.000 | 0.2203 | -2107.000 | 0.0391 |
| FXYD3 | 4987.000 | 4359.000 | 0.9163 | -2135.000 | 0.03913 |
| KLHL4 | 0.2074 | 0.1466 | 1.22 | -2096.000 | 0.03919 |
| SHPK | 2521.000 | 2.77 | 1102.000 | 2.15 | 0.0392 |
| CD38 | 0.9725 | 0.6436 | 15.79 | -2.1 | 0.0394 |
| CHTF18 | 2896.000 | 3262.000 | 1112.000 | 2132.000 | 0.03942 |
| RALY | 4545.000 | 4255.000 | 0.9563 | -2157.000 | 0.03964 |

| | | | | | |
|---|---|---|---|---|---|
| UBXN11 | 2.97 | 3.28 | 1091.000 | 2142.000 | 0.03967 |
| CEP89 | 2.24 | 2473.000 | 1123.000 | 2141.000 | 0.03967 |
| CTDP1 | 2169.000 | 1971.000 | 0.8766 | -2145.000 | 0.03972 |
| SCAMP2 | 5244.000 | 4949.000 | 0.9651 | -2152.000 | 0.04002 |
| GNG3 | 0.749 | 1013.000 | -0.04609 | 2148.000 | 0.04008 |
| RBM28 | 1861.000 | 2.07 | 1171.000 | 2143.000 | 0.04018 |
| SNRPD3 | 5729.000 | 5459.000 | 0.9723 | -2126.000 | 0.04018 |
| WNT5B | 2499.000 | 1981.000 | 0.7464 | -2.13 | 0.0402 |
| SOSTDC1 | 0.4227 | 0.204 | 1846.000 | -2084.000 | 0.04023 |
| NIPSNAP3A | 4.6 | 4286.000 | 0.9537 | -2.12 | 0.04026 |
| LACTB | 2707.000 | 2429.000 | 0.8913 | -2136.000 | 0.04028 |
| RNF148 | 0.2314 | 0.3503 | 0.7168 | 2149.000 | 0.04029 |
| ZBTB38 | 3.11 | 2794.000 | 0.9056 | -2139.000 | 0.04031 |
| CLEC4D | 0.2555 | 0.1144 | 1589.000 | -2078.000 | 0.04039 |
| STARD10 | 4.93 | 4458.000 | 0.9369 | -2132.000 | 0.04048 |
| C3orf14 | 1278.000 | 0.7785 | -1021.000 | -2112.000 | 0.0405 |
| C2CD4D | 1168.000 | 0.9034 | -0.6545 | -2123.000 | 0.04058 |
| C1orf56 | 2761.000 | 2443.000 | 0.8797 | -2128.000 | 0.04064 |
| ATG4C | 2386.000 | 2102.000 | 0.8544 | -2147.000 | 0.04071 |
| PROSER3 | 1815.000 | 2077.000 | 1226.000 | 2135.000 | 0.04076 |
| MLH3 | 2087.000 | 2256.000 | 1106.000 | 2126.000 | 0.04077 |
| LHX4 | 0.8561 | 1063.000 | -0.396 | 2138.000 | 0.0408 |
| FRS3 | 2.1 | 2345.000 | 1148.000 | 2131.000 | 0.04091 |
| PALLD | 4044.000 | 3624.000 | 0.9215 | -2127.000 | 0.04092 |
| NAA30 | 2365.000 | 2191.000 | 0.911 | -2109.000 | 0.04094 |
| C11orf1 | 2376.000 | 2.07 | 0.8407 | -2122.000 | 0.04094 |
| RFK | 3577.000 | 3176.000 | 0.9066 | -2125.000 | 0.04094 |
| PELP1 | 3725.000 | 3988.000 | 1052.000 | 2127.000 | 0.04095 |
| EZH2 | 3.21 | 3677.000 | 1116.000 | 2136.000 | 0.04096 |
| CAND2 | 0.7185 | 1135.000 | -0.3833 | 2166.000 | 0.04098 |
| NUDT13 | 1501.000 | 1791.000 | 1436.000 | 2147.000 | 0.041 |
| ANKRD16 | 2173.000 | 2536.000 | 1199.000 | 2129.000 | 0.04101 |
| FUCA2 | 4822.000 | 4465.000 | 0.9511 | -2136.000 | 0.04107 |
| CECR2 | 0.773 | 0.483 | 2826.000 | -2112.000 | 0.04111 |
| MARK2 | 4343.000 | 4087.000 | 0.9587 | -2121.000 | 0.04111 |
| RPS16 | 9919.000 | 9561.000 | 0.984 | -2114.000 | 0.04116 |
| HLA-DMA | 5019.000 | 4522.000 | 0.9354 | -2107.000 | 0.04133 |
| SYCE2 | 1433.000 | 1664.000 | 1415.000 | 2122.000 | 0.0414 |
| PRDM6 | 0.2183 | 0.43 | 0.5546 | 2.16 | 0.04146 |
| HAX1 | 6181.000 | 5.93 | 0.9772 | -2096.000 | 0.04154 |
| AVPR1A | 0.7485 | 1074.000 | -0.2468 | 2121.000 | 0.04157 |
| THOC7 | 5603.000 | 5245.000 | 0.9616 | -2126.000 | 0.04161 |
| C14orf119 | 3634.000 | 3341.000 | 0.9348 | -2109.000 | 0.04169 |
| BRAT1 | 4548.000 | 4282.000 | 0.9602 | -2116.000 | 0.04179 |
| CLLU1OS | 0.1485 | 0.05804 | 1492.000 | -2064.000 | 0.0418 |
| MYDGF | 6226.000 | 5859.000 | 0.9668 | -2126.000 | 0.0418 |

| | | | | | |
|---|---|---|---|---|---|
| ANKRD53 | 0.2798 | 0.4382 | 0.6478 | 2147.000 | 0.04183 |
| TUBGCP4 | 1649.000 | 1848.000 | 1227.000 | 2123.000 | 0.04183 |
| PSMB1 | 7109.000 | 6729.000 | 0.9719 | -2122.000 | 0.04186 |
| CLCN6 | 2209.000 | 2498.000 | 1155.000 | 2129.000 | 0.04187 |
| METTL24 | 0.2893 | 0.4837 | 0.5856 | 2143.000 | 0.04192 |
| RNF125 | 1167.000 | 0.9489 | -0.3402 | -2104.000 | 0.04203 |
| SPN | 0.9227 | 0.6542 | 5277.000 | -2078.000 | 0.04209 |
| TOMM22 | 4972.000 | 4657.000 | 0.9592 | -2112.000 | 0.04211 |
| PRKG1 | 0.9156 | 1196.000 | -2027.000 | 2134.000 | 0.04212 |
| CRYGS | 1183.000 | 1532.000 | 2545.000 | 2134.000 | 0.04222 |
| HIRIP3 | 2746.000 | 3098.000 | 1.12 | 2126.000 | 0.04227 |
| HIBADH | 4915.000 | 4671.000 | 0.968 | -2099.000 | 0.0423 |
| OTUD3 | 1933.000 | 2194.000 | 1192.000 | 2126.000 | 0.04245 |
| PHF7 | 1888.000 | 2176.000 | 1223.000 | 2125.000 | 0.04247 |
| CAMKK1 | 1846.000 | 2225.000 | 1305.000 | 2121.000 | 0.0425 |
| MAP3K13 | 2543.000 | 2302.000 | 0.8935 | -2085.000 | 0.04253 |
| ADRA1D | 0.1276 | 0.3412 | 0.5223 | 2145.000 | 0.04258 |
| CFAP73 | 0.4489 | 0.8862 | 0.1508 | 2128.000 | 0.04263 |
| AGPAT5 | 3306.000 | 3577.000 | 1066.000 | 2084.000 | 0.0427 |
| BLOC1S2 | 3502.000 | 3233.000 | 0.9363 | -2096.000 | 0.0429 |
| LCN12 | 1511.000 | 1951.000 | 1.62 | 2108.000 | 0.04291 |
| TAPBP | 5725.000 | 5386.000 | 0.965 | -2108.000 | 0.04295 |
| THAP5 | 3411.000 | 3.07 | 0.914 | -2106.000 | 0.04297 |
| MED24 | 4124.000 | 4376.000 | 1042.000 | 2121.000 | 0.04299 |
| TTYH1 | 0.7014 | 0.8835 | 0.3491 | 2112.000 | 0.04299 |
| MIB2 | 3357.000 | 3.73 | 1087.000 | 2115.000 | 0.04313 |
| LYSMD2 | 2801.000 | 2456.000 | 0.8724 | -2101.000 | 0.04328 |
| CLDND1 | 4.32 | 4041.000 | 0.9543 | -2081.000 | 0.04332 |
| MZB1 | 3431.000 | 2876.000 | 0.8571 | -2.08 | 0.04333 |
| MEIS1 | 0.6357 | 1081.000 | -0.1715 | 2137.000 | 0.04352 |
| SULT1A1 | 3004.000 | 3417.000 | 1117.000 | 2.1 | 0.04352 |
| CCDC61 | 2377.000 | 2.65 | 1125.000 | 2099.000 | 0.04353 |
| C1orf115 | 3574.000 | 2967.000 | 0.854 | -2102.000 | 0.04357 |
| MINDY4 | 2371.000 | 2659.000 | 1132.000 | 2094.000 | 0.0436 |
| P4HB | 8944.000 | 8584.000 | 0.9812 | -2104.000 | 0.04372 |
| SPINK1 | 1454.000 | 0.8057 | -0.5776 | -2045.000 | 0.04377 |
| CASP2 | 3523.000 | 3849.000 | 1.07 | 2075.000 | 0.04386 |
| CDA | 2128.000 | 1657.000 | 0.6686 | -2071.000 | 0.04393 |
| TMSB4X | 8773.000 | 8315.000 | 0.9753 | -2096.000 | 0.04393 |
| SQSTM1 | 6054.000 | 5683.000 | 0.9648 | -2093.000 | 0.04394 |
| CCNJ | 1742.000 | 2009.000 | 1257.000 | 2094.000 | 0.04395 |
| ABCB8 | 3043.000 | 3274.000 | 1066.000 | 2074.000 | 0.04403 |
| ANKRD49 | 2644.000 | 2451.000 | 0.9221 | -2076.000 | 0.04404 |
| SWAP70 | 3399.000 | 3113.000 | 0.9281 | -2.09 | 0.04405 |
| ACSL5 | 3467.000 | 2.84 | 0.8396 | -2079.000 | 0.04407 |
| RELB | 2609.000 | 2299.000 | 0.868 | -2078.000 | 0.04413 |

| | | | | | |
|---|---|---|---|---|---|
| ZC3H18 | 3503.000 | 3281.000 | 0.948 | -2.08 | 0.04419 |
| HS2ST1 | 2971.000 | 2695.000 | 0.9105 | -2077.000 | 0.04424 |
| TAS2R4 | 0.4115 | 0.586 | 0.6018 | 2109.000 | 0.04425 |
| NR4A1 | 3368.000 | 4022.000 | 1146.000 | 2105.000 | 0.04428 |
| GATS | 1322.000 | 1658.000 | 1813.000 | 2.11 | 0.04429 |
| TAF4 | 2869.000 | 3117.000 | 1079.000 | 2093.000 | 0.04432 |
| ATP6V1A | 4607.000 | 4338.000 | 0.9606 | -2086.000 | 0.04442 |
| ARSA | 4362.000 | 4053.000 | 0.9501 | -2088.000 | 0.04442 |
| TCF25 | 4573.000 | 4302.000 | 0.9598 | -2099.000 | 0.04445 |
| PRDX6 | 7344.000 | 6.99 | 0.9752 | -2079.000 | 0.04446 |
| LAMTOR5 | 5591.000 | 5289.000 | 0.9677 | -2088.000 | 0.04449 |
| FAM102B | 2.39 | 2088.000 | 0.845 | -2097.000 | 0.04457 |
| UFL1 | 3676.000 | 3.37 | 0.9331 | -2092.000 | 0.04463 |
| PRMT3 | 3023.000 | 2779.000 | 0.9238 | -2085.000 | 0.04465 |
| HOXD3 | 0.6279 | 1065.000 | -0.1346 | 2117.000 | 0.04468 |
| C8orf59 | 5229.000 | 4.73 | 0.9393 | -2.07 | 0.04483 |
| N4BP1 | 3082.000 | 2857.000 | 0.9327 | -2072.000 | 0.04491 |
| CFAP36 | 4035.000 | 3676.000 | 0.9333 | -2092.000 | 0.04502 |
| RAB4B | 3681.000 | 3402.000 | 0.9395 | -2086.000 | 0.04503 |
| RBM17 | 4112.000 | 4345.000 | 1039.000 | 2084.000 | 0.04506 |
| PCMT1 | 5087.000 | 4747.000 | 0.9576 | -2093.000 | 0.04508 |
| CHST5 | 0.2254 | 0.3385 | 0.727 | 2099.000 | 0.04515 |
| WDR62 | 1685.000 | 2113.000 | 1433.000 | 2096.000 | 0.04519 |
| FBXO2 | 1679.000 | 2306.000 | 1613.000 | 2.09 | 0.04521 |
| NOL7 | 5399.000 | 5065.000 | 0.9621 | -2083.000 | 0.04533 |
| UBE2F | 4.1 | 3877.000 | 0.9605 | -2.08 | 0.04537 |
| MRPS33 | 3683.000 | 3.36 | 0.9297 | -2079.000 | 0.04538 |
| SLAMF7 | 1878.000 | 1444.000 | 0.5828 | -2055.000 | 0.04539 |
| SPINT2 | 7342.000 | 7.000 | 0.9761 | -2082.000 | 0.04539 |
| ACOT8 | 4524.000 | 4295.000 | 0.9656 | -2.08 | 0.04543 |
| SLC2A13 | 1058.000 | 0.8545 | -2768.000 | -2073.000 | 0.0455 |
| KCNQ4 | 0.7646 | 1106.000 | -0.376 | 2.1 | 0.04553 |
| PCDH19 | 0.3449 | 0.8137 | 0.1937 | 2109.000 | 0.04556 |
| MRPS17 | 3905.000 | 3593.000 | 0.9389 | -2087.000 | 0.04557 |
| ERP29 | 6114.000 | 5.67 | 0.9584 | -2.1 | 0.04559 |
| DDX58 | 2716.000 | 2.31 | 0.838 | -2073.000 | 0.04562 |
| ANKRD20A3 | 0.001646 | 0.007184 | 0.7701 | 2111.000 | 0.04575 |
| FABP9 | 0.32 | 0.1472 | 1682.000 | -2032.000 | 0.04584 |
| C12orf60 | 1.02 | 0.779 | -12.79 | -2049.000 | 0.04585 |
| MAPK15 | 1219.000 | 1792.000 | 2949.000 | 2086.000 | 0.04589 |
| GABRE | 0.799 | 1308.000 | -1197.000 | 2096.000 | 0.04591 |
| GSTO1 | 5058.000 | 4618.000 | 0.9438 | -2065.000 | 0.04591 |
| AGBL5 | 3867.000 | 3641.000 | 0.9555 | -2057.000 | 0.04601 |
| LIN52 | 2442.000 | 2695.000 | 1111.000 | 2072.000 | 0.04605 |
| MRPL38 | 5063.000 | 4751.000 | 0.9608 | -2082.000 | 0.04606 |
| VNN2 | 1286.000 | 0.829 | -0.7448 | -2049.000 | 0.04607 |

| | | | | | |
|---|---|---|---|---|---|
| TM7SF3 | 4469.000 | 4167.000 | 0.9534 | -2067.000 | 0.0461 |
| AK1 | 3.47 | 3153.000 | 0.923 | -2.07 | 0.04614 |
| CBARP | 0.4068 | 0.6592 | 0.4633 | 2096.000 | 0.04619 |
| UQCC3 | 3932.000 | 3507.000 | 0.9164 | -2084.000 | 0.04622 |
| TAF10 | 5352.000 | 4966.000 | 0.9554 | -2083.000 | 0.04623 |
| SMTN | 3468.000 | 3846.000 | 1083.000 | 2075.000 | 0.04624 |
| CACNA2D2 | 1815.000 | 2374.000 | 1451.000 | 2089.000 | 0.04626 |
| SLCO1B1 | 0.5704 | 0.1935 | 2925.000 | -2035.000 | 0.04627 |
| PDCD11 | 3211.000 | 3445.000 | 1.06 | 2068.000 | 0.04628 |
| RPS6KA6 | 1.63 | 1252.000 | 0.4601 | -2.08 | 0.04633 |
| ARMT1 | 3046.000 | 2753.000 | 0.9091 | -2078.000 | 0.04642 |
| COX6C | 6027.000 | 5646.000 | 0.9636 | -2076.000 | 0.04648 |
| UQCR10 | 6506.000 | 6168.000 | 0.9716 | -2068.000 | 0.04661 |
| CXCL8 | 1441.000 | 0.8536 | -0.4334 | -2036.000 | 0.04663 |
| DLL4 | 2402.000 | 2789.000 | 1.17 | 2.06 | 0.04669 |
| MMS19 | 3879.000 | 4129.000 | 1046.000 | 2067.000 | 0.04681 |
| FPGT | 2644.000 | 2393.000 | 0.8974 | -2066.000 | 0.04683 |
| RNFT2 | 0.827 | 1.17 | -0.8254 | 2084.000 | 0.0469 |
| NIPAL1 | 1276.000 | 1.02 | 0.07982 | -2051.000 | 0.04698 |
| KCNIP2 | 0.792 | 1124.000 | -0.5028 | 2086.000 | 0.04699 |
| LACTB2 | 3995.000 | 3612.000 | 0.9272 | -2062.000 | 0.04706 |
| PPP1R15B | 4394.000 | 4.12 | 0.9566 | -2.07 | 0.04713 |
| UGT1A3 | 0.465 | 0.2173 | 1994.000 | -2.02 | 0.04714 |
| ACYP1 | 2623.000 | 2967.000 | 1128.000 | 2.07 | 0.04715 |
| LRP8 | 1194.000 | 1528.000 | 2391.000 | 2069.000 | 0.04723 |
| ZYG11B | 2625.000 | 2426.000 | 0.9182 | -2069.000 | 0.04723 |
| TYW3 | 2977.000 | 2692.000 | 0.9079 | -2069.000 | 0.04726 |
| ATP6V1F | 6851.000 | 6476.000 | 0.9708 | -2077.000 | 0.04727 |
| EFNA2 | 0.9097 | 0.5978 | 5435.000 | -2032.000 | 0.04728 |
| DNAJA1 | 6743.000 | 6.48 | 0.9792 | -2057.000 | 0.04737 |
| ZDHHC11 | 1235.000 | 1614.000 | 2272.000 | 2072.000 | 0.04741 |
| GRIN2A | 0.06646 | 0.1888 | 0.6149 | 2093.000 | 0.04755 |
| KYNU | 0.2708 | 0.1758 | 1331.000 | -2006.000 | 0.04756 |
| TSTD1 | 6404.000 | 5926.000 | 0.9582 | -2061.000 | 0.0476 |
| MLLT6 | 4386.000 | 4.68 | 1044.000 | 2047.000 | 0.04769 |
| MIEN1 | 5742.000 | 5391.000 | 0.9639 | -2062.000 | 0.04777 |
| PHKB | 4283.000 | 3965.000 | 0.947 | -2048.000 | 0.04779 |
| IQCG | 2382.000 | 2693.000 | 1141.000 | 2056.000 | 0.0479 |
| FAAH2 | 2732.000 | 2.42 | 0.8793 | -2048.000 | 0.04792 |
| RIDA | 4403.000 | 4039.000 | 0.9418 | -2029.000 | 0.04794 |
| PRAF2 | 2812.000 | 3117.000 | 1.1 | 2041.000 | 0.04803 |
| RNF139 | 4346.000 | 4011.000 | 0.9453 | -2053.000 | 0.04809 |
| DLG4 | 1804.000 | 2166.000 | 1309.000 | 2.05 | 0.04811 |
| AC009477.8 | 0.1076 | 0.04093 | 1434.000 | -2.01 | 0.0482 |
| SLC26A4 | 0.7205 | 0.4366 | 2528.000 | -2015.000 | 0.04821 |
| CEBPZOS | 3.84 | 3.56 | 0.9437 | -2053.000 | 0.04824 |

| | | | | | |
|---|---|---|---|---|---|
| MAL2 | 5523.000 | 4935.000 | 0.9342 | -2063.000 | 0.04828 |
| CST5 | 0.1029 | 0.7648 | 0.1179 | 2093.000 | 0.04829 |
| METTL9 | 5144.000 | 4791.000 | 0.9566 | -2047.000 | 0.04839 |
| SLX4 | 1.74 | 1925.000 | 1183.000 | 2054.000 | 0.04848 |
| TIAF1 | 0.4982 | 0.8345 | 0.2597 | 2083.000 | 0.04852 |
| PDZD2 | 1192.000 | 1559.000 | 2522.000 | 2061.000 | 0.04855 |
| AP4B1 | 2931.000 | 2757.000 | 0.9433 | -2031.000 | 0.04856 |
| HEATR4 | 0.3715 | 0.5132 | 0.6738 | 2057.000 | 0.04862 |
| UPF3B | 3169.000 | 3475.000 | 1.08 | 2055.000 | 0.04869 |
| OS9 | 6809.000 | 6.51 | 0.9766 | -2058.000 | 0.04879 |
| EDA2R | 0.8755 | 1376.000 | -2.4 | 2066.000 | 0.04881 |
| SDSL | 3642.000 | 3216.000 | 0.9038 | -2065.000 | 0.04886 |
| CADM3 | 0.3421 | 0.6838 | 0.3544 | 2065.000 | 0.0489 |
| MSANTD1 | 0.1487 | 0.2119 | 0.814 | 2054.000 | 0.04896 |
| PSMB8 | 4909.000 | 4372.000 | 0.9272 | -2.04 | 0.04898 |
| INTS9 | 2628.000 | 2849.000 | 1084.000 | 2038.000 | 0.0491 |
| NPR1 | 1363.000 | 1748.000 | 1806.000 | 2053.000 | 0.04923 |
| SEMA5A | 1606.000 | 1267.000 | 0.4988 | -2028.000 | 0.04924 |
| UVSSA | 2195.000 | 2574.000 | 1202.000 | 2051.000 | 0.04926 |
| CNTROB | 3178.000 | 3382.000 | 1054.000 | 2036.000 | 0.04928 |
| HIST1H2BI | 0.237 | 0.1472 | 1331.000 | -1.99 | 0.04928 |
| SUPT4H1 | 5107.000 | 4838.000 | 0.9668 | -2036.000 | 0.04937 |
| ZBTB34 | 1964.000 | 1789.000 | 0.8616 | -2043.000 | 0.04941 |
| NRN1L | 0.7735 | 1042.000 | -0.1602 | 2057.000 | 0.04945 |
| AGAP1 | 2227.000 | 2506.000 | 1147.000 | 2049.000 | 0.04946 |
| PHF11 | 2915.000 | 2513.000 | 0.8612 | -2056.000 | 0.04947 |
| MRPL19 | 3533.000 | 3176.000 | 0.9156 | -2.06 | 0.0495 |
| TAC4 | 0.742 | 0.9925 | 0.02529 | 2048.000 | 0.04954 |
| DCAF4 | 2079.000 | 2273.000 | 1122.000 | 2.04 | 0.04956 |
| KTI12 | 2472.000 | 2231.000 | 0.8866 | -2045.000 | 0.0497 |
| SLC35D1 | 2302.000 | 2061.000 | 0.8676 | -2034.000 | 0.0498 |
| ISM1 | 0.943 | 1497.000 | -6875.000 | 2063.000 | 0.04989 |
| NUDT1 | 3.79 | 3388.000 | 0.9158 | -2045.000 | 0.0499 |

**Supplementary table 2. Pathway enrichment analysis on the Hallmarks pathways of the MSigDB**

| PATHWAY NAME | SIZE (genes) | ES | NES | NOM p-val | FDR q-val | FWER p-val | RANK AT MAX |
|---|---|---|---|---|---|---|---|
| KRAS_SIGNALING_DN | 198 | 0.2922989 | 1.6091298 | 0 | 0.04009493 | 0.058 | 3363 |
| SPERMATOGENESIS | 133 | 0.2656372 | 1.385252 | 0.01677852 | 0.12491626 | 0.318 | 7489 |
| E2F_TARGETS | 199 | 0.2282677 | 1.2353984 | 0.07392996 | 0.29266143 | 0.727 | 4248 |
| G2M_CHECKPOINT | 195 | 0.2093369 | 1.1450602 | 0.13333334 | 0.44077528 | 0.928 | 5464 |
| HEDGEHOG_SIGNALING | 36 | 0.2919669 | 1.1416843 | 0.24808185 | 0.36112 | 0.933 | 5911 |
| APICAL_SURFACE | 43 | 0.2569605 | 1.0485766 | 0.34375 | 0.5620716 | 0.996 | 4615 |
| WNT_BETA_CATENIN_SIGNALING | 42 | 0.2487561 | 1.0190428 | 0.42894056 | 0.5760681 | 0.999 | 4073 |
| MYC_TARGETS_V2 | 58 | 0.2150904 | 0.9452585 | 0.5605263 | 0.7445407 | 1 | 3141 |
| MYOGENESIS | 200 | 0.1582564 | 0.8576837 | 0.8745247 | 0.90126055 | 1 | 3462 |
| MITOTIC_SPINDLE | 197 | 0.1327497 | 0.7275849 | 0.9920949 | 0.9686404 | 1 | 1808 |
| INTERFERON_ALPHA_RESPONSE | 95 | -0.6776878 | -2.9997575 | 0 | 0 | 0 | 4023 |
| INTERFERON_GAMMA_RESPONSE | 198 | -0.5985509 | -2.9901364 | 0 | 0 | 0 | 4045 |
| OXIDATIVE_PHOSPHORYLATION | 185 | -0.5912627 | -2.8904471 | 0 | 0 | 0 | 3710 |
| PROTEIN_SECRETION | 95 | -0.603103 | -2.6498737 | 0 | 0 | 0 | 3166 |
| HEME_METABOLISM | 193 | -0.5318503 | -2.6248302 | 0 | 0 | 0 | 4556 |
| ALLOGRAFT_REJECTION | 199 | -0.5228689 | -2.5535924 | 0 | 0 | 0 | 4207 |
| MTORC1_SIGNALING | 199 | -0.5105973 | -2.5160353 | 0 | 0 | 0 | 5381 |
| REACTIVE_OXYGEN_SPECIES_PATHWAY | 49 | -0.6131204 | -2.3946645 | 0 | 0 | 0 | 4256 |
| IL6_JAK_STAT3_SIGNALING | 86 | -0.5443632 | -2.349432 | 0 | 0 | 0 | 3603 |
| FATTY_ACID_METABOLISM | 157 | -0.4656699 | -2.2135122 | 0 | 0 | 0 | 4032 |
| PI3K_AKT_MTOR_SIGNALING | 105 | -0.4686388 | -2.1050143 | 0 | 0 | 0 | 4531 |
| COMPLEMENT | 199 | -0.4270765 | -2.092526 | 0 | 0 | 0 | 4598 |
| PEROXISOME | 104 | -0.4546987 | -2.04527 | 0 | 0 | 0 | 3003 |
| ANDROGEN_RESPONSE | 98 | -0.4548532 | -2.0024736 | 0 | 8.16E-05 | 0.001 | 4459 |
| ADIPOGENESIS | 195 | -0.4048453 | -1.9884899 | 0 | 7.62E-05 | 0.001 | 3448 |
| INFLAMMATORY_RESPONSE | 200 | -0.3987486 | -1.933964 | 0 | 2.05E-04 | 0.003 | 5474 |
| GLYCOLYSIS | 198 | -0.3890247 | -1.9124 | 0 | 3.37E-04 | 0.005 | 4818 |

| | | | | | | | |
|---|---|---|---|---|---|---|---|
| APOPTOSIS | 161 | -0.3860727 | -1.8172923 | 0 | 5.76E-04 | 0.009 | 4059 |
| XENOBIOTIC_METABOLISM | 198 | -0.3659605 | -1.7959694 | 0 | 7.06E-04 | 0.012 | 4010 |
| DNA_REPAIR | 148 | -0.3624897 | -1.7274706 | 0 | 0.00140326 | 0.024 | 4291 |
| BILE_ACID_METABOLISM | 112 | -0.3719493 | -1.688499 | 0 | 0.00236115 | 0.041 | 3666 |
| TNFA_SIGNALING_VIA_NFKB | 198 | -0.3398308 | -1.672165 | 0 | 0.00309908 | 0.057 | 4059 |
| KRAS_SIGNALING_UP | 199 | -0.3406309 | -1.6704649 | 0 | 0.00301134 | 0.058 | 4932 |
| CHOLESTEROL_HOMEOSTASIS | 74 | -0.3942043 | -1.6668487 | 0.00154799 | 0.00297875 | 0.06 | 4644 |
| IL2_STAT5_SIGNALING | 199 | -0.3373771 | -1.6494422 | 0.00138696 | 0.0032963 | 0.07 | 5115 |
| P53_PATHWAY | 198 | -0.335552 | -1.6375108 | 0.0013459 | 0.00346592 | 0.077 | 4790 |
| UV_RESPONSE_UP | 158 | -0.3381449 | -1.6012648 | 0.00137174 | 0.00469433 | 0.108 | 3775 |
| MYC_TARGETS_V1 | 199 | -0.3246678 | -1.5981044 | 0.0013369 | 0.00461461 | 0.109 | 6088 |
| UNFOLDED_PROTEIN_RESPONSE | 111 | -0.3319021 | -1.4925838 | 0.01144492 | 0.01315236 | 0.284 | 3712 |
| COAGULATION | 138 | -0.2957965 | -1.3890865 | 0.03623188 | 0.03795161 | 0.646 | 6476 |
| TGF_BETA_SIGNALING | 54 | -0.3163944 | -1.2717184 | 0.12422361 | 0.10680526 | 0.949 | 5047 |
| NOTCH_SIGNALING | 32 | -0.3424361 | -1.2276186 | 0.19769357 | 0.14568757 | 0.992 | 1363 |
| PANCREAS_BETA_CELLS | 40 | -0.2964255 | -1.0991397 | 0.32961783 | 0.33191034 | 1 | 483 |
| APICAL_JUNCTION | 199 | -0.2170201 | -1.0631317 | 0.32666665 | 0.39695343 | 1 | 4038 |
| HYPOXIA | 195 | -0.2195579 | -1.0614858 | 0.33241758 | 0.38879797 | 1 | 4879 |
| ESTROGEN_RESPONSE_EARLY | 197 | -0.2152124 | -1.0610038 | 0.352459 | 0.3787836 | 1 | 4870 |
| ESTROGEN_RESPONSE_LATE | 198 | -0.2126083 | -1.0436997 | 0.36649215 | 0.40242147 | 1 | 5313 |
| EPITHELIAL_MESENCHYMAL_TRANSITION | 198 | -0.1829763 | -0.8977385 | 0.6986667 | 0.7183605 | 1 | 5537 |
| UV_RESPONSE_DN | 140 | -0.1294529 | -0.608626 | 0.9986111 | 1 | 1 | 4305 |
| ANGIOGENESIS | 36 | -0.1494 | -0.5394515 | 0.99312717 | 0.9986594 | 1 | 2825 |

| Supplementary table 3. CXCR2 and BCL-2 correlation to Hallmark pathway enrichment | | | | | | | | | | |
|---|---|---|---|---|---|---|---|---|---|---|
| | | | | | Gene associated values | | eatment associated valu | | e-treatment interaction val | |
| **Pathway** | **gene** | **rho** | **pvalue** | **fdr** | **T-stats** | **Pvalue** | **T-stats** | **Pvalue** | **T-stats** | **Pvalue** |
| TNFA_SIGNALING_VIA_NFKB | CXCR2 | 0.3496 | | 0.0033 | 2.4125 | 0.0177 | 0.1574 | 0.8753 | 1.0477 | 0.2973 |
| HYPOXIA | CXCR2 | 0.1151 | 0.2441 | 0.3344 | -0.2477 | 0.8048 | 0.4994 | 0.6186 | 1.9796 | 0.0505 |
| CHOLESTEROL_HOMEOSTASIS | CXCR2 | -0.1417 | 0.1510 | 0.2288 | -1.3823 | 0.1700 | -1.0139 | 0.3131 | 1.7586 | 0.0817 |
| MITOTIC_SPINDLE | CXCR2 | -0.1519 | 0.1235 | 0.1960 | -1.6708 | 0.0979 | 0.9741 | 0.3323 | 0.7294 | 0.4675 |
| WNT_BETA_CATENIN_SIGNALING | CXCR2 | -0.0160 | 0.8716 | 0.8804 | -1.0222 | 0.3092 | 0.8077 | 0.4213 | 1.7903 | 0.0765 |
| TGF_BETA_SIGNALING | CXCR2 | 0.0947 | 0.3383 | 0.4282 | -0.8534 | 0.3955 | -0.0600 | 0.9523 | 1.9271 | 0.0568 |
| IL6_JAK_STAT3_SIGNALING | CXCR2 | 0.4987 | 0.0000 | 0.0000 | 4.0399 | 0.0001 | -1.2388 | 0.2183 | 1.0316 | 0.3047 |
| DNA_REPAIR | CXCR2 | -0.1248 | 0.2063 | 0.2948 | 0.6225 | 0.5350 | -0.4650 | 0.6429 | -1.3140 | 0.1918 |
| G2M_CHECKPOINT | CXCR2 | -0.1994 | 0.0425 | 0.0861 | -1.2713 | 0.2066 | 0.7741 | 0.4407 | 0.3273 | 0.7441 |
| APOPTOSIS | CXCR2 | 0.3498 | 0.0003 | 0.0033 | 1.5151 | 0.1329 | -0.6847 | 0.4951 | 2.0277 | 0.0453 |
| NOTCH_SIGNALING | CXCR2 | -0.0166 | 0.8672 | 0.8804 | -1.8715 | 0.0642 | 0.2504 | 0.8028 | 1.9085 | 0.0592 |
| ADIPOGENESIS | CXCR2 | -0.0602 | 0.5433 | 0.6104 | -1.2880 | 0.2008 | -1.5080 | 0.1347 | 0.3684 | 0.7134 |
| ESTROGEN_RESPONSE_EARLY | CXCR2 | -0.1082 | 0.2738 | 0.3700 | -1.8822 | 0.0627 | -0.0643 | 0.9488 | 1.4340 | 0.1547 |
| ESTROGEN_RESPONSE_LATE | CXCR2 | 0.1042 | 0.2919 | 0.3827 | -0.4280 | 0.6696 | 0.0278 | 0.9779 | 1.9433 | 0.0548 |
| ANDROGEN_RESPONSE | CXCR2 | -0.2338 | 0.0171 | 0.0462 | -1.6703 | 0.0980 | -1.0778 | 0.2837 | -0.1356 | 0.8924 |
| MYOGENESIS | CXCR2 | 0.2501 | 0.0106 | 0.0376 | -1.0445 | 0.2988 | 0.4039 | 0.6872 | 1.9880 | 0.0496 |
| PROTEIN_SECRETION | CXCR2 | -0.1508 | 0.1264 | 0.1975 | -0.7869 | 0.4332 | -2.1924 | 0.0307 | -0.4679 | 0.6409 |
| INTERFERON_ALPHA_RESPONSE | CXCR2 | 0.2480 | 0.0113 | 0.0377 | 2.0138 | 0.0467 | -1.8179 | 0.0721 | 0.3521 | 0.7255 |
| INTERFERON_GAMMA_RESPONSE | CXCR2 | 0.2916 | 0.0028 | 0.0145 | 2.1335 | 0.0354 | -1.0299 | 0.3056 | 0.5291 | 0.5979 |
| APICAL_JUNCTION | CXCR2 | 0.2509 | 0.0104 | 0.0376 | -0.6007 | 0.5494 | 0.0272 | 0.9784 | 2.4813 | 0.0148 |
| APICAL_SURFACE | CXCR2 | 0.3076 | 0.0016 | 0.0092 | 0.9165 | 0.3616 | 0.7812 | 0.4365 | 2.3103 | 0.0229 |
| HEDGEHOG_SIGNALING | CXCR2 | -0.1666 | 0.0910 | 0.1492 | -2.1098 | 0.0374 | 0.4157 | 0.6785 | 1.0180 | 0.3112 |
| COMPLEMENT | CXCR2 | 0.4001 | 0.0000 | 0.0005 | 2.8179 | 0.0058 | -0.9594 | 0.3397 | 0.8579 | 0.3930 |
| UNFOLDED_PROTEIN_RESPONSE | CXCR2 | -0.2137 | 0.0296 | 0.0674 | -0.2512 | 0.8022 | 0.3174 | 0.7516 | 0.1571 | 0.8755 |
| PI3K_AKT_MTOR_SIGNALING | CXCR2 | 0.0201 | 0.8395 | 0.8745 | 2.0609 | 0.0419 | -2.4007 | 0.0182 | -0.2565 | 0.7981 |
| MTORC1_SIGNALING | CXCR2 | -0.1207 | 0.2221 | 0.3084 | -0.2653 | 0.7913 | -1.2927 | 0.1991 | 0.7867 | 0.4333 |

| | | | | | | | | | | |
|---|---|---|---|---|---|---|---|---|---|---|
| E2F_TARGETS | CXCR2 | -0.1786 | 0.0698 | 0.1224 | -1.3024 | 0.1958 | 0.9852 | 0.3269 | 0.3858 | 0.7004 |
| MYC_TARGETS_V1 | CXCR2 | -0.2492 | 0.0109 | 0.0376 | -1.3439 | 0.1820 | 0.8188 | 0.4149 | -0.0936 | 0.9256 |
| MYC_TARGETS_V2 | CXCR2 | -0.2373 | 0.0155 | 0.0456 | -1.0782 | 0.2835 | 0.6370 | 0.5255 | -0.5387 | 0.5913 |
| EPITHELIAL_MESENCHYMAL_TRANSITION | CXCR2 | 0.1986 | 0.0434 | 0.0861 | -1.0081 | 0.3159 | 0.2778 | 0.7818 | 2.3611 | 0.0202 |
| INFLAMMATORY_RESPONSE | CXCR2 | 0.4572 | 0.0000 | 0.0000 | 3.7491 | 0.0003 | -1.0025 | 0.3186 | 0.6573 | 0.5125 |
| XENOBIOTIC_METABOLISM | CXCR2 | 0.2342 | 0.0169 | 0.0462 | 0.5835 | 0.5608 | -0.5633 | 0.5745 | 0.4696 | 0.6396 |
| FATTY_ACID_METABOLISM | CXCR2 | -0.2136 | 0.0297 | 0.0674 | -1.1179 | 0.2663 | -1.5068 | 0.1351 | -0.3146 | 0.7537 |
| OXIDATIVE_PHOSPHORYLATION | CXCR2 | -0.2095 | 0.0330 | 0.0702 | -0.7776 | 0.4386 | -1.9434 | 0.0548 | -0.1524 | 0.8792 |
| GLYCOLYSIS | CXCR2 | -0.1755 | 0.0749 | 0.1269 | -2.5544 | 0.0122 | -1.2699 | 0.2071 | 1.9080 | 0.0593 |
| REACTIVE_OXYGEN_SPECIES_PATHWAY | CXCR2 | 0.3193 | 0.0010 | 0.0073 | 4.1614 | 0.0001 | -2.2139 | 0.0291 | -0.1606 | 0.8728 |
| P53_PATHWAY | CXCR2 | 0.4053 | 0.0000 | 0.0005 | 2.3027 | 0.0234 | -0.1580 | 0.8747 | 1.4694 | 0.1449 |
| UV_RESPONSE_UP | CXCR2 | 0.1491 | 0.1307 | 0.2011 | 1.0625 | 0.2906 | 0.5653 | 0.5731 | 1.6881 | 0.0946 |
| UV_RESPONSE_DN | CXCR2 | 0.0317 | 0.7489 | 0.8159 | -1.5366 | 0.1276 | -0.3231 | 0.7473 | 0.7833 | 0.4354 |
| ANGIOGENESIS | CXCR2 | 0.3164 | 0.0011 | 0.0075 | 1.2544 | 0.2126 | 0.3563 | 0.7224 | 1.4054 | 0.1630 |
| HEME_METABOLISM | CXCR2 | 0.4486 | 0.0000 | 0.0001 | 5.4606 | 0.0000 | -2.7810 | 0.0065 | 1.3479 | 0.1808 |
| COAGULATION | CXCR2 | 0.2190 | 0.0257 | 0.0627 | -0.4336 | 0.6655 | -0.1214 | 0.9036 | 1.3942 | 0.1664 |
| IL2_STAT5_SIGNALING | CXCR2 | 0.2827 | 0.0038 | 0.0163 | 1.5054 | 0.1354 | 0.0471 | 0.9625 | 1.7832 | 0.0776 |
| BILE_ACID_METABOLISM | CXCR2 | -0.2267 | 0.0208 | 0.0549 | -2.0367 | 0.0444 | -0.8826 | 0.3796 | -0.4571 | 0.6486 |
| PEROXISOME | CXCR2 | -0.2155 | 0.0282 | 0.0672 | -2.0081 | 0.0473 | -0.8931 | 0.3740 | -0.1034 | 0.9179 |
| ALLOGRAFT_REJECTION | CXCR2 | 0.4856 | 0.0000 | 0.0000 | 3.8843 | 0.0002 | -1.0731 | 0.2858 | 0.6982 | 0.4867 |
| SPERMATOGENESIS | CXCR2 | -0.2864 | 0.0033 | 0.0158 | -1.6460 | 0.1029 | 1.3467 | 0.1811 | -0.2061 | 0.8371 |
| KRAS_SIGNALING_UP | CXCR2 | 0.3583 | 0.0002 | 0.0030 | 1.7449 | 0.0841 | -0.8389 | 0.4035 | 1.1957 | 0.2346 |
| KRAS_SIGNALING_DN | CXCR2 | 0.0391 | 0.6931 | 0.7702 | -0.2590 | 0.7961 | 1.0734 | 0.2857 | 0.6642 | 0.5081 |
| PANCREAS_BETA_CELLS | CXCR2 | -0.0829 | 0.4023 | 0.4789 | -2.2694 | 0.0254 | -1.3284 | 0.1871 | 1.3871 | 0.1685 |
| TNFA_SIGNALING_VIA_NFKB | BCL2 | 0.1072 | 0.2783 | 0.3710 | 0.6213 | 0.5358 | -0.1798 | 0.8577 | 0.3703 | 0.7119 |
| HYPOXIA | BCL2 | 0.0851 | 0.3896 | 0.4722 | 0.4866 | 0.6276 | 0.2357 | 0.8142 | 1.3902 | 0.1676 |
| CHOLESTEROL_HOMEOSTASIS | BCL2 | -0.2085 | 0.0338 | 0.0705 | -1.6773 | 0.0966 | -1.0561 | 0.2935 | -0.2809 | 0.7794 |
| MITOTIC_SPINDLE | BCL2 | 0.1872 | 0.0572 | 0.1040 | 3.2489 | 0.0016 | 0.7327 | 0.4655 | -2.3191 | 0.0224 |
| WNT_BETA_CATENIN_SIGNALING | BCL2 | 0.1965 | 0.0458 | 0.0880 | 1.2854 | 0.2016 | 0.7264 | 0.4693 | 0.9389 | 0.3501 |

| | | | | | | | | | | |
|---|---|---|---|---|---|---|---|---|---|---|
| TGF_BETA_SIGNALING | BCL2 | 0.1982 | 0.0439 | 0.0861 | 0.7273 | 0.4687 | -0.4548 | 0.6502 | 1.0250 | 0.3078 |
| IL6_JAK_STAT3_SIGNALING | BCL2 | 0.2668 | 0.0063 | 0.0244 | 2.1085 | 0.0375 | -0.6029 | 0.5480 | -0.5441 | 0.5876 |
| DNA_REPAIR | BCL2 | 0.0799 | 0.4192 | 0.4932 | 0.4632 | 0.6442 | 0.1103 | 0.9124 | -1.2659 | 0.2085 |
| G2M_CHECKPOINT | BCL2 | 0.0887 | 0.3699 | 0.4567 | 1.4236 | 0.1577 | 0.8074 | 0.4213 | -1.2386 | 0.2184 |
| APOPTOSIS | BCL2 | 0.2466 | 0.0118 | 0.0381 | 1.1704 | 0.2447 | -1.0588 | 0.2923 | 1.3715 | 0.0302 |
| NOTCH_SIGNALING | BCL2 | -0.0315 | 0.7507 | 0.8159 | 0.1517 | 0.8797 | 0.1726 | 0.8633 | -0.3276 | 0.7439 |
| ADIPOGENESIS | BCL2 | -0.0761 | 0.4418 | 0.5078 | -0.8592 | 0.3923 | -1.5106 | 0.1341 | -0.8917 | 0.3747 |
| ESTROGEN_RESPONSE_EARLY | BCL2 | -0.0847 | 0.3920 | 0.4722 | -1.2302 | 0.2215 | -0.1631 | 0.8708 | 1.5407 | 0.1266 |
| ESTROGEN_RESPONSE_LATE | BCL2 | 0.0299 | 0.7631 | 0.8206 | -0.0604 | 0.9520 | 0.1324 | 0.8949 | 1.4307 | 0.1556 |
| ANDROGEN_RESPONSE | BCL2 | -0.2340 | 0.0170 | 0.0462 | -1.8564 | 0.0663 | -0.8643 | 0.3895 | -1.8678 | 0.0647 |
| MYOGENESIS | BCL2 | 0.3332 | 0.0006 | 0.0058 | 2.1900 | 0.0309 | -0.0035 | 0.9972 | 0.4365 | 0.6634 |
| PROTEIN_SECRETION | BCL2 | -0.3234 | 0.0009 | 0.0072 | -3.1609 | 0.0021 | -2.8818 | 0.0049 | -1.2577 | 0.2115 |
| INTERFERON_ALPHA_RESPONSE | BCL2 | 0.0787 | 0.4268 | 0.4963 | -0.3354 | 0.7380 | -2.0153 | 0.0466 | 0.9603 | 0.3392 |
| INTERFERON_GAMMA_RESPONSE | BCL2 | 0.1537 | 0.1192 | 0.1923 | 0.6084 | 0.5443 | -1.1916 | 0.2363 | 0.4247 | 0.6720 |
| APICAL_JUNCTION | BCL2 | 0.3188 | 0.0010 | 0.0073 | 1.6329 | 0.1056 | -0.0165 | 0.9869 | 0.7245 | 0.4705 |
| APICAL_SURFACE | BCL2 | 0.1839 | 0.0617 | 0.1102 | 0.1524 | 0.8792 | 0.3257 | 0.7453 | 2.2386 | 0.0274 |
| HEDGEHOG_SIGNALING | BCL2 | 0.1918 | 0.0512 | 0.0949 | 1.8320 | 0.0699 | 0.8902 | 0.3755 | 1.5365 | 0.1276 |
| COMPLEMENT | BCL2 | 0.2790 | 0.0043 | 0.0177 | 2.3105 | 0.0229 | -1.1989 | 0.2334 | 0.8078 | 0.4211 |
| UNFOLDED_PROTEIN_RESPONSE | BCL2 | -0.2233 | 0.0229 | 0.0586 | -1.7103 | 0.0903 | -0.0127 | 0.9899 | -0.7603 | 0.4489 |
| PI3K_AKT_MTOR_SIGNALING | BCL2 | -0.1711 | 0.0825 | 0.1376 | -0.5758 | 0.5660 | -2.3208 | 0.0223 | -1.0992 | 0.2744 |
| MTORC1_SIGNALING | BCL2 | -0.0732 | 0.4599 | 0.5226 | -0.4060 | 0.6856 | -1.1235 | 0.2639 | -0.3665 | 0.7148 |
| E2F_TARGETS | BCL2 | 0.1021 | 0.3017 | 0.3868 | 1.4247 | 0.1574 | 0.7439 | 0.4587 | -1.0363 | 0.3026 |
| MYC_TARGETS_V1 | BCL2 | 0.0138 | 0.8891 | 0.8891 | 0.5051 | 0.6146 | 0.3744 | 0.7089 | -0.9548 | 0.3420 |
| MYC_TARGETS_V2 | BCL2 | -0.1036 | 0.2947 | 0.3827 | -0.7583 | 0.4501 | 0.8447 | 0.4003 | -1.3488 | 0.1805 |
| EPITHELIAL_MESENCHYMAL_TRANSITION | BCL2 | 0.3134 | 0.0013 | 0.0079 | 2.0297 | 0.0450 | -0.2001 | 0.8418 | 0.6037 | 0.5474 |
| INFLAMMATORY_RESPONSE | BCL2 | 0.2212 | 0.0242 | 0.0605 | 1.5199 | 0.1317 | -1.1816 | 0.2402 | 0.0407 | 0.9676 |
| XENOBIOTIC_METABOLISM | BCL2 | 0.1919 | 0.0511 | 0.0949 | 1.5255 | 0.1303 | -0.8388 | 0.4036 | 1.9627 | 0.0525 |
| FATTY_ACID_METABOLISM | BCL2 | -0.3020 | 0.0019 | 0.0106 | -1.7528 | 0.0827 | -1.4223 | 0.1581 | 0.4395 | 0.6613 |
| OXIDATIVE_PHOSPHORYLATION | BCL2 | -0.2846 | 0.0035 | 0.0160 | -2.1654 | 0.0327 | -1.8502 | 0.0672 | -0.6938 | 0.4894 |

| | | | | | | | | | | |
|---|---|---|---|---|---|---|---|---|---|---|
| GLYCOLYSIS | BCL2 | -0.2402 | 0.0142 | 0.0431 | -1.7808 | 0.0780 | -1.3183 | 0.1904 | 1.1892 | 0.2372 |
| REACTIVE_OXYGEN_SPECIES_PATHWAY | BCL2 | 0.0172 | 0.8626 | 0.8804 | 0.3097 | 0.7574 | -2.1591 | 0.0332 | -0.4487 | 0.6546 |
| P53_PATHWAY | BCL2 | 0.2112 | 0.0315 | 0.0701 | 1.4158 | 0.1600 | 0.3336 | 0.7394 | -0.7039 | 0.4831 |
| UV_RESPONSE_UP | BCL2 | 0.0242 | 0.8069 | 0.8584 | 0.3448 | 0.7310 | -0.0665 | 0.9471 | 0.7405 | 0.4608 |
| UV_RESPONSE_DN | BCL2 | 0.2101 | 0.0325 | 0.0702 | 1.3334 | 0.1854 | 0.1068 | 0.9152 | 0.1963 | 0.8448 |
| ANGIOGENESIS | BCL2 | 0.3293 | 0.0007 | 0.0062 | 2.2817 | 0.0246 | 0.1015 | 0.9194 | 0.0969 | 0.9230 |
| HEME_METABOLISM | BCL2 | 0.0204 | 0.8372 | 0.8745 | 0.5284 | 0.5984 | -2.3487 | 0.0208 | -1.0445 | 0.2988 |
| COAGULATION | BCL2 | 0.2875 | 0.0032 | 0.0158 | 2.5858 | 0.0112 | -0.2727 | 0.7856 | 1.5145 | 0.1331 |
| IL2_STAT5_SIGNALING | BCL2 | 0.1767 | 0.0728 | 0.1255 | 0.6780 | 0.4994 | -0.6430 | 0.5217 | 0.6964 | 0.4878 |
| BILE_ACID_METABOLISM | BCL2 | -0.1341 | 0.1746 | 0.2531 | 0.4758 | 0.6353 | -0.3667 | 0.7147 | 0.6236 | 0.5343 |
| PEROXISOME | BCL2 | -0.1239 | 0.2100 | 0.2957 | -0.3833 | 0.7023 | -0.8449 | 0.4002 | 1.0119 | 0.3140 |
| ALLOGRAFT_REJECTION | BCL2 | 0.2672 | 0.0062 | 0.0244 | 2.0249 | 0.0455 | -1.1094 | 0.2699 | -0.1460 | 0.8842 |
| SPERMATOGENESIS | BCL2 | -0.1376 | 0.1634 | 0.2404 | -0.5581 | 0.5780 | 1.6315 | 0.1059 | -0.0477 | 0.9620 |
| KRAS_SIGNALING_UP | BCL2 | 0.2437 | 0.0129 | 0.0402 | 1.7883 | 0.0768 | -1.0732 | 0.2858 | 0.1740 | 0.8622 |
| KRAS_SIGNALING_DN | BCL2 | -0.1406 | 0.1544 | 0.2305 | -1.5054 | 0.1354 | 1.0054 | 0.3171 | 1.4964 | 0.1377 |
| PANCREAS_BETA_CELLS | BCL2 | 0.0930 | 0.3471 | 0.4339 | 1.2131 | 0.2280 | -1.1128 | 0.2685 | 2.0800 | 0.0401 |
